# Dental topography and diet in marsupials and comparisons with primates

**DOI:** 10.1101/2025.08.29.673041

**Authors:** Dorien de Vries, Amy Gao, Darin A. Croft, Garrett Brown, Nicholas Burkart, April Connell, Ellie Gahan, Emily Spinks, Robin M.D. Beck

## Abstract

Dental topography is a powerful tool for investigating the relationship between diet and tooth shape in mammals. Here, we test whether dental topographic metrics (DTMs) can accurately predict diet in marsupials based on 81 lower molars from 43 extant species in 12 families (representing six of the seven extant orders), and compare them to DTMs of primates with similar diets. Specific DTMs tested included Dirichlet normal energy (DNE) and variants (ariaDNE, convex DNE, and ariaDNE CV), orientation patch count rotated (OPCR), and Relief Index (RFI). We also investigate the use of the ratio of trigonid to talonid height (TriTaHI) as an additional DTM. Highest dietary classification accuracy using a leave-one-out quadratic discriminant function analysis for marsupials was reached when ariaDNE, RFI, molar size, and TriTaHI of lower second and third molars were combined, resulting in 69.2% dietary accuracy. In contrast, the primate sample of lower second molars reached the highest classification accuracy of 80.4% when only ariaDNE, RFI, and molar size were used. Notably, 3D meshes from epoxy casts showed consistently higher OPCR and ariaDNE CV values than meshes from original specimens. When using a cross-validated approach, using the primate sample to predict marsupial diets and vice versa, only ariaDNE, RFI, and molar size were needed to obtain the maximum classification accuracy (69.7% when classifying primates using the marsupial training set; and 61.8% vice versa). The comparative dataset of this study will be very useful for studies aiming to reconstruct the diets of fossil mammals that lack phylogenetically close extant analogues.

## INTRODUCTION

The molars of most mammals are highly complex (Ungar 2010; Berkovitz and Shellis 2018), and it has long been recognised that this complexity is influenced by functional, developmental, and phylogenetic constraints (Evans and Sanson 2003; Lucas 2004; Davis 2011; Burroughs 2019; Martin et al. 2020; Couzens et al. 2021). Dental topographic metrics (DTMs) attempt to quantify aspects of tooth shape so that different teeth with disparate morphologies can be objectively compared with each other (Evans 2013; Berthaume et al. 2020). Three dimensional DTMs (3D-DTMs) such as relief index (RFI; Boyer 2008), Dirichlet Normal Energy (DNE; Bunn et al. 2011) and its variants (e.g., ariaDNE: Shan et al. 2019; convex DNE: Pampush et al. 2022), and Orientation Patch Count (OPC; Evans et al. 2007) and its variants (e.g., OPCR, Evans and Jernvall 2009), do not require homologous structures (e.g., specific cusps or crests) to be identified on the teeth being compared, and so represent a particularly promising approach for quantifying and comparing tooth shape across distantly-related mammal clades (Evans 2013; Evans and Pineda-Munoz 2018; Berthaume et al. 2020).

Many studies have used 3D-DTMs to investigate the relationship between the shape of teeth (usually molars) and diet in mammals. Most of these have focused on primates (e.g., Boyer 2008; Bunn and Ungar 2009; Bunn et al. 2011; Ledogar et al. 2013; Winchester et al. 2014; Berthaume and Schroer 2017; Fulwood et al. 2021a; b; Avià et al. 2022; de Vries et al. 2024a; Selig et al. 2024), but some have looked at other mammalian clades, including carnivorans (Selig 2023; Waldman et al. 2023), bats (Gutzwiller and Hunter 2015; López-Aguirre et al. 2022a; b; Villalobos-Chaves and Santana 2022), treeshrews (Selig et al. 2019), horses (Evans and Janis 2014), pigs (Rannikko et al. 2020), rodents (Spradley 2017), and marsupials (Spradley and Kay 2016; Spradley 2017; Spradley and Phillips 2018; Brannick et al. 2023). These studies have found that 3D-DTMs are typically relatively accurate tools for discriminating among species with different diets within clades, particularly when multiple metrics are used in combination, with dietary prediction accuracy often > 80%.

There have, however, been far fewer studies using 3D-DTMs to compare tooth shape across a phylogenetically broad swathe of mammalian diversity. Those that have done so have found that the accuracy of 3D-DTMs for predicting diet is often much less when comparing members of multiple (often distantly-related) clades at the same time. For example, Winchester et al. (2014) found that dietary prediction accuracy was consistently lower when using a combined sample of lower second molars (m2s) from two different primate groups (platyrrhines and “prosimians” [= strepsirrhines and tarsiers]) than when considering these groups separately, although this accuracy varied depending on the precise combination of metrics used (see Winchester et al. 2014: table 7). Pineda-Munoz et al. (2017) compared members of the marsupial order Diprotodontia with members of the placental orders Carnivora, Primates and Rodentia and found that 3D-DTMs correctly predicted diet 67% of the time (increasing to 82% when Rodentia was excluded); however, several metrics in that study were calculated from entire postcanine tooth rows rather than (as in most other studies) single molars. Most recently, Brannick et al. (2023) found that a combination of DNE, RFI and OPCR of individual upper molars successfully discriminated diet in only 33 out of 56 extant mammals (i.e., 58.9%) from three marsupial and five placental orders.

The reduced accuracy of 3D-DTMs for dietary discrimination when comparing phylogenetically diverse mammals is important because 3D-DTMs are increasingly used to predict diet in extinct species, including members of clades that lack close extant relatives (Harper et al. 2019; White et al. 2021; Brannick et al. 2023), and to test hypotheses of competition between clades (Prufrock et al. 2016a; b; Christison et al. 2022). Both of these types of studies assume that taxa with similarly-shaped teeth (as quantified by 3D-DTMs) are likely to have (or have had) similar diets, and so their conclusions may be questionable if this general relationship does not apply when comparing phylogenetically distant clades. Here we investigate this issue in detail by using 3D-DTMs to compare tooth shape of members of two phylogenetically distantly-related mammalian clades that nevertheless include some species with similar diets, namely marsupials and primates (the latter including platyrrhines, strepsirrhines, and tarsiers).

As members of Metatheria and Eutheria respectively, marsupials and primates last shared a common ancestor at least 125 million years ago (Bi et al. 2018; Beck 2022); this common ancestor almost certainly weighed less than 1kg and had a generalised tribosphenic dentition suited for a primarily insectivorous diet (O’Leary et al. 2013). Recent studies suggest that the primate crown clade (= Euprimates) probably began to diversify in the Palaeocene (Vanderpool et al. 2020; de Vries and Beck 2023; Kuderna et al. 2023); the age of crown clade Marsupialia is more uncertain due to a poor fossil record, with any age from the Late Cretaceous to the Palaeocene-Eocene boundary congruent with current evidence (Beck 2022), but is probably within 10-15 million years of the age of Euprimates, and so the two clades are likely of broadly similar age.

Today, many marsupials and some primates are largely insectivorous, but both clades are also characterised by species that have independently evolved more frugivorous and more folivorous diets, as well as species in which exudates make up a major part of the diet (Amador and Giannini 2021; Dickman and Calver 2022; Lessa et al. 2022; Grabowski et al. 2023; Lintulaakso et al. 2023; Machado et al. 2023). However, other diets are unique to one of the two clades: unlike primates, the diets of some extant marsupials (e.g., larger didelphids and dasyurids) include a large proportion of vertebrate prey (Amador and Giannini 2021; Voss and Jansa 2021; Dickman and Calver 2022; Lessa et al. 2022), and several modern primates are specialised hard-object feeders (Kay et al. 2013; Winchester et al. 2014; Avià et al. 2022; de Vries et al. 2024a), whereas no modern marsupial is.

Similarities in diet (as well as other aspects of their ecology) between marsupials and primates have been the subject of several studies, particularly in the context of understanding the origin of distinctive primate adaptations (Rasmussen 1990; Rasmussen and Sussman 2007; Sussman et al. 2013; Spradley and Phillips 2018; St Clair et al. 2018; Stroik and Schwartz 2018). Further investigation of this issue is also of relevance given the possibility that some marsupial and primate species may compete trophically where they co-occur, specifically in the Neotropics (Stroik 2014; Voss and Jansa 2021), and that some fossil South American metatherians have been described as “primate-like” (Marshall 1982; Goin 2006). However, the extent to which dietary similarities between marsupials and primates are associated with similar dental morphologies has not been studied in detail using 3D-DTMs (but see Christensen 2014, who shows similar associations between tooth morphology and diet across extant marsupial and placental mammals; and Stroik 2014 for a 3D geometric morphometric study of an extant mammalian guild including marsupials and primates). Pineda-Munoz et al. (2016) compared marsupials with primates (and also carnivorans and rodents), but their marsupial sample comprised only diprotodontians (28 species in eight families), and their primate sample was also relatively restricted (three platyrrhines, three strepsirrhines, four cercopithecoids, and one hominoid). In addition, Pineda-Munoz et al. (2016) examined whole lower postcanine tooth rows, whereas most available comparative datasets comprise single teeth only (e.g., Winchester et al. 2014; Brannick et al. 2023; de Vries et al. 2024a; Selig et al. 2024). Brannick et al. (2023), meanwhile, included 14 marsupials (ten didelphids, three dasyurids, and one peramelemorphian) but only one primate (the strepsirrhine *Cheirogaleus medius*) in their comparative set of 30 extant mammals (which also included chiropterans, carnivorans, eulipotyphlans, a rodent, and an artiodactyl).

In this study, we investigate the accuracy of 3D-DTMs for inferring diet from lower molars of a comparative set of 43 extant marsupial species, comprising representatives of six of the seven currently recognised orders (only the notoryctemorphian marsupial moles are not represented), and 12 out of the 21 currently recognised families (see Beck et al. 2022: table 1). We investigate what combination of 3D-DTMs result in greatest accuracy of dietary prediction, and whether incorporation of a simple 2D metric (ratio of trigonid to talonid height) results in improved accuracy, specifically better discrimination between faunivores and folivores. We also test whether the second (m2) or third (m3) lower molar of marsupials (likely homologous with the m1 and m2 of primates, respectively; Averianov et al. 2010; O’Leary et al. 2013; Williamson et al. 2014) results in more accurate dietary predictions.

**Table 1.** Marsupial sample, all *n* = 1 unless specified otherwise. All specimens are original specimens (i.e., not casts). Taxa are listed alphabetically by diet, then family, and then species.

| Diet | Family | Species | Abbreviation | m2 | m3 | Scan type |
| --- | --- | --- | --- | --- | --- | --- |
| **Faunivore** | Dasyuridae | *Dasyercus blythi* | Das bl | x | x | μCT |
|  | Dasyuridae | *Dasyurus hallucatus* | Das ha |  | x | Surface (structured light) |
|  | Dasyuridae | *Dasyurus maculatus* | Das ma | x | x | Surface (structured light) |
|  | Didelphidae | *Chironectes minimus* | Chi mi | x | x | μCT |
|  | Didelphidae | *Didelphis virginiana* | Did vi | x | x | μCT |
|  | Didelphidae | *Philander opossum* | Phi op | x | x | μCT |
| **Folivore** | Macropodidae | *Dendrolagus lumholtzi* | Den lu | x | x | μCT |
|  | Macropodidae | *Dorcopsis muelleri* | Dor mu | x |  | Surface (structured light) |
|  | Macropodidae | *Macropus eugenii* | Mac eu | x | x | μCT |
|  | Macropodidae | *Onychogalea frenata* | Ony fr | x |  | μCT |
|  | Macropodidae | *Petrogale penicillata* | Pet pe | x | x | μCT |
|  | Macropodidae | *Thylogale billardieri* | Thy bi | x | x | Surface (structured light) |
|  | Macropodidae | *Thylogale stigmatica* | Thy st |  | x | μCT |
|  | Phalangeridae | *Phalanger carmelitae* | Pha ca | x |  | μCT |
|  | Phalangeridae | *Phalanger orientalis* | Pha or | x | x | μCT |
|  | Phalangeridae | *Spilocuscus maculatus* | Spi ma | x | x | μCT |
|  | Phalangeridae | *Trichosurus arnhemensis* | Tri ar | x | x | Surface (structured light) |
|  | Phalangeridae | *Trichosurus caninus* | Tri ca | x | x | Surface (structured light) |
|  | Phalangeridae | *Trichosurus vulpecula* | Tri vu | x | x | μCT |
|  | Phascolarctidae | *Phascolarctos cinereus* | Pha ci | x | x | μCT |
|  | Pseudocheiridae | *Petauroides volans* | Pet vo | x | x | μCT |
|  | Pseudocheiridae | *Petropseudes dahli* | Pet da | x | x | μCT |
|  | Pseudocheiridae | *Pseudochirulus forbesi* | Pse fo | x | x | μCT |
|  | Potoroidae | *Aepyprymnus rufescens* | Aep ru | x | x | μCT |
| **Frugivore** | Hypsiprymnodontidae | *Hypsiprymnodon moschatus* | Hyp mo | x | x | μCT |
| **Frugivore-insectivore** | Didelphidae | *Caluromys philander* | Cal ph | x | x | μCT |
|  | Didelphidae | *Caluromys* sp. | Cal sp | x |  | μCT |
|  | Didelphidae | *Caluromysiops irrupta* | Cal ir | x | x | μCT |
|  | Didelphidae | *Marmosa murina* | Mar mu | x | x | μCT |
|  | Microbiotheriidae | *Dromiciops gliroides* | Dro gl | x | x | μCT |
| **Insectivore** | Caenolestidae | *Caenolestes fuliginosus* | Cae fu | x | x | μCT |
|  | Caenolestidae | *Lestoros inca* | Les in | x | x | μCT |
|  | Caenolestidae | *Rhyncholestes raphanurus* | Rhy ra | x | x | μCT |
|  | Dasyuridae | *Antechinus flavipes* | Ant fl | x | x | μCT |
|  | Dasyuridae | *Phascogale tapoatafa* | Pha ta | x | x | μCT |
|  | Dasyuridae | *Sminthopsis crassicaudata* | Smi cr | x | x | μCT |
|  | Dasyuridae | *Murexia longicaudata* | Mur lo | x | x | μCT |
|  | Didelphidae | *Monodelphis domestica* | Mon do | x | x | μCT |
|  | Petauridae | *Dactylopsila trivirgata* (n=2) | Dac tr | x | x | μCT |
|  | Petauridae | *Petaurus australis* | Pet au | x | x | μCT |
| **Omnivore** | Peramelidae | *Isoodon obesulus* | Iso ob | x | x | μCT |
|  | Peramelidae | *Perameles gunnii* | Per gu | x | x | Surface (structured light) |
|  | Peramelidae | *Perameles nasuta* | Per na |  | x | Surface (structured light) |

We combine our marsupial sample with a primate sample comprising 52 species (32 platyrrhines, 18 strepsirrhines, 2 tarsiers) to compare 3D-DTM values of marsupials and primates with similar diets. We test the accuracy of 3D-DTMs for dietary prediction when applied to this combined marsupial and primate sample using a leave-one-out approach. Finally, we investigate the accuracy of 3D-DTMs for predicting diet using a cross-validation approach classifying the marsupial sample based on a primate training set, and vice versa, to test whether comparative dental topographic data from one clade can be used to accurately predict diets of members of another, phylogenetically distant clade. The results of our study may be of particular relevance to future studies that attempt to infer the diets of fossil mammals that may have been ecologically similar (but not particularly closely related) to modern marsupials and primates.

## MATERIALS AND METHODS

### Study sample

Marsupial specimens were downloaded from Morphosource (Boyer et al. 2016), as either raw μCT scan data (.tiff or .dcm image stacks) or 3D surface meshes (.ply files) that had already been generated from μCT or surface scan data (see Online Resource 1 for a full list of specimens and their raw dental topographic data). Following the freeware protocol of de Vries et al. (2024a: online resource 2), the μCT scan data were processed in Slicer (v.5.0.3, Kikinis et al. 2014) using the SlicerMorph module (v.e8d4a2e, Rolfe et al. 2021) to create a surface mesh. The 3D surface meshes were opened in MeshLab (v.2022.02, Cignoni et al. 2008) to crop individual molars for further processing. The plesiomorphic dental formula of marsupials is four molars, which contrasts with three molars in placentals; the marsupial m1 is likely a molariform deciduous premolar that is retained into adulthood (Averianov et al. 2010; O’Leary et al. 2013; Williamson et al. 2014). If this hypothesis is correct, then the marsupial m2 is homologous with the m1 of primates and other placentals, and the marsupial m3 is homologous with the placental m2; based on these homologies, the most appropriate marsupial tooth to compare with the m2 of placentals is arguably m3 rather than m2. For this reason, both m2 and m3 were isolated as separate files for further processing.

The “prosimian” primate (= strepsirrhine and tarsier) specimens were downloaded from Morphosource as 3D surface meshes of epoxy casts of individual m2s created for previous studies (Boyer 2008; Bunn et al. 2011). Because we use the freeware processing pipeline of de Vries et al. (2024a) here (rather than the one followed by Boyer 2008; and Winchester et al. 2014), we used the cropped but unsmoothed (“raw”) versions of these files as our starting point (see Online Resource 1 for a full list of specimens).

For both the marsupial and the “prosimian” sample, each tooth mesh was examined by one us (RMDB) in MeshMixer (v.3.5.474, Autodesk 2017), and any specimens that showed evidence of major damage (e.g., cracks, large chips in the enamel) or anything more than minor wear were excluded from further processing. Following this step, we retained 81 marsupial specimens (41 m2s, 40 m3s) representing 43 species in 12 families (see Table 1), and 30 “prosimian” specimens (all m2s), representing 20 species in eight families (see Online Resource 1). Subsequent processing steps followed the pipeline and scripts of de Vries et al. (2024a: online resource 2), which we briefly summarise here. Firstly, minor damage and other anomalies (e.g., bubbles in scans of casts) in the retained surface meshes were repaired in MeshMixer. Meshlab and bash scripts were then used to remove duplicate faces and isolated pieces, to centre the meshes and orient them into occlusal view, and to downsample the meshes to 10,000 triangles. Several specimens (the m2 and m3 of *Caluromysiops irrupta* FMNH 60698; the m3 of *Onychogale frenata* UMZC A12.59/3) were of too low resolution to yield meshes with at least 10,000 triangles, and could therefore not be downsampled. Their triangle counts were 8,251, 7,039, and 8,874 respectively. As triangle count affects DTMs such as DNE and OPCR, special care was given to these specimens when interpreting results. Finally, the centred, orientated, and simplified meshes were smoothed using the ‘HC Laplacian Smooth’ (HCL) option in Meshlab, as this yielded the highest dietary prediction accuracy for a sample of extant platyrrhines (de Vries et al. 2024a).

The platyrrhine sample comprises 145 m2s from 32 species in five families (Atelidae, Aotidae, Callitrichidae, Cebidae, Pitheciidae), which had already been processed according to the de Vries et al. (2024a) protocol; these fully processed platyrrhine specimens can be downloaded from MorphoSource (project ID 000471738). See Online Resource 1 for a full list of specimens.

### Calculation of dental topographic metrics

The dental topographic metrics relief index (RFI; Boyer 2008), orientation patch count rotated (OPCR; Evans and Jernvall 2009), Dirichlet normal energy (DNE; Bunn et al. 2011), convex DNE (Pampush et al. 2022), and 2D outline area were calculated in R (v4.2.0, R Core Team 2022) and RStudio (v2022.07.2, RStudio Team 2020) using the ‘molaR_Batch’ function of the molaR package (v.5.3, Pampush et al. 2016, 2022) with findAlpha = TRUE, following the recommended settings in the R package manual (v.5.3, Pampush et al. 2025). Relief index (RFI) measures the 3D relief of a surface; specifically, it is calculated as the natural log of the ratio between the square roots of the 2D outline area and 3D surface area of the tooth (Boyer 2008). Orientation patch count rotated (OPCR) measures the number of distinct patches of a surface that face in one of eight directions (Evans et al. 2007; Evans and Jernvall 2009). The natural log of the 2D outline area was taken as a measure of size and will be referred to as ‘lnOA’ hereafter.

Dirichlet normal energy measures the total curvature of a surface (Bunn et al. 2011). We calculated three variants of DNE and compared its dietary signal across the three variants: DNE, convex DNE, and ariaDNE. Whereas DNE and ariaDNE include both the convex and concave curvatures of a tooth, convex DNE only measures the outwardly facing curvature of a tooth, and thus reflects a more straightforward functional signal (i.e., only the cutting surfaces of a tooth; (Pampush et al. 2022). On the other hand, ariaDNE has been shown to be less susceptible to differences in surface mesh generation and processing protocols, triangle counts, and mesh anomalies in triangle distribution and density (Shan et al. 2019). Standard DNE and convex DNE were calculated in R using the molaR package. For calculation of ariaDNE, MATLAB (vR2021b, The MathWorks Inc. 2021) was used, with ε set to 0.08; this is nearer the upper end of the recommended range of ε (0.04 to 0.1), and thus captures mostly larger structures while ignoring smaller features of the crown (Shan et al. 2019). Fulwood et al. (2021b) found that the coefficient of variance (CV) of ariaDNE of a tooth, or ariaDNE CV, is a powerful additional metric for distinguishing diet in strepsirrhine primates. These authors found that folivores had the highest ariaDNE CV values, and insectivores had the lowest values among their comparative sample, whereas previous studies had found that these two that are often hard to distinguish based on DTMs unless a measure of size was included (Fulwood et al. 2021b). We calculated ariaDNE CV in MatLab using code provided by EL Fulwood.

In primates, insectivores typically do not exceed 500 g in body mass (a possible exception is the aye aye, *Daubentonia madagascariensis*, which weighs ∼2kg (Sefczek et al. 2020), but we have classified it as a hard object feeder here; see “Dietary Classification” below), whilst all folivores are > 500 g (Kay’s threshold: Kay 1975; Kay and Hylander 1978). Because of this, combining DTMs with a measure of molar size (e.g., lnOA, as used here) improves discrimination between primate folivore and insectivore categories, which otherwise have overlapping DTM values due to their steeply sloped cusps (Bunn et al. 2011). Most marsupials confirm the general trend of insectivores being smaller than folivores, although there are a few marsupial folivores below the 500 g threshold (e.g., the folivorous pseudocheirid *Pseudochirulus mayeri*, which weighs only 152g, Hogue and ZiaShakeri 2010). However, among marsupials, faunivores that regularly consume vertebrate prey have ariaDNE and lnOA that are non-significantly different to those of folivores (see Results), and, unlike most marsupial insectivores (Amador and Giannini 2021), marsupial faunivores overlap folivores in size.

To aid in successfully discriminating between faunivorous and folivorous marsupials, we propose a simple 2D ratio here: the trigonid-talonid height index (TriTaHI; Fig. 1). The theoretical basis for this ratio is the observation that the trigonid of marsupials and primates is typically markedly taller than the talonid in insectivorous and faunivorous mammals with recognisably tribosphenic lower molars, whereas they are more similar in height in folivores (Zimicz 2012, 2014; St Clair and Boyer 2016; Goin et al. 2020). Versions of this metric have been proposed by several previous authors (Şenyürek 1951; Dewar 2003; Stroik 2014; Goin et al. 2020). A positive TriTaHI value indicates a trigonid taller than the talonid, a value close to zero indicates similar trigonid and talonid heights, and a negative value indicates a taller talonid than trigonid. To calculate TriTaHI, a picture of the 3D tooth mesh was taken in lingual view in MeshLab. In ImageJ (v.13.0.6, Schneider et al. 2012), the picture was used to measure the maximum length of the molar parallel to the plane of the base of the enamel crown. A straight line running from the junction of the crown and the root at the mesial and distal ends of the molar was drawn as a ‘baseline’ to represent this plane (see Fig. 1: baseline). Maximum molar length was measured parallel to this baseline (see Fig. 1: max. length). The maximum heights of the trigonid and talonid were measured perpendicular to the baseline (see Fig. 1) from the tip of the highest cusp in the trigonid or talonid to the baseline. The difference in heights (maximum trigonid height - maximum talonid height) was then divided by the maximum molar length to obtain the relative trigonid-talonid height. We calculated TriTaHI for all marsupial and primate specimens and tested if the inclusion of this 2D metric resulted in improved dietary classification accuracy, particularly when distinguishing faunivores from folivores.

**Fig. 1.**
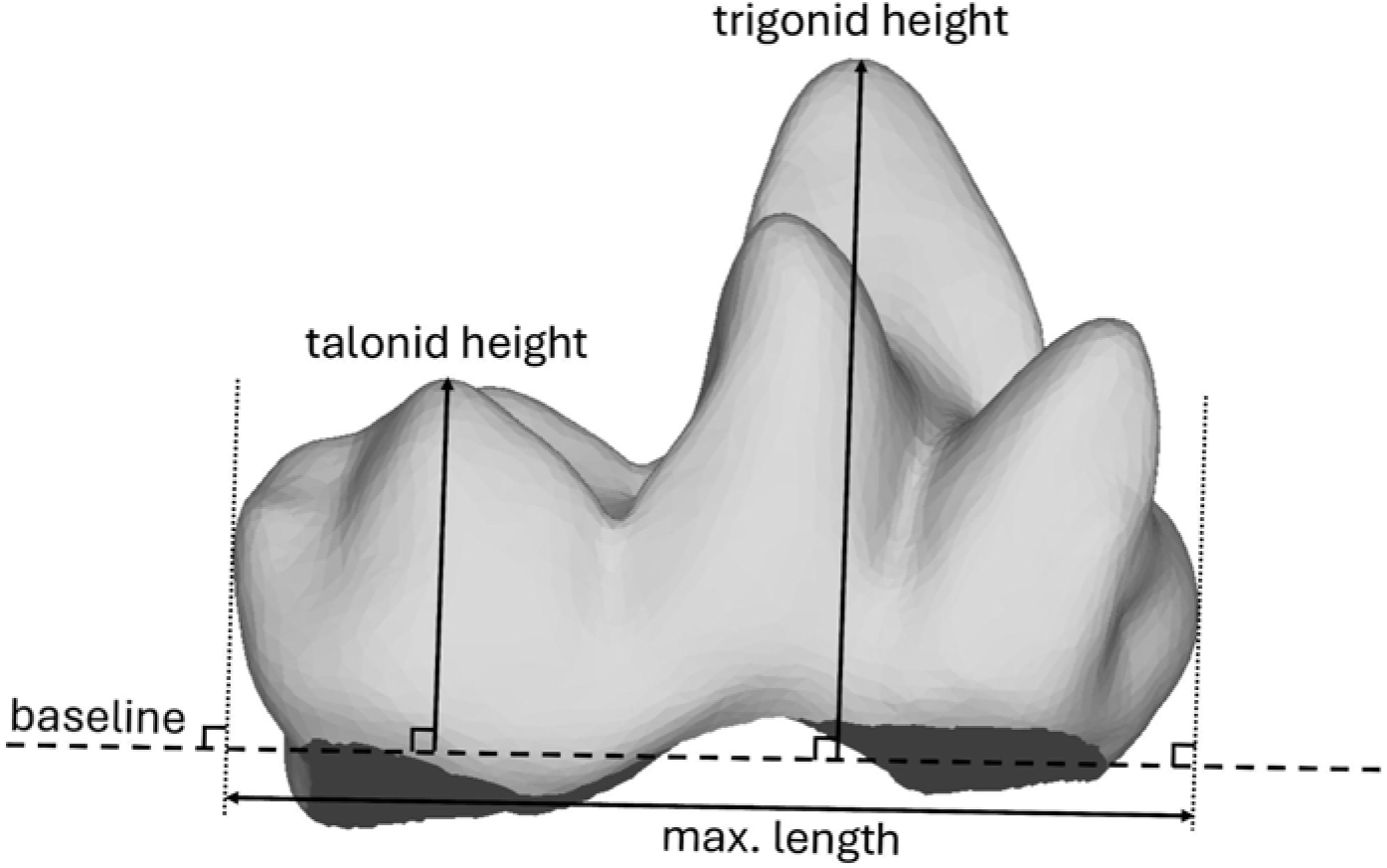
Measurements for calculating the Trigonid-Talonid Relief illustrated on the lingual view of *Chironectes minimus* AMNH 129701 m3, anterior to the right. Abbreviation: **max. length**, maximum length of molar measured parallel to the baseline.

### Dietary classification

There are multiple published dietary classifications available for the marsupial, “prosimian” and platyrrhine species included in our study (see summary in Tables 2 and 3); however, we use our own scheme here (see Online Resource 2 for full details). We largely retained the dietary categories and classification of platyrrhines proposed by de Vries et al. (2024a), which is based on a cluster analysis of quantitative dietary data from 98 primary studies; where possible, we classified our marsupial and “prosimian” species using the relative proportions of food types in different dietary categories calculated for platyrrhines by de Vries et al. (2024a: table 3) as a guide. For example, de Vries et al. (2024a: table 3) found that platyrrhine folivores consume > 50% leaves and < 50% “fruits” (= fruits, flowers, and fungi combined), whereas platyrrhine frugivores consume > 50% “fruits” and < 30% leaves. Based on this, the phalangerid marsupials *Phalanger orientalis*, *Phalanger carmelitae*, *Spilocuscus maculatus*, and *Trichosurus vulpecula* have been classified as folivores here, because they appear to consume a greater proportion of leaves than fruits (Menzies and Pernetta 1986; Evans 1992; How and Hillcox 2000; Salas 2002; Farida 2022) similar to platyrrhine folivores; by contrast, some other studies have classified these phalangerids as frugivores or frugivore-folivores (see Table 2).

**Table 2.** Previous dietary classifications of extant marsupial species in our dataset, and the classification scheme used in the current study. Taxa are listed alphabetically by family and then species.

| Family | Species | Pineda-Munoz et al (2016) | Spradley and Kay (2016)/MammalDIET | Brannick et al. (2023) | Lintulaakso et al. (2023) | This study |
| --- | --- | --- | --- | --- | --- | --- |
| Caenolestidae | *Caenolestes fuliginosus* | n/a | Insectivore | n/a | Vermivore-insectivore | Insectivore |
| Caenolestidae | *Lestoros inca* | n/a | n/a | n/a | n/a | Insectivore |
| Caenolestidae | *Rhyncholestes raphanurus* | n/a | n/a | n/a | Insectivore-omnivore | Insectivore |
| Didelphidae | *Caluromys philander* | n/a | Frugivore<sup>a</sup> | Plant-dominated omnivore<sup>a</sup> | Frugivore-omnivore | Frugivore-insectivore |
| Didelphidae | *Caluromys* sp | n/a | Frugivore<sup>a</sup> | Plant-dominated omnivore<sup>a</sup> | n/a | Frugivore- insectivore |
| Didelphidae | *Cauromysiops irrupta* | n/a | n/a | n/a | Frugivore-nectarivore | Frugivore-insectivore |
| Didelphidae | *Chironectes minimus* | n/a | Insectivore | Carnivore | Insectivore-carnivore | Faunivore |
| Didelphidae | *Didelphis virginiana* | n/a | Frugivore<sup>b</sup> | Animal-dominated omnivore | Insectivore-omnivore | Faunivore |
| Didelphidae | *Marmosa murina* | n/a | Insectivore | n/a | Insectivore-omnivore | Frugivore-insectivore |
| Didelphidae | *Monodelphis domestica* | n/a | Insectivore | Animal-dominated omnivore | Carnivore-scavenger | Insectivore |
| Didelphidae | *Philander opossum* | n/a | n/a | Animal-dominated omnivore | Insectivore-omnivore | Faunivore |
| Petauridae | *Dactylopsila trivirgata* | Insectivore | Frugivore | n/a | Insectivore-myrmecophage | Insectivore |
| Petauridae | *Petaurus australis* | Gumivore | Frugivore | n/a | Nectarivore-palynivore | Insectivore |
| Potoroidae | *Aepyprymnus rufescens* | n/a | Grazer | n/a | Browser-rootivore | Folivore |
| Macropodidae | *Dendrolagus lumholtzi* | Frugivore | Folivore | n/a | Browser | Folivore |
| Macropodidae | *Macropus eugenii* | Herbivore<sup>c</sup> | n/a | n/a | Grazer | Folivore |
| Macropodidae | *Onychogalea frenata* | Herbivore | n/a | n/a | Browser-grazer | Folivore |
| Macropodidae | *Petrogale penicillata* | Herbivore<sup>d</sup> | n/a | n/a | Grazer | Folivore |
| Microbiotheriidae | *Dromiciops gliroides* | n/a | Insectivore | n/a | Vermivore-insectivore | Insectivore-Frugivore |
| Phascolarctidae | *Phascolarctos cinereus* | n/a | Folivore | n/a | Browser | Folivore |
| Pseudocheiridae | *Petauroides volans* | Herbivore | Folivore | n/a | Browser | Folivore |
| Pseudocheiridae | *Petropseudes dahli* | n/a | Folivore | n/a | Florivore-frugivore | Folivore |
| Pseudocheiridae | *Pseudochirulus forbesi* | n/a | n/a | n/a | Browser-frugivore | Folivore |
| Phalangeridae | *Phalanger orientalis* | n/a | Frugivore<sup>e</sup> | n/a | Browser-frugivore | Folivore |
| Phalangeridae | *Spilocuscus maculatus* | n/a | n/a | n/a | Browser-frugivore | Folivore |
| Phalangeridae | *Trichosurus vulpecula* | Herbivore | Frugivore | n/a | Frugivore-browser | Folivore |

<sup>a</sup> Based on *Caluromys derbianus*
<sup>b</sup> Based on *Didelphis albiventris*
<sup>c</sup> Based on seven other *Macropus* species
<sup>d</sup> Based on three other *Petrogale* species
<sup>e</sup> Based on *Phalanger gymnotis*

**Table 3.** Previous dietary classifications of extant marsupial species in our dataset, and the classification scheme used in the current study. Taxa are listed alphabetically by family and then species.

| Family | Species | Boyer (2008) | Bunn et al. (2011) | Winchester et al. (2014) | Lintulaakso et al. (2023) | This study |
| --- | --- | --- | --- | --- | --- | --- |
| Cheirogaleidae | *Microcebus griseorufus* | Omnivore | Omnivore | Omnivore | Insectivore-omnivore | Frugivore-insectivore |
| Cheirogaleidae | *Phaner furcifer* | Omnivore | Omnivore | Omnivore | Gumivore-omnivore | Insectivore |
| Daubentoniidae | *Daubentonia madagascariensis* | Frugivore-granivore | Frugivore | Hard object feeder (Frugivore in supplementary, but HOF in main text) | Vermivore-granivore | Hard-object feeder |
| Galagidae | *Galagoides demidoff* | Insectivore | Insectivore | Insectivore | Insectivore-myrmecophage | Insectivore |
| Galagidae | *Galago senegalensis* | n/a | Insectivore | Insectivore | Gumivore-insectivore | Insectivore |
| Indriidae | *Avahi laniger* | Folivore | Folivore | Folivore | Browser | Folivore |
| Indriidae | *Indri indri* | Folivore | Folivore | Folivore | Browser | Folivore |
| Indriidae | *Propithecus deckenii* | Folivore or omnivore | Folivore | Folivore | Browser-florivore | Folivore |
| Lemuridae | *Eulemur fulvus* | Folivore or omnivore | Omnivore | Omnivore | Browser-frugivore | Folivore |
| Lemuridae | *Hapalemur griseus* | n/a | Folivore | Folivore | Grazer-browser | Folivore |
| Lemuridae | *Lemur catta* | Folivore or omnivore | Omnivore | Omnivore | Browser-frugivore | Folivore |
| Lepilemuridae | *Lepilemur edwardsi* | Folivore | Folivore | Folivore | Browser-frugivore | Folivore |
| Lepilemuridae | *Lepilemur leucopus* | Folivore | Folivore | Folivore | Browser | Folivore |
| Lepilemuridae | *Lepilemur ruficaudatus* | Folivore | Folivore | Folivore | Frugivore-browser | Folivore |
| Lorisidae | *Arctocebus calabarensis* | Insectivore | Insectivore | Insectivore | Vermivore-insectivore | Insectivore |
| Lorisidae | *Loris tardigradus* | Insectivore | Insectivore | Insectivore | Insectivore-omnivore | Insectivore |
| Lorisidae | *Nycticebus javanicus* | Omnivore | Unknown | Omnivore | Frugivore-insectivore (based on *Nycticebus coucang*) | Omnivore |
| Lorisidae | *Perodicticus potto* | Frugivore-gramnivore | Frugivore | Frugivore | Gumivore-omnivore | Frugivore-insectivore |
| Tarsiidae | *Tarsius pelengensis* | Insectivore | Insectivore | Insectivore | Insectivore | Insectivore |

We added three additional categories for diets present in our marsupial and/or “prosimian” samples that are not observed in platyrrhines: insectivory, in which invertebrates form the majority of the diet, with much less fruit consumption than in the frugivore-insectivore category of de Vries et al. (2024a); faunivory, in which there is regular consumption of vertebrate prey; and omnivory, in which animals, fruit, and non-reproductive parts of plants (e.g., leaves, roots) all form major parts of the diet - this differs from frugivory-insectivory, in which non-reproductive parts of plants are rarely eaten (see de Vries et al. 2024a: table S2).

We also made the following two changes to the categories of de Vries et al. (2024a), to ensure that they reflected the mechanical properties of their dietary components (see Table 4). Firstly, the ‘seed eating’ category was renamed to ‘hard-object feeder’, and *Cebus* and *Sapajus* were added to this category (following Winchester et al. 2014) based on the habitual intake of mechanically challenging plant tissues such as palm nuts and the bases of palm leaves (Taylor and Vinyard 2009), albeit at a lower frequency than other hard-object feeding primates like *Cacajao* and *Chiropotes* (de Vries et al. 2024a: table 3). We also assigned the strepsirrhine *Daubentonia madagascariensis* to the hard-object feeder category. Although *Daubentonia* is well known for feeding on invertebrates that it extracts from wood using enlarged incisors and its specialised third digit (e.g., Erickson 1995; Sefczek et al. 2020), field studies show that *Canarium* seeds (which are extremely hard-shelled; Sterling 1994) can form > 25% of its diet (Randimbiharinirina et al. 2018), and are the most consumed food item during certain times of the year (Sefczek 2018); by comparison, the diet of the platyrrhine *Pithecia*, which has been consistently classified as a seed-eater or hard-object feeder in previous studies (Ledogar et al. 2013; Winchester et al. 2014; de Vries et al. 2024a), consumes on average only 21% seeds (de Vries et al., 2024a: table S2).

**Table 4.**
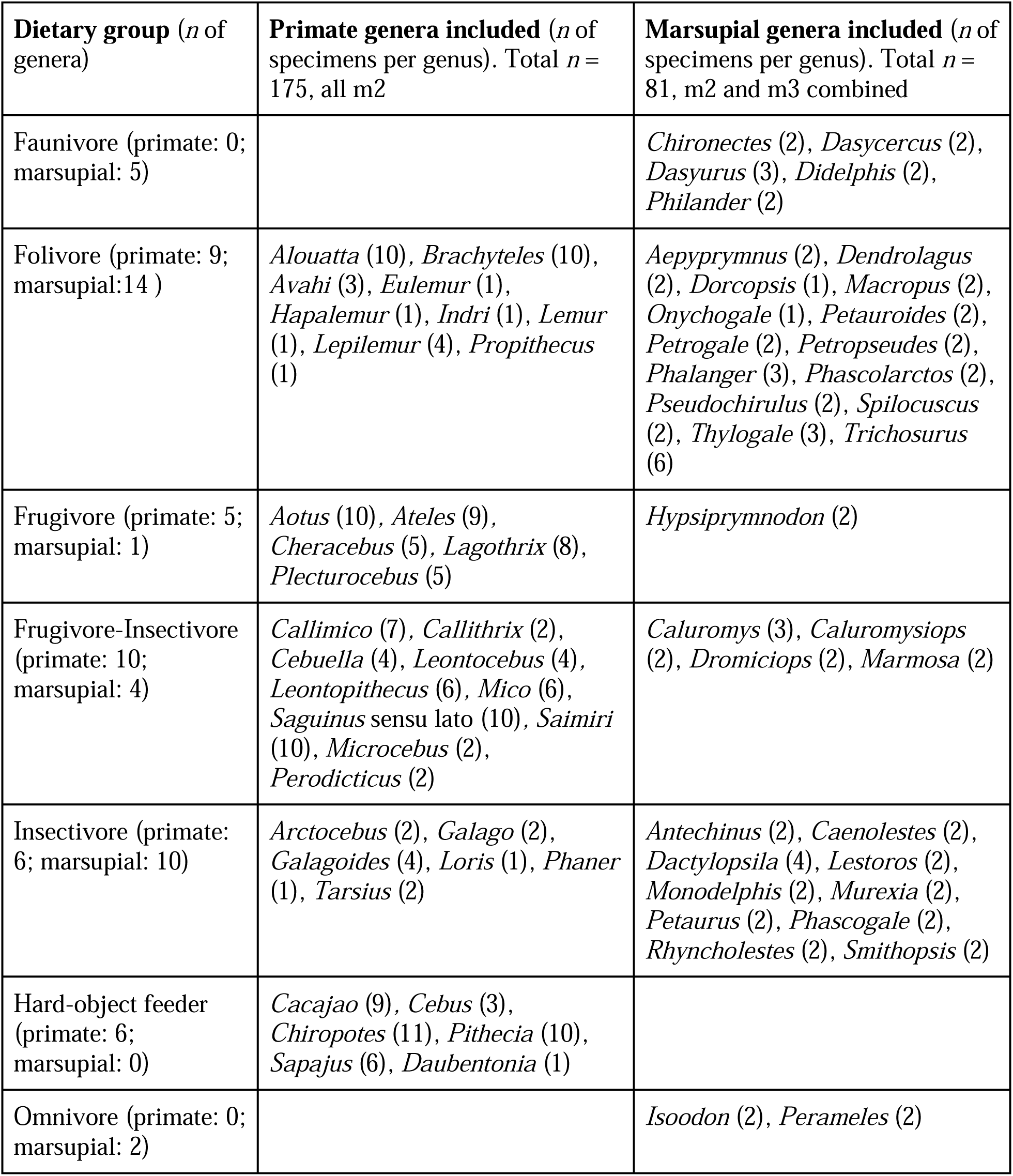
Total sample breakdown of number of genera per dietary category.

Secondly, species that primarily feed on exudates were not grouped into a separate exudate-feeding category here (contra de Vries et al., 2024a and following Selig et al. 2024), but were instead classified based on the proportions of their remaining (non-exudate) dietary items that are mechanically demanding (note that Boyer 2008; Bunn et al. 2011; and de Vries et al. 2024b likewise did not employ an exudate-feeding category for their “prosimian” sample). This decision was based on the principle that exudates are liquids that require no mastication (Nash 1986; Selig et al. 2024) and thus place no mechanical constraints on molars. Thus, the platyrrhines *Callithrix penicillata*, *Cebuella pygmaea*, and *Mico argentata* and the strepsirhine *Microcebus griseorufus* were classified as frugivore-insectivores because fruits and invertebrates both form major parts of their non-exudate diets (Crowley et al. 2014; de Vries et al. 2024a: table S2); by contrast, the strepsirrhines *Galago senegalensis* and *Phaner furcifer* and the marsupial *Petaurus australis* were classified as insectivores, as invertebrates form the majority of their non-exudate diets, with relatively little consumption of fruit (Smith and Russell 1982; Harcourt 1986; Schülke 2003 fig. 2). A full justification of the dietary scheme used here is given in Online Resource 2. See Fig. 2 for the range of included marsupial and primate dietary categories and an example of a molar surface mesh.

**Fig. 2.**
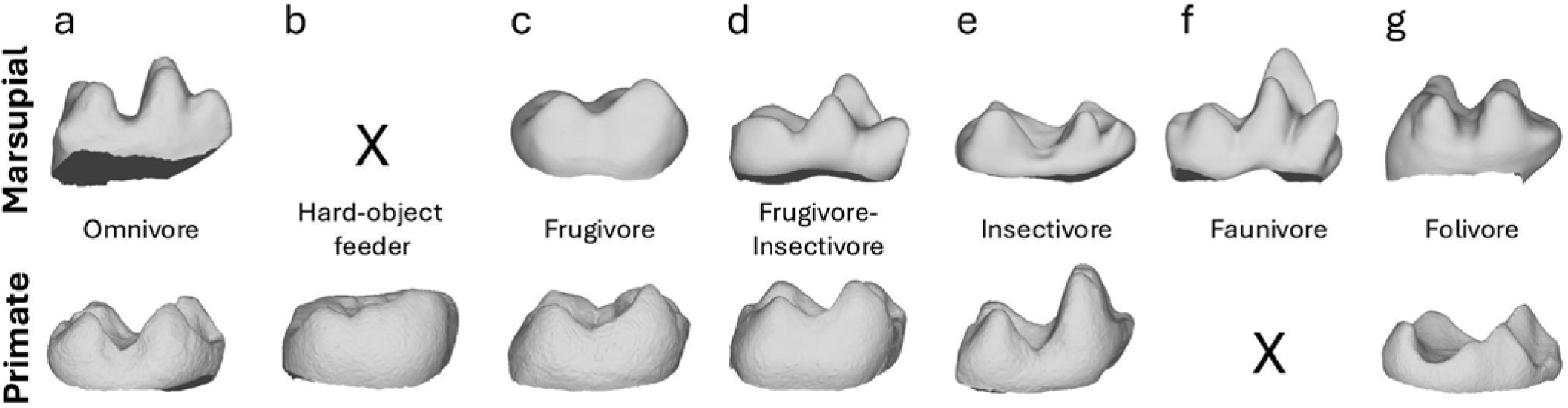
Examples of lower molar shape of marsupial (m2 or m3; above) and primate (m2, below) per dietary category. All images are in lingual view with anterior to the right, but not to scale. **a**. Omnivore: *Perameles nasuta* MAGNT U7608 m3 (above) and *Nycticebus javanicus* AMNH 101508 m2 (below). **b**. Hard-object feeder: *Cacajao calvus* AMNH 73720 m2 (below). **c**. Frugivore: *Hypsiprymnodon moschatus* AMNH 184580 m3 (above) and *Aotus nigriceps* AMNH 75996 m2 (below). **d**. Frugivore-Insectivore: *Caluromys* sp. DU-EA-162 m2 (above) and *Saimiri boliviensis* AMNH 38792 m2 (below). **e**. Insectivore: *Lestoros inca* MVZ 116049 m2 (above) and *Galago senegalensis* AMNH 187362 m2 (below). **f**. Faunivore: *Chironectes minimus* AMNH 129701 m3 (below); **g**. Folivore: *Dendrolagus lumholtzi* AMNH 65254 m2 (above) and *Lemur catta* AMNH 170741 m2 (below).

### Statistical analyses

To test whether DTMs differ significantly between the m2 and m3 in marsupials, paired t-tests or Wilcoxon signed-rank tests (depending on whether the data were normally distributed or not) were used to compare the m2 and m3 of each marsupial specimen for which both of these teeth were available for comparison (thus excluding seven specimens that are represented by only an m2 or an m3). First, normality of data was tested for each group using the ‘shapiro.test’ function part of the R base package (R v4.2.0, R Core Team 2022). When data of both groups were normally distributed, paired t-tests were run for each DTM and dietary category comparing the m2 to the m3 of the same specimen. Paired t-tests were run using the ‘t.test’ function of the stats R package, which is part of the R base package, setting ‘paired’ to ‘TRUE’. When data of a group were not normally distributed, a Wilcoxon signed-rank test was run using the ‘wilcox.test’ with ‘paired = TRUE’ from the stats R package. The frugivore category was excluded from these tests, as its sample size was only a single pair (one m2 and one m3).

To check whether DTMs differ significantly between different dietary categories within marsupials, we ran Kruskal-Wallis or one-way ANOVA tests for each metric. Residuals of the ANOVA were tested for normality using the ‘shapiro.test’ function as described above. If residuals were normally distributed, a one-way ANOVA test was run using the ‘aov’ function of the stats R package. If residuals were non-normally distributed, a Kruskal-Wallis test was run using the ‘kruskal.test’ function of the stats R package. If the one-way ANOVA or Kruskal-Wallis test returned a significant *p*-value (*p* > 0.05), a post-hoc test was run to identify which dietary categories differed significantly for that metric: a Tukey HSD following the ANOVA tests using the ‘TukeyHSD’ function of the stats R package, or a Dunn test following the Kruskal-Wallis tests using the ‘dunn.test’ function of the dunn.test R package (v.1.3.5, Dinno 2017). We ran these tests on specimen values, both for the marsupial m2 and m3 data separately, as well as for a combined (m2 + m3) marsupial sample.

When comparing marsupial DTMs to those of primates of the same dietary category, a Welch t-test or a Wilcoxon rank-sum test was run on specimen values. Marsupial m2 and m3 data were combined for the marsupial sample, and platyrrhine and strepsirrhine data were combined for the primate sample. These tests were only run on dietary categories that had sample sizes > 30 specimens for marsupials and primates combined, namely frugivore-insectivores (n = 62), insectivores (n = 34), and folivores (n = 64). Again, the normality of each group was first tested using the ‘shapiro.test’ function. The Welch t-test was applied to normally distributed data by using the same t-test function but setting ‘paired’ and ‘var.equal’ to ‘FALSE’. For the non-normally distributed data, the Wilcoxon rank-sum tests were run using the same ‘wilcox.test’ function as above, and again setting paired to ‘FALSE’.

To test the dietary classification accuracy of various sets of DTMs for a marsupial and primate sample, a quadratic discriminant analysis (QDA) was run using the ‘qda’ function of the MASS R package (v.7.3.56, Ripley et al. 2013) on species-mean data. All DTMs were tested for homogeneity of variance by running a Levene’s test for the combined marsupial and primate sample per dietary category using the ‘leveneTest’ function of the car R package (v3.0.5, Fox et al. 2007). As variance was not equal for every DTM, a QDA was used as (unlike a linear discriminant analysis) a QDA does not assume equal variance of groups and is therefore more appropriate for our data (James et al. 2013). The prior probabilities of dietary group membership were set equal for all dietary categories rather than using the class proportions of the training set. To run the QDA using a leave-one-out approach, the ‘CV=TRUE’ command was used. Additionally, we tested the classification accuracy of the marsupial and primate sample using a cross-validated QDA. We did this by setting the training set as marsupial-only, to then test the classification accuracy of the model using the primate sample as the test sample, and vice versa. Dietary classification accuracy was calculated based on the test accuracy of the dietary categories that were present in the training set only, thus excluding the test results of the dietary categories not present in the training set.

QDAs were run on species-mean data for the following two reasons: 1) as the marsupial m2 and m3 data most often came from the same specimen and, as for most DTMs, there was no significant difference in m2 and m3 shape (see Results), QDA results using the leave-one-out approach would have artificially high accuracy for example, when the m2 was left out but the m3 of the same specimen was retained in the training set; and 2) when running QDAs on the combined marsupials and primates dataset, species means provided a more balanced sample of species per dietary category across marsupials and primates (average of 7.2 marsupial species per diet and 8.5 primate species per diet) compared to the relatively unbalanced sample of specimens per diet (i.e., higher for primates, with an average of 29 primate specimens per dietary category versus an average of only 14 marsupial specimens per diet), thus ensuring primates and marsupials were equally represented across our samples in the QDAs. See Online Resource 3 for R-code and input files used in this study.

### Institutional abbreviations

AMNH, American Museum of Natural History; DU-EA, Duke Evolutionary Anthropology Department; FMNH, Field Museum of Natural History; MAGNT, Northern Territory Museum and Art Gallery; MVZ, Museum of Vertebrate Zoology; UMZC, University Museum of Zoology Cambridge.

## RESULTS

### Marsupial dental topography

#### Second versus third molar

Paired t-tests and Wilcoxon signed-rank tests reveal that few DTMs differ significantly between the m2 and m3 of marsupials (*p* > 0.05; see Online Resource 2: Table S1 for *p*-values of all pairwise comparisons). The only variables that differ significantly are ariaDNE and TriTaHI in folivores (*p* = 0.014 and *p* = 0.017, respectively, both with greater values in m3 than in m2) and lnOA in insectivores and faunivores (*p* = 0.041 and *p* = 0.007, respectively, with larger m2s in insectivores and larger m3s in faunivores). The t-values, degrees of freedom, and *p*-values of all paired t-tests, and the V and *p*-values of the Wilcoxon signed-rank tests are reported in the Online Resource 2. Since only four of the 30 m2-m3 pairs differed significantly, we group the m2-m3 data for analyses that combine or compare the marsupial and primate samples, as this increases our sample sizes and allows us to run QDAs with more variables.

#### Molar shape per dietary category

For all ANOVA and Kruskal-Wallis results reported below, we exclude the frugivore category as its sample size was only two specimens (one m2, one m3) from a single specimen of *Hypsiprymnodon moschatus*. We do not find significant differences in ariaDNE, OPCR, RFI or ariaDNE CV between diets for the m2-only and m3-only marsupial samples. However, lnOA is significantly greater in faunivores and folivores than in insectivores and frugivore-insectivores, for both m2 and m3. Values of the trigonid-talonid height index (TriTaHI) are significantly higher in faunivores, insectivores, and frugivore-insectivores than in folivores for m2. For m3, only faunivores and frugivore-insectivores show significantly higher TriTaHI values than folivores.

When considering the combined m2 and m3 sample, which doubles the sample sizes for most dietary categories, we find more significant differences between DTMs: ariaDNE is significantly greater in faunivores, insectivores, and frugivore-insectivores than in frugivores; RFI is significantly greater in faunivores than in insectivores and folivores, and also significantly greater in omnivores than in folivores; lnOA is significantly greater in faunivores and folivores than in insectivores and frugivore-insectivores; TriTaHI is significantly greater in faunivores, frugivore-insectivores, and insectivores than in folivores. These results are illustrated in Fig. 3, and full test results are reported in the Online Resource 2.

**Fig. 3.**
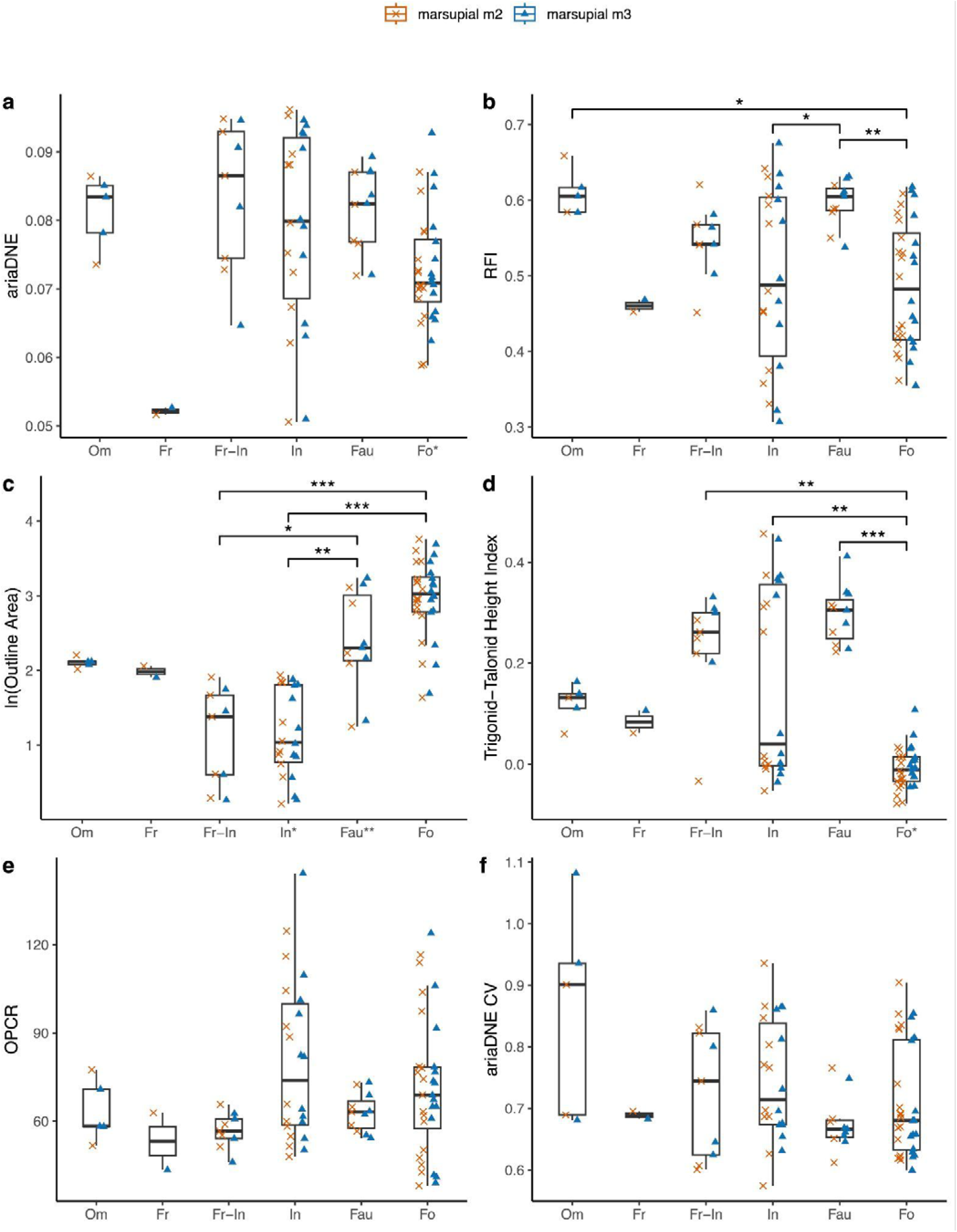
Dental metrics for marsupial sample. Boxplots are of the total marsupial sample (m2 + m3), whereas m2 values are shown as orange crosses, and m3 values as blue triangles. Dietary categories that differed significantly are marked by brackets and asterisks. Abbreviations: Om, omnivore; Fr, frugivore; Fr-In, frugivore-insectivore; In, insectivore; Fau, faunivore; and Fo, folivore. Dietary categories that differed significantly in dental metrics between the m2 and m3 *within* a given dietary category are marked with an asterisk at their category along the x-axis. * = *p* < 0.05; ** = *p* < 0.01; and *** = *p* < 0.001. Note that frugivores were excluded from both tests.

#### Principal component analyses

Table 5 shows the dietary predictive power of DTMs in isolation, highlighting that for the marsupial-only sample, ariaDNE outperforms both DNE and convex DNE (47.3% versus 13.5% and 21.6% classification accuracy, respectively). As we aim to use DTMs for their dietary predictive power, we elected to use ariaDNE over DNE or convex DNE for the principal component analysis (PCA); similarly, we excluded ariaDNE CV from the PCAs because of its low predictive power (< 25% accuracy).

**Table 5.** Dietary predictive power of individual dental metrics measured as classification accuracy of quadratic discriminant analysis per subset. The marsupial sample contains four dietary categories (frugivore-insectivore, insectivore, faunivore, folivore); the primate sample contains five dietary categories (hard-object feeder, frugivore, frugivore-insectivore, insectivore, folivore); the marsupial and primate sample contains seven dietary categories (omnivore, hard-object feeder, frugivore, frugivore-insectivore, insectivore, folivore, faunivore). Abbreviations: **RFI**, relief index; **ariaDNE**, a robustly implemented algorithm for Dirichlet normal energy; **DNE**, Dirichlet normal energy; **ln(OA)**, natural log of 2D outline area; **OPCR**, orientation patch count rotated; **CV**, coefficient of variation; **TriTaHI**, trigonid-talonid height index.

| Metric | Marsupial (m2 + m3)<br>[4 diets] |  | Primates<br>[5 diets] |  | Marsupial (m2 + m3) + primates<br>[7 diets] |  |
| --- | --- | --- | --- | --- | --- | --- |
|  | Specimen data | Species mean data | Specimen data | Species mean data | Specimen data | Species mean data |
| RFI | 43.2% | 33.3% | 39.1% | 33.3% | 32.0% | 21.1% |
| ariaDNE | 47.3% | 41.0% | 60.3% | 66.7% | 40.6% | 38.9% |
| DNE | 13.5% | 17.9% | 41.4% | 41.2% | 28.5% | 22.1% |
| Convex DNE | 21.6% | 20.5% | 42.5% | 41.2% | 35.2% | 22.1% |
| Size ln(OA) | 54.1% | 59.0% | 56.9% | 58.8% | 53.5% | 40.0% |
| OPCR | 41.9% | 33.3% | 43.7% | 41.2% | 37.1% | 25.2% |
| ariaDNE CV | 24.3% | 23.1% | 48.9% | 54.9% | 38.7% | 22.1% |
| TriTaHI | 56.8% | 61.5% | 36.2% | 51.0% | 24.2% | 23.2% |

Based on the box plots (Fig. 3) and the PCA (Fig. 4) including ariaDNE, RFI, and lnOA of marsupial data of both the m2 and m3 specimen data combined (referred to as ‘marsupial (m2 + m3)’ sample after this), we identify some general trends in marsupial molar shape: faunivores have large molars with high ariaDNE and RFI; folivores also have large molars that cover a wide range of low to high ariaDNE and RFI; our small omnivore sample (*n* = 5) exhibits large molars that fall entirely within the range of faunivores for ariaDNE and RFI (see Fig. 4). The small sample of frugivores consists of two medium-sized molars with low molar ariaDNE and medium RFI, both belonging to *Hypsiprymnodon moschatus*. Of the smaller-sized specimens, frugivore-insectivores tend to have high curvature and relief (high ariaDNE and RFI), whereas insectivores display a wide range of ariaDNE and RFI values, covering the entire range of the marsupial sample (see Fig. 3a-b).

**Fig. 4.**
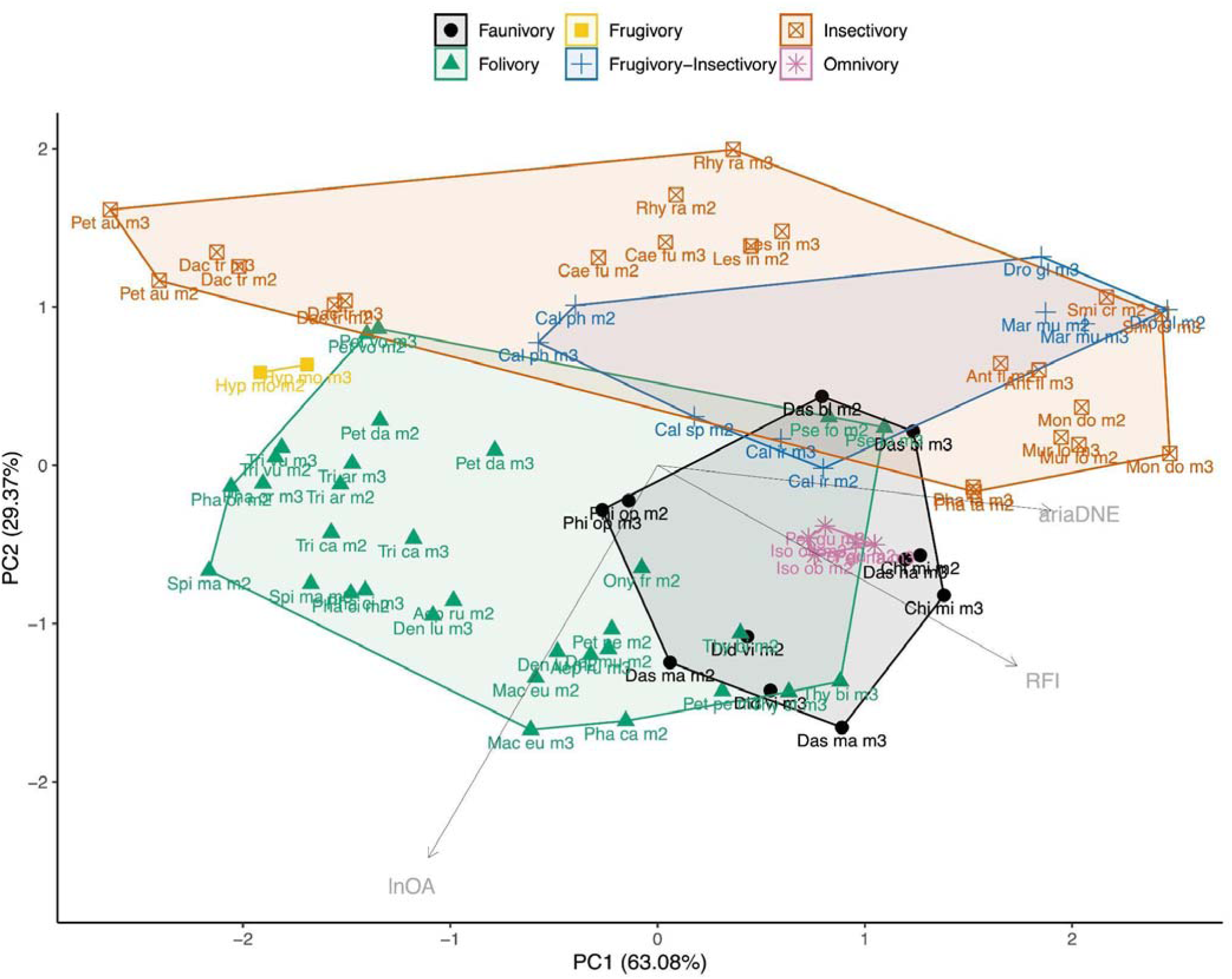
Principal component analysis of ariaDNE, RFI, and lnOA plot showing PC1 and PC2 scores for total marsupial (m2 + m3) specimen data, capturing 92.43% of the variation. Taxon abbreviations can be found in Table 3.

The insectivores occupy the largest area of any diet in the morphospace, showing three clusters: ‘group 1’, comprising the petaurids *Dactylopsila trivirgata* and *Petaurus australis*, with flat and blunt molars with very low RFI (< 0.4 RFI) and low ariaDNE (< 0.067 ariaDNE); ‘group 2’, comprising the caenolestids *Caenolestes fuliginosus*, *Lestoros inca*, and *Rhyncholestes raphanurus*, with molars that have medium RFI (0.43 < RFI < 0.5) and ariaDNE (0.072 < ariaDNE < 0.081); and ‘group 3’, comprising the didelphid *Monodelphis domestica,* and the dasyurids *Antechinus flavipes, Murexia longicaudata,* and *Smithopsis crassicaudata*, with molars characterized by very high relief (> 0.56 RFI) and curvature (> 0.088 ariaDNE), both at the highest end of the marsupial range (see Online Resource 2: Fig. S4 for an example of the molar shape of each group). These results suggest a strong phylogenetic influence on lower molar shape in insectivorous marsupials, as ‘group 1’ comprises only petaurids, and ‘group 2’ comprises only caenolestids. Although ‘group 3’ comprises species from two relatively distantly related families (Didelphidae and Dasyuridae), both of these are characterised by relatively unmodified tribosphenic molars, in contrast to the differently specialised molars of petaurids and caenolestids (Archer 1984a; b; Moore and Sanson 1995; Ungar 2010; Beck et al. 2022). Even though the convex hull of frugivore-insectivores appears to overlap that of insectivores (Fig. 4), the frugivore-insectivores occupy a space left mostly unoccupied by insectivore specimens, in the gap between the insectivore group 2 (caenolestids) and group 3 (*Monodelphis* and dasyurids) mentioned above.

The 2D TriTaHI metric we present here has high dietary predictive power; when used alone, it is 56.8% accurate in predicting diet for the marsupial (m2 + m3) sample (see Table 5). When TriTaHI is included in the PCA (see Fig. 5), folivores (low TriTaHI) are much better separated from faunivores (high TriTaHI) and omnivores (intermediate TriTaHI). Additionally, the overlap in morphospace between folivores and high TriTaHI frugivore-insectivores and some insectivores (see Fig. 3D) is reduced compared to the PCA excluding this metric (see Fig. 4). The grouping of insectivores into three distinct clusters remains when TriTaHI is included, and insectivore ‘group 3’ is separated from the other insectivores by having a markedly higher trigonid than talonid (TriTaHI > 0.26) compared to members of both group 1 and 2 (which have similar values: -0.05 < TriTaHI < 0.06; see Fig. 3d for the bimodal distribution in insectivore TriTaHI), again suggesting a strong phylogenetic influence on molar morphology in insectivorous marsupials. See Online Resource 2: Fig. S1 for the PCA plot that also includes OPCR and Fig. S2 for PCA plots of marsupial m2-only and m3-only data.

**Fig. 5.**
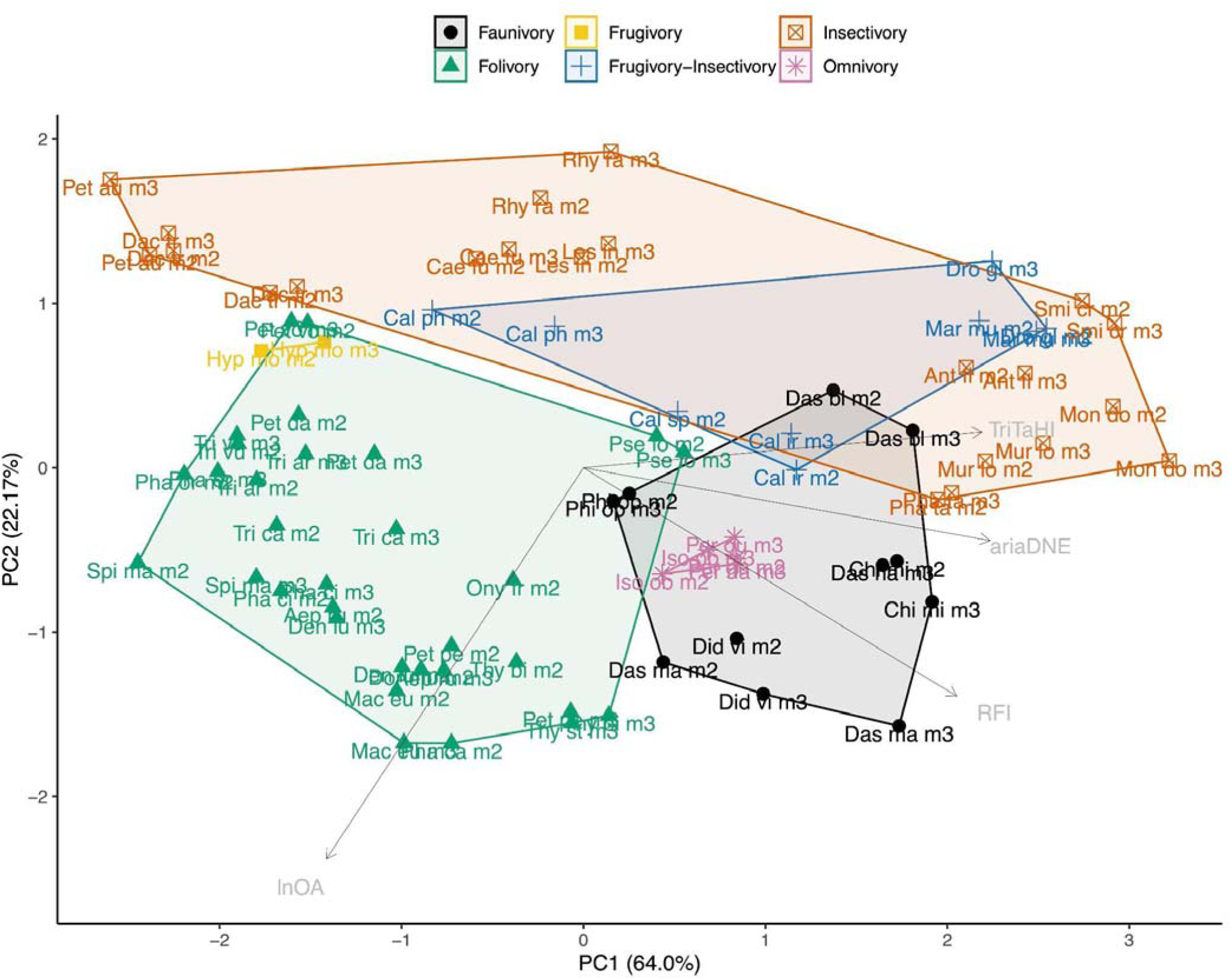
Principal Component Analysis of ariaDNE, RFI, lnOA, and TriTiHI plot showing PC1 and PC2 scores for total marsupial (m2 + m3) specimen data, capturing 86.17% of the variation. Note that the PC loading arrows are not to scale. Taxon abbreviations can be found in Table 3.

#### Dietary classification accuracy

Accuracy of dietary classification of the marsupial sample using leave-one-out QDA on species-mean data is relatively low when only RFI and ariaDNE are used (< 60%; see Table 6), particularly for the m2-only sample (40%; see Table 6). The addition of lnOA to the metrics RFI and ariaDNE results in a marked increase in classification accuracy: > 10% for each marsupial subset, reaching a classification accuracy as high as 69% for the m3-only sample. Incorporating other metrics results in smaller increases in accuracy, up to an accuracy of 62% for the m2-only sample (RFI + ariaDNE + lnOA + OPCR) and 69% for the combined m2 and m3 sample (RFI + ariaDNE + lnOA + TriTaHI). Despite not having significantly different values between m2 and m3 for most DTMs, the m3-only marsupial sample performs substantially better than the m2-only sample (>15% increase in classification accuracy in QDAs using ariaDNE + RFI and ariaDNE + RFI + lnOA; see Table 6). As our m2-only and m3-only samples include a slightly different set of taxa, we controlled for this and repeated the same tests on exactly the same set of taxa for the m2-only and m3-only samples. The results of these sub-set analyses confirm a higher dietary predictive power of the marsupial m3 over the m2 (see Online Resource 2: Table S3).

**Table 6.** Leave-one-out QDAs species means for marsupial samples (m2-only, m3-only, and total sample) and primate sample in isolation. See Online Resource 3: Table S2 for side-by-side results of QDAs using specimen-based data versus species averages. The marsupial samples (m2 only, m3 only, and m2 + m3) contained four dietary categories (folivory, insectivory, frugivory-insectivory, faunivory), unless marked with an *, when there were only three categories (folivory, insectivory, faunivory). The primate sample contained five dietary categories (hard-object feeder, insectivory, folivory, frugivory, frugivory-insectivory). Abbreviations: **RFI**, relief index; **ariaDNE**, a robustly implemented algorithm for Dirichlet normal energy; **OPCR**, orientation patch count rotated; **CV**, coefficient of variation; **TriTaHI**, trigonid-talonid height index.

| Metric | Marsupial m2-only | Marsupial m3-only | Marsupial m2 + m3 | Primate |
| --- | --- | --- | --- | --- |
| RFI + ariaDNE | 40.5% | 57.1% / 61.3% * | 46.2% | 62.7% |
| RFI + ariaDNE + size | 51.4% | 68.6% / 77.4% * | 64.1% | 80.4% |
| RFI + ariaDNE + size + TriTaHI | 56.8% | 80.6% * | 69.2% | 74.5% |
| RFI + ariaDNE + size + ariaDNE CV | 51.4% | 80.6% * | 66.7% | 76.5% |
| RFI + ariaDNE + size + OPCR | 62.2% | 77.4% * | 61.5% | 74.5% |

For the marsupial m3-only sample, QDAs with four or more metrics included required an additional dietary category to be excluded (frugivore-insectivore) due to its small sample size. Unsurprisingly, this resulted in higher classification accuracies, as only three rather than four diets were tested. Thus, m3-only QDAs with more than three variables are not directly comparable with those of the m2-only and combined (= ‘m2 + m3’) samples. Nevertheless, our PCA plots for these input variables also support higher dietary prediction accuracy for the marsupial m3 over the m2 (see Online Resource 2: Fig. S2) when more than three input variables are used, with m3-only data showing less overlap in morphospace between faunivores and frugivore-insectivores; there is also less overlap between faunivores and insectivores with the m3-only data when OPCR is included (see Online Resource 2: Fig. S2). Finally, for the combined m2 + m3 sample, there is little difference in classification accuracy between RFI + ariaDNE + lnOA (64%), RFI + ariaDNE + lnOA + TriTaHI (69%), RFI + ariaDNE + lnOA + ariaDNE CV (67%), and RFI + ariaDNE + lnOA + OPCR (62%).

### Primate dental topography

Numerous studies have investigated the performance of most of most dental metrics used here in primates (Boyer 2008; Bunn et al. 2011; Winchester et al. 2014; St Clair and Boyer 2016; Shan et al. 2019; Fulwood et al. 2021a; b; de Vries et al. 2024a; Selig et al. 2024), and so we do not present a detailed reanalysis and report of these here. We report that our new TriTaHI metric is, on average, highest in primate insectivores (TriTaHI > 0.05, with all but two specimens > 0.1), second-highest in folivores (0.01 < TriTaHI > 0.16), second-lowest in hard-object feeders (-0.06 > TriTaHI < 0.1, see Fig. 6D), and lowest in frugivores (-0.06 > TriTaHI < 0.05), similar to the general pattern observed for ariaDNE and RFI. In contrast, for ariaDNE and RFI, it is the hard-object feeders that have the lowest and bluntest crowns, whereas frugivores have a considerably higher ariaDNE and RFI values on average than the hard-object feeders (which Winchester et al. 2014 found to be significantly different), although still low compared to the other diet groups. In contrast, the TriTaHI results show that hard-object feeders, despite having on average the lowest ariaDNE and RFI, show a greater difference in height between the trigonid and talonid than do frugivores. In isolation, the TriTaHI metric predicts diet in primates with 51% accuracy, which is 10% lower than for marsupials. Primates do include an additional dietary category, which may account for the lower predictive power in this group (see Table 5).

**Fig. 6.**
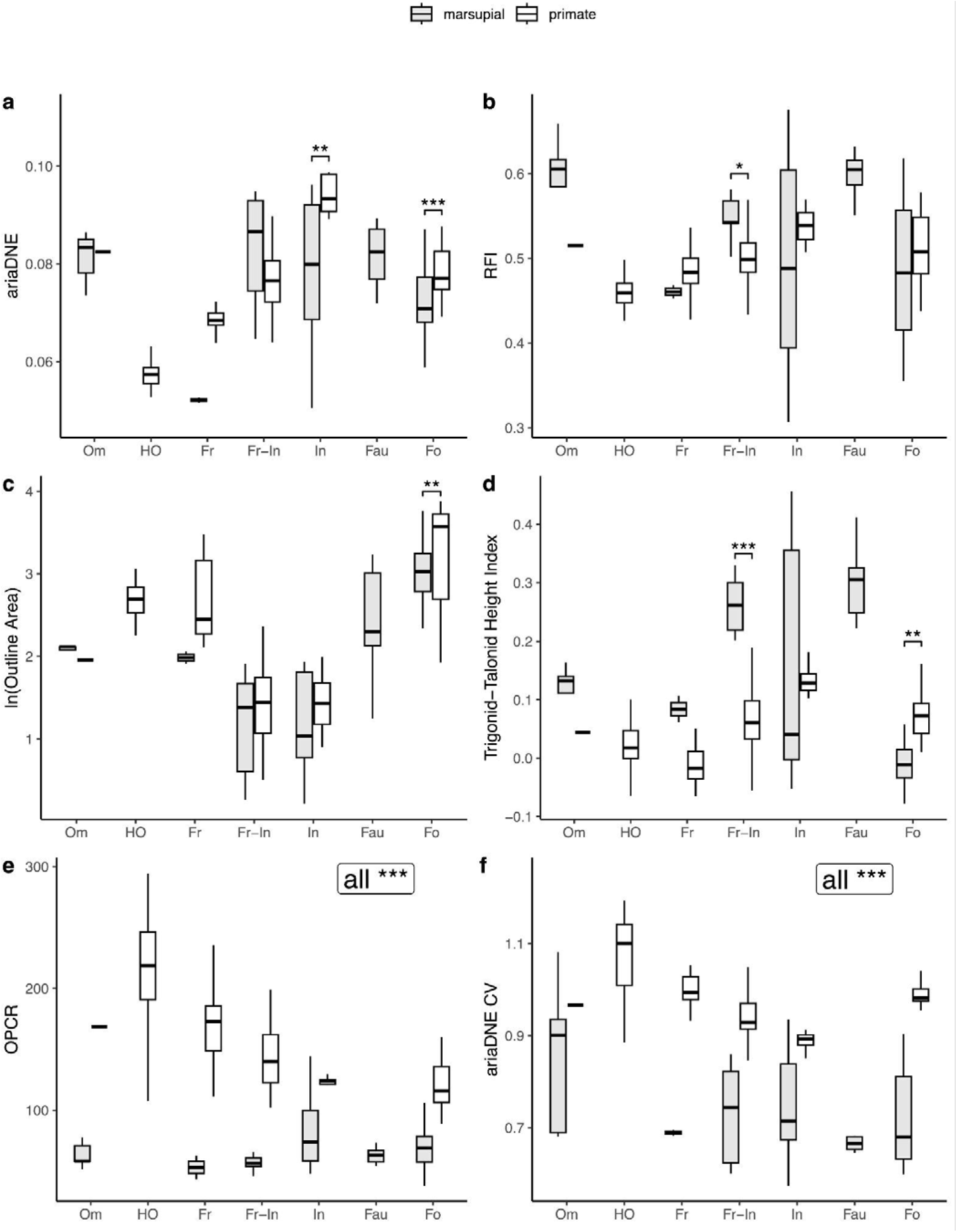
Boxplots of marsupial (m2 + m3) data in grey and primate (platyrrhine and strepsirrhine) in white, both displaying specimen data distribution for different diets. All significantly different pairs are marked by brackets and asterisks (*p* < 0.05). Abbreviations: Om, omnivore; Fr, frugivore; Fr-In, frugivore-insectivore; In, insectivore; Fau, faunivore; and Fo, folivore. * = *p* < 0.05; ** = *p* < 0.01; and *** = *p* < 0.001. Note that omnivores, hard-object feeders, frugivores, and faunivores were excluded from the t-tests due to small sample sizes. See Online Resource 2: Fig. S3 for individual data points plotted over the boxplots.

It is notable that QDA classification accuracies are consistently around 10% higher for the primate sample than for any of the marsupial samples, even though the primate-only QDAs include an additional dietary category (five in total rather than four for most of the marsupial-only QDAs).

### Comparing marsupial and primate dental topography

#### Issues with OPCR and ariaDNE CV across primates and marsupials

The Welch t-tests that we ran on specimen data for the frugivore-insectivores, insectivores, and folivores show OPCR and ariaDNE CV to be significantly lower for marsupials than primates (all *p* < 0.001), which can also be observed in the boxplots of Fig. 6e-f. We strongly suspect that our OPCR and ariaDNE CV results (as well as DNE and ConvexDNE to a lesser extent, see Online Resource 2: Fig. S5) reflect sensitivity of these metrics to sample material, since epoxy specimen casts were used for all primates, whereas original specimens were used for all marsupials (this is also observable in Fig. 2, with marsupial teeth appearing smoother than the primate casts). The epoxy primate casts exhibit consistently higher OPCR and ariaDNE CV values than the original marsupial specimens, even after meshes were simplified and smoothed consistently. This confirms an earlier report of this phenomenon by López-Torres et al. (2018, figure 3) for OPCR in plesiadapiform primates. We therefore recommend caution when interpreting OPCR and ariaDNE CV results of analyses that include different sample materials, as we believe these metrics not only reflect molar shape, but also sample material (see also López-Torres et al. 2018). For this reason, we exclude OPCR and ariaDNE CV from descriptions and analyses of the combined marsupial and primate sample. However, we consider these metrics suitable for our marsupial-only and primate-only analyses, given that specimen type (original for marsupials, epoxy replica casts for primates) is consistent within these groups.

#### Boxplots

It is striking that there is clear overlap between primates and marsupials in the same dietary categories for RFI, ariaDNE, and lnOA, but very little overlap for TriTaHI (see Fig. 6a-d). Marsupials consistently exhibit higher values of TriTaHI than primates for each dietary category except for the marsupial folivores, which exhibit lower TriTaHI values than do primate folivores. The Welch t-tests on specimen data show TriTaHI to be significantly different for the frugivore-insectivores (higher for marsupials, lower for primates; *p* < 0.01) and folivores (lower for marsupials, higher for primates; *p* < 0.01), whereas the difference in TriTaHI values is not significant between insectivorous marsupials and primates, mostly due to the great variation in marsupial insectivores. It can also be seen that when the marsupial and primate samples are combined, the TriTaHI metric experiences a considerable drop in dietary predictive power when used alone: from > 50% in marsupial-only and primate-only samples to 23% accuracy when marsupial and primate samples are combined (see Table 5). Although there is some overlap in ariaDNE between marsupials and primates for most dietary categories, it is only the frugivore-insectivores for which there is no statistical difference between marsupial and primate ariaDNE, with a *p-*value > 0.05 of the Welch t-test. The insectivores and folivores differ significantly in ariaDNE between marsupials and primates (*p* < 0.01), with marsupials in these dietary categories having lower average curvature than primates do. Relief Index values also overlap between marsupials and primates for most diets, except for the RFI of frugivore-insectivores, for which marsupials have a significantly higher RFI than primates (*p* < 0.001). Molar size, measured as lnOA, appears to be most similar between marsupials and primates with the same diets, and the only significant difference is between folivorous marsupials and primates (*p* = 0.044). For these categories, the ranges of lnOA values are fairly similar, but overall, the average primate folivore molar is larger than the average marsupial folivore molar. Upon closer inspection of the distributions (see Online Resource 2: Fig. S3), it is clear that the larger average molar size of primate folivores is driven by the platyrrhine folivore sample (3 species; *n* = 20).

#### Principal Component Analyses

The PCA plot of the species means of ariaDNE, RFI, and lnOA is shown in Fig. 7, and the PCA plot of those three metrics plus TriTaHI is shown in Fig. 8. The lnOA metric separates dietary categories consistently across marsupials and primates roughly along the y-axis (PC2), with folivores, faunivores, omnivores, and hard-object feeders being larger on average (lnOA > 1.9) than frugivore-insectivores and insectivores (lnOA < 1.9). The exception to this pattern are frugivores; the primate frugivores group with the larger species, whereas the sole marsupial frugivore (*Hypsiprymnodon moschatus*, which has a body mass of ∼500g, Johnson and Strahan 1982) groups with the smaller species. Marsupial frugivore-insectivores and insectivores show a greater overlap in morphospace than do their primate counterparts (Figs. 7-8). This is due to marsupial frugivore-insectivores *Dromiciops gliroides* and *Marmosa murina* extending the medium RFI and ariaDNE range seen in primate frugivore-insectivores into the higher RFI and ariaDNE range; in primates, this is only occupied by insectivorous species. Conversely, whereas most primate insectivores exhibit high RFI and ariaDNE (RFI > 0.45, ariaDNE > 0.075), this is only true for about half of the marsupial insectivores; the other half in our sample show medium or even low RFI and ariaDNE (see group 1 and 2 above, with RFI as low as 0.31 and ariaDNE as low as 0.051), resulting in more overlap between frugivore-insectivore and the insectivore categories for marsupials In fact, marsupial insectivore group 1 (formed by the petaurids *Dactylopsila trivirgata* and *Petaurus australis*, both of which are characterised by low, blunt molars; see Online Resource 2: Fig. S4) falls outside both the primate frugivore-insectivore and insectivore ranges on PC1 and overlaps primate hard-object feeders.

**Fig. 7.**
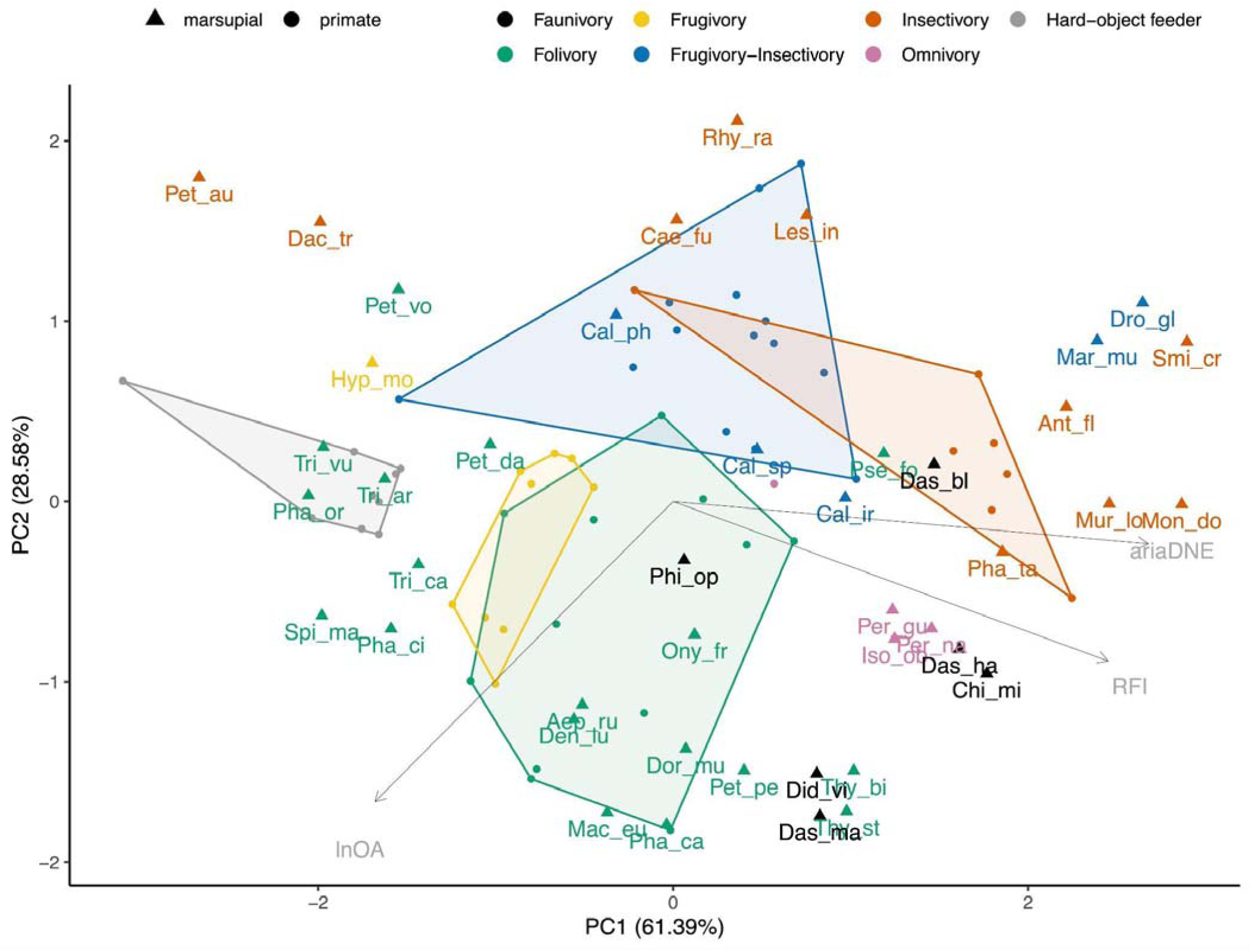
Principal Component Analysis of ariaDNE, RFI, and lnOA plot showing PC1 and PC2 scores for total marsupial (m2 + m3) and primate species mean data, capturing 89.97% of the variation. Note that the PC loading arrows are not to scale. Triangles are marsupial data, dots are primate data, convex hulls are primate-only. Taxon abbreviations can be found in Table 3.

**Fig. 8.**
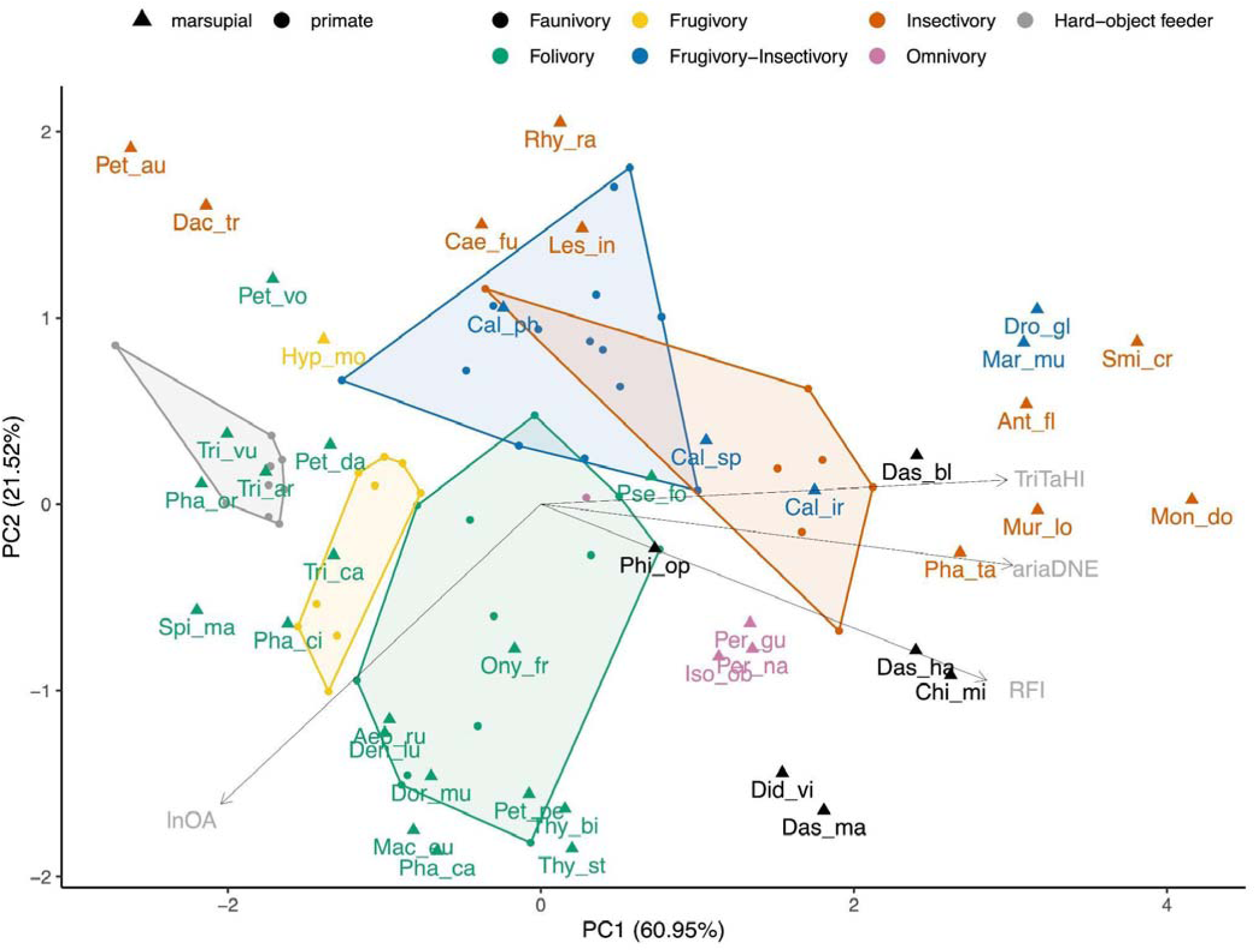
Principal Component Analysis of ariaDNE, RFI, lnOA, and TriTaHI plot showing PC1 and PC2 scores for total marsupial (m2 + m3) and primate species averaged data, capturing 82.47% of the variation. Note that the PC loading arrows are not to scale. Triangles are marsupial data, dots are primate data, convex hulls are primate-only. Taxon abbreviations can be found in Table 3.

Marsupial folivores occupy a considerably larger area in the morphospace than their primate counterparts do, and this is driven particularly by greater variation in PC1 (ariaDNE, RFI, and TriTaHI). For molar size (lnO; mostly PC2), marsupial folivores show a similar but slightly broader range than primates (marsupial lnOA range: 1.6-3.8; primate lnOA range: 1.9-3.9). The primate folivores are split; strepsirrhine folivores have small-to medium-sized m2s (lnOA range: 1.9-3.5), whereas platyrrhine folivores have medium-to large-sized molars only (lnOA range: 3.4-3.9). When marsupials and primates are both included in the PCA, four clusters of marsupial folivore molar shapes can be identified that are not as distinct in the marsupial-only PCA. Two of these each comprise a single smaller-sized folivorous species: *Pseudochirulus forbesi* and *Petauroides volans.* Both *Pseudochirulus* and *Petauroides* are members of Pseudocheiridae, and have lnOA ranging from 1.64 to 2.09, comparable to only a single folivorous primate in our sample, the strepsirrhine *Lepilemur leucopus* (lnOA = 1.93). Between these two smaller-sized marsupial folivores, *Pseudochirulus* has relatively high RFI (> 0.51), in contrast to *Petauroides* (RFI < 0.36). The larger-sized marsupial folivores (lnOA range: 2.34-3.76) form two groups: (1) the macropodoids *Aepyprymnus rufescens*, *Dendrolagus lumholtzi*, *Dorcopsis muelleri*, *Macropus eugenii*, *Onychogalea frenata*, *Petrogale penicillata*, *Thylogale billardieri*, *Thylogale stigmatica*, and the phalangerid *Phalanger carmelitae*, all of which are large-sized and have high RFI values (> 0.49); (2) non-macropodoids, including the pseudocheirid *Petropseudes dahli*, the phascolarctid (koala) *Phascolarctos cinereus*, and the phalangerids *Phalanger orientalis*, *Spilocuscus maculatus*, *Trichosurus arnhemensis*, *Trichosurus caninus*, and *Trichosurus vulpecula*, most of which are large-sized but have low RFI values (< 0.47). The macropodoid group, with higher RFI values, includes marsupials that include grasses in their diet, whereas most non-macropodoids, which have lower RFI values, do not include grasses in their diet (see Discussion). Of the four marsupial folivore groups, it is only the macropodoid folivore group (i.e., large-sized with high RFI) that partially overlaps with primate folivores in morphospace. Both *Petauroides* and the group of non-macropodoid marsupial folivores fall outside the range of primate folivores and have lower RFI. Some members of the non-macropodoid group (*Trichosurus arnhemensis*, *Trichosurus vulpecula,* and *Phalanger orientalis*) overlap with primate hard-object feeders in morphospace because they share low RFI values. On the other hand, *Pseudochirulus* has higher RFI and falls within the convex hull of primate insectivores, possibly due to its small molar size (lnOA).

The marsupial omnivores plot away from the single primate omnivore (*Nycticebus javanicus*) due to their larger molar size and higher RFI values. Marsupial faunivores show a wide range of molar shapes; one species overlaps primate insectivores (*Dasycercus blythi*), and another overlaps primate folivores (*Philander opossum*). The other four marsupial faunivores group close to marsupial omnivores (*Dasyurus hallucatus* and *Chironectes minimus* due to their medium size and high RFI) or to macropodoid marsupial folivores (*Dasyurus maculatus* and *Didelphis virginiana*, due to their large size and high RFI).

### Marsupials and primates combined

#### Classification accuracy

Quadratic discriminant analyses of the combined marsupial and primate samples show a considerable improvement in classification accuracy when lnOA is included (average of 13.4% accuracy increase), as well as a considerable improvement in classification accuracy when the TriTaHI metric is included. This pattern occurs regardless of whether the marsupial sample includes only m2, only m3, or both (see Table 7). The primate plus marsupial m3-only sample performs slightly better than primate plus the marsupial m2-only sample (see Table 7), and the primate plus marsupial m3-only sample performs as well as the primate plus marsupial m2 + m3 sample. The highest classification results are achieved when ariaDNE, RFI, lnOA, and TriTaHI are included (70.1% for primates + marsupial m3-only; 69.2% for primates + marsupial m2 + m3). When the classification accuracies of each dietary category are considered (see Online Resource 2) adding TriTaHI improves the accuracy of the folivore category but slightly lowers the accuracy of the faunivore category, resulting in an overall improvement of 4.4-6.9% (see Table 7).

**Table 7.** Leave-one-out QDAs species averages for combined marsupial and primate sample. All QDAs included six dietary categories (faunivory, folivory, frugivory, frugivory-insectivory, hard-object feeder, insectivory). Breakdown of accuracy per dietary category in Supplementary Materials. See Table S2 for side-by-side results of QDAs using specimen-based data versus species averages. Abbreviations: **RFI**, relief index; **ariaDNE**, a robustly implemented algorithm for Dirichlet normal energy; **TriTaHI**, trigonid-talonid height index.

| Metric | Marsupial m2s + Primates | Marsupial m3s + Primates | Marsupial all + Primates |
| --- | --- | --- | --- |
| RFI+ariaDNE | 47.2% | 52.9% | 48.4% |
| RFI+ariaDNE+size | 60.7% | 63.2% | 64.8% |
| RFI+ariaDNE+size+TriTaHI | 67.4% | 70.1% | 69.2% |

#### Cross-validated tests: classifying primates using the marsupial training set

The marsupial (m2 + m3) species-mean data set includes faunivores, folivores, frugivore-insectivores, and insectivores. Test accuracy was assessed based only on dietary categories present in both the marsupial and primate data sets (i.e., folivory, frugivory-insectivory, and insectivory were included; frugivory, hard-object feeding, and omnivory were excluded). The results of the cross-validated tests of the different marsupial subsets (m2 + m3, m2-only, m3-only) follow the same general pattern. We present the results of analyses of the marsupial m2 + m3 sample here and those of the marsupial m2-only and m3-only samples in the supplementary materials (see Online Resource 2: Table S4-6).

Highest test accuracy is achieved by including ariaDNE, RFI, and lnOA (69.7%; see Table 8, S4-6). Even though including TriTaHI increases training accuracy, it reduces the test accuracy, as, for most diets, marsupials show consistently higher TriTaHI values than primates for equivalent diets. The best model (i.e., ariaDNE, RFI, and lnOA) classifies primate folivores and insectivores with high accuracy (> 80%; see Table 9 for classification breakdown per diet) but struggles to correctly classify primate frugivore-insectivores (50% accurate). Primate frugivore-insectivores are usually misclassified as insectivores (see Table 9). Among dietary categories not present in the training set (see Table 9), primate frugivores are mostly misclassified as folivores (67% of the time); this is consistent with their morphospace in Fig.7, which falls within the marsupial folivore morphospace based on their shared larger size and low RFI and ariaDNE. Primate hard-object feeders are misclassified as folivores when marsupial m2 and m3 are assessed in isolation (see Online Resource 2: Tables S5-6) but mostly as frugivore-insectivores for the marsupial m2 + m3 sample. The sole primate omnivore is classified as a frugivore-insectivore. Specimen data are presented in Online Resource 2: Table S4.

**Table 8.**
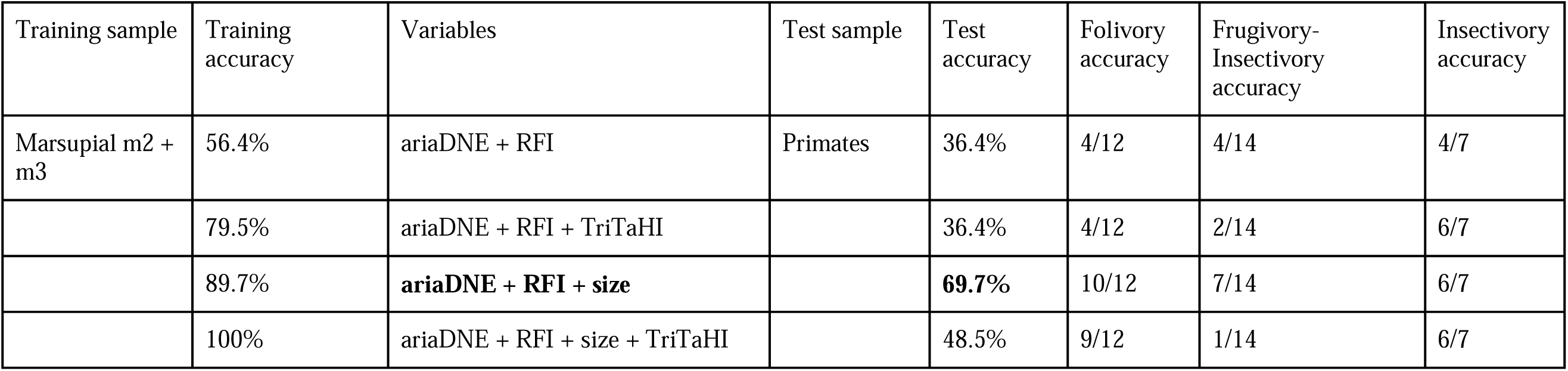
Classification accuracy of species-average QDAs with marsupial samples as the training set (using the entire training sample, not leave-one-out as in Table 7) and the total primate sample as the test sample. Training sets included faunivory, folivory, frugivory-insectivory, and insectivory categories. Accuracy of the QDA test was calculated using shared categories only, i.e., Folivory, insectivory, and frugivory-insectivory. See Table S3 for species average QDA results side by side with specimen data QDA results. Bolded text indicates the QDA settings of the model with highest test accuracy. Abbreviations: **RFI**, relief index; **ariaDNE**, a robustly implemented algorithm for Dirichlet normal energy; **TriTaHI**, trigonid-talonid height index.

| Training sample | Training accuracy | Variables | Test sample | Test accuracy | Folivory accuracy | Frugivory-Insectivory accuracy | Insectivory accuracy |
| --- | --- | --- | --- | --- | --- | --- | --- |
| Marsupial m2 + m3 | 56.4% | ariaDNE + RFI | Primates | 36.4% | 4/12 | 4/14 | 4/7 |
|  | 79.5% | ariaDNE + RFI + TriTaHI |  | 36.4% | 4/12 | 2/14 | 6/7 |
|  | 89.7% | **ariaDNE + RFI + size** |  | **69.7%** | 10/12 | 7/14 | 6/7 |
|  | 100% | ariaDNE + RFI + size + TriTaHI |  | 48.5% | 9/12 | 1/14 | 6/7 |

**Table 9.**
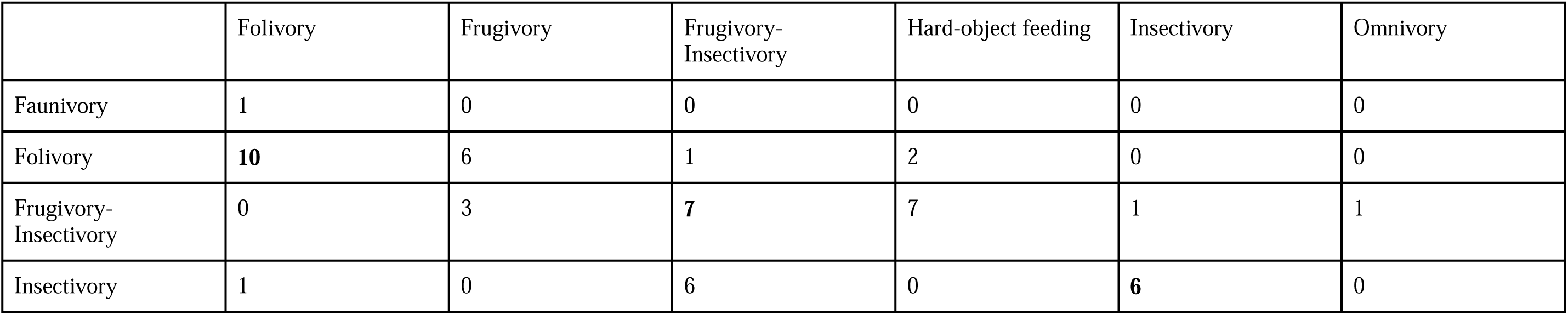
QDA category-specific results with ariaDNE, RFI, size as metrics. Training sample = marsupial m2 + m3 sample (vertical categories), test sample = primate sample (horizontal categories). Bolded numbers show the results of overlapping diets.

#### Cross-validated tests: classifying marsupials using the primate training set

Using species-mean data, the platyrrhine-only and strepsirrhine-only samples could not be used as training sets in isolation because the sample sizes of various dietary categories were too small. We thus used the total primate sample as the training set. Marsupials were classified correctly based on primate training data 61.8% of the time (including ariaDNE, RFI, and lnOA; see Table 10), which is slightly lower than the most accurate reverse model (69.7% accuracy when primates were classified using a marsupial training set; see Table 8). However, excluding the additional dietary category in these marsupial tests (frugivory, see Table 10) results in greater accuracy of 65.6%. Although this is slightly less accurate than the reverse analysis (69.7%; Table 8), it is comparable. We report the results of specimen data analyses, which confirm the findings of the species-mean analyses, in Online Resource 2: Table S7.

**Table 10.**
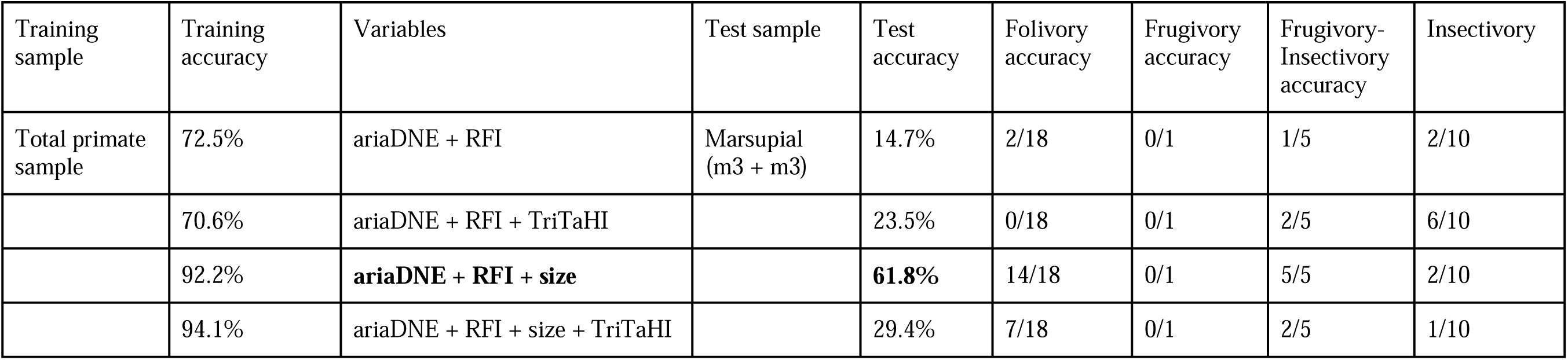
Classification accuracy of species-average QDAs with total primate sample as the training set (using the entire training sample, not leave-one-out as in Table 7) and the total marsupial sample as the test sample. Training set of the primate sample included the folivory, frugivory, frugivory-insectivory, hard-object feeder, and insectivory categories. Accuracy of QDA test was calculated using shared categories only, i.e., folivory, frugivory, frugivory-insectivory, and insectivory. See Table S4 for species average QDA results side by side with specimen data QDA results. Bolded text indicates the QDA settings of the model with highest test accuracy. Abbreviations: **RFI**, relief index; **ariaDNE**, a robustly implemented algorithm for Dirichlet normal energy; **TriTaHI**, trigonid-talonid height index.

| Training sample | Training accuracy | Variables | Test sample | Test accuracy | Folivory accuracy | Frugivory accuracy | Frugivory-Insectivory accuracy | Insectivory |
| --- | --- | --- | --- | --- | --- | --- | --- | --- |
| Total primate sample | 72.5% | ariaDNE + RFI | Marsupial (m3 + m3) | 14.7% | 2/18 | 0/1 | 1/5 | 2/10 |
|  | 70.6% | ariaDNE + RFI + TriTaHI |  | 23.5% | 0/18 | 0/1 | 2/5 | 6/10 |
|  | 92.2% | **ariaDNE + RFI + size** |  | **61.8%** | 14/18 | 0/1 | 5/5 | 2/10 |
|  | 94.1% | ariaDNE + RFI + size + TriTaHI |  | 29.4% | 7/18 | 0/1 | 2/5 | 1/10 |

The best model (i.e., based on ariaDNE, RFI, and lnOA; Table 10) shows that marsupial folivores are classified with 78% accuracy, the single marsupial frugivore taxon is misclassified as a hard-object feeder, all marsupial frugivore-insectivores are classified correctly, and most (80%) marsupial insectivores are misclassified as frugivore-insectivores instead (the remaining 20% are classified correctly; see Table 11). The high classification accuracy of marsupial folivores based on a primate training set is confirmed by the PCA plot, in which both groups overlap (Fig. 7). The same plot (Fig. 7) shows that half of the marsupial frugivore-insectivores do not overlap primates of the same diet, and some plot far away. The high classification accuracy of marsupial insectivores (100%) is likely due to frugivore-insectivores being the ‘default’ category of the QDA model for small specimens, based on inspection of the 2D distribution of the QDA models (see Online Resource 2: Fig. S7 and note the large area occupied by frugivore-insectivores). Among marsupial dietary categories absent from the primate training set, faunivores are classified as frugivore-insectivores in 67% of cases, and folivores in the remaining 33% of cases, whereas the three marsupial omnivores are all classified as frugivore-insectivores.

**Table 11.** QDA category-specific results with ariaDNE, RFI, size as metrics. Training sample = primate sample (vertical categories), test sample = marsupial m2 + m3 sample (horizontal categories). Bolded numbers show the results of overlapping diets.

|  | Faunivory | Folivory | Frugivory | Frugivory-Insectivory | Insectivory | Omnivory |
| --- | --- | --- | --- | --- | --- | --- |
| Folivory | 2 | **14** | 0 | 0 | 0 | 0 |
| Frugivory | 0 | 1 | **0** | 0 | 0 | 0 |
| Frugivory-Insectivory | 4 | 1 | 0 | **5** | 8 | 3 |
| Hard-object feeding | 0 | 1 | 1 | 0 | 0 | 0 |
| Insectivory | 0 | 1 | 0 | 0 | **2** | 0 |

## DISCUSSION

Our study, which incorporates six out of the seven orders and 12 out of 21 families, is the broadest study of marsupial dental topography yet published. Our analyses provide new information about several aspects of marsupial m2 topography versus that of the m3, sensitivity of some DTMs to non-biological factors, marsupial dental topographic metrics in relation to diet, and a comparison of marsupial dental topography to that of primates with similar diets.

### Marsupial m2 versus m3

Marsupial m2s and m3s are usually more similar to each other in size and shape than either is to m1 or m4 (see e.g., figures in Archer 1984a; b). Specifically, m1 (likely a retained deciduous premolar) is often quite distinct from other lower molars (e.g., Beck et al. 2022: ch. 159-163), and m4 is usually much smaller, particularly the talonid, although it is typically the largest tooth in specialised faunivores (e.g., the thylacinid Thylacinus cynocephalus, Warburton et al. 2019). We found few significant differences between marsupial m2s and m3s, confirming that the general resemblance of these loci extends to quantitative dental topographic metrics and size: only four of 30 m2-m3 metric pairs differed significantly. Size of m2 and m3, measured as lnOA, only differed significantly between insectivores and faunivores; with m3 larger than m2 in faunivores and m2 larger than m3 in insectivores. This pattern may be the result of opposite inhibitory cascade models of dental development (Kavanagh et al. 2007): increasing inhibition in insectivores (resulting in m2 > m3) and a weak inhibitory cascade in faunivores (resulting in m2 < m3). Besides these size-related differences, the m3s of folivores have a significantly higher curvature (measured as ariaDNE) and greater trigonid and talonid height difference (TriTaHI) than m2s.

Despite few differences in DTMs between m2 and m3, our QDA results support a stronger dietary signal in m3. This is supported by both marsupial-only analyses and combined marsupial-primate analyses. The m2 erupts before m3 (see e.g., Kingsmill 1962; Guiler and Heddle 1974; van Nievelt and Smith 2005; Astúa and Leiner 2008; Kido et al. 2018), and typically shows greater wear, which could influence dental topography (Pampush et al. 2018; Li et al. 2020; Morse et al. 2023). However, as we used only unworn to lightly worn teeth in our study, any such effects should be minimal. Considering meristic gradients, m3 often shows a slightly more exaggerated or “extreme” morphology than m2. For example, the height difference between the trigonid and talonid is greater in m3 than m2 in some insectivorous (e.g., *Antechinus flavipes;*; Werdelin 1987: fig. 2A) and faunivorous marsupials (e.g., *Dasyurus viverrinus*; Werdelin 1987: fig. 2B; Beck and Taglioretti 2020). In support of this, for some of the the marsupial insectivores (*Antechinus, Murexia*, *Rhynolestes*) and faunivores (*Chironectes* and *Dasyurus*) in our comparative dataset, TriTaHI values are 10% greater for m3 than m2. Even so, this is not the case for the other members of these dietary categories, and TriTaHI did not differ significantly between m2 and m3 for either insectivores or faunivores as a group. Even when only including ariaDNE, RFI, and lnOA (i.e., excluding TriTaHI), classification accuracy was considerably higher for the m3-only sample than the m2-only sample (69% vs. 51%, respectively), showing that the difference in accuracy between m2 and m3 is not driven simply the result of differences in the relative heights of the trigonid and talonid in some of our specimens.

In summary, our QDA results suggest that the m3 typically results in more accurate dietary prediction or reconstruction than the m2 in marsupials. More DTM studies of marsupials are needed to confirm that this is not a sample-specific result. Regardless of the cause of the difference in classification accuracy results, the m2 still performs much better than chance (i.e., 25% accuracy, assuming four dietary categories). Therefore, m2s are still useful for dietary reconstruction when no m3s are available, as may be the case for many extinct marsupials and relatives (metatherians).

### Sensitivity of some DTMs to non-biological factors

Our analyses reveal a potential problem with using the dental topographic metrics OPCR and ariaDNE CV and - to a lesser extent - DNE and convex DNE. Specifically, values for these four metrics appear to be considerably affected by the type of specimen analysed (i.e., original versus epoxy cast), even after consistently simplifying and smoothing all surface meshes. This finding parallels an observation of López-Torres et al. (2018), who reported that OPCR values tend to differ between original fossil specimens and casts, with the former having consistently higher values, likely due to the rougher surface of casts compared to enamel. Our findings are congruent with this inference (see Fig. 6e-f and Online Resource 2: Fig. S5). We do, however, find that the remaining 3D-DTMs (lnOA, RFI, ariaDNE) are more robust to samples composed of mixed materials and scanning modalities and resolutions (see Table 5, Fig. 6 and Online Resource 2: Fig. S5).Our findings add to those from previous studies that suggest that OPCR and DNE may be less useful than RFI and ariaDNE in dental topographic studies due to their greater sensitivity to noise, mesh-preparation, and specimen material (Bunn et al. 2011; Winchester et al. 2014; Prufrock et al. 2016b; López-Torres et al. 2018; Pampush et al. 2018; Berthaume et al. 2019; DeMers and Hunter 2024). Based on these collective results, we consider RFI and ariaDNE to be the most reliable 3D-DTMs currently used for quantifying and comparing the shape of lower molars (as done here), particularly when using meshes produced from different scanning modalities and resolutions, and/or specimen materials.

### Dental topographic metrics and diet

#### General trends

In marsupials, curvature (ariaDNE) is particularly low in frugivores, relief (RFI) is usually highest in faunivores and omnivores, and size (lnOA) is greatest in folivores and faunivores and smallest in insectivores and frugivore-insectivores. These results are broadly similar to those of other dental topographic studies of both primates and other mammals (Boyer 2008; Bunn et al. 2011; Allen et al. 2015; Pineda-Munoz et al. 2017; López-Aguirre et al. 2022b). However, among dietary categories we tested (frugivore-insectivore, insectivore, and folivore), roughly half of the primate-marsupial pairs differed significantly in these three metrics (4/9; see Results). This contrasts with the results of Spradley and Phillips (2018), who found that m2s of phalangeriform marsupials and “prosimian” primates did not differ significantly in DNE and RFI but did differ in OPCR (significantly higher in phalangeriforms).

#### Insectivores

The greater dental shape variation of marsupial insectivores compared to that of primate insectivores is mostly driven by the distinctive morphology of two marsupial species: the petaurids *Petaurus australis* and *Dactylopsila trivirgata* (see Online Resource 2: Fig. S4A). In contrast to the other insectivores in our analysis, the m2 and m3 of these species are characterised by extremely low curvature (ariaDNE) and relief (RFI). If these species are excluded, the marsupial insectivore morphospace, as seen in Fig. 7 and 8, is much closer in size to that of primates.

The low ariaDNE and RFI values for *P. australis* may be related to a diet predominantly comprising liquids such as exudates and nectar that do not require mastication (Smith and Russell 1982; Goldingay 1986, 1990; Carthew et al. 1999). The molars of exudativorous primates like *Callithrix, Cebuella*, *Nycticebus, Perodicticus* and *Phaner* do not show the extremely blunt and simple morphology seen in *Petaurus*. This may be because these primates appear to consume a greater proportion of invertebrates than do *Petaurus* species. For example, exudates make up 83% of dietary observations of *Petaurus australis* and arthropods only 9%, (Carthew et al. 1999), versus an average of 52% exudates and 21% animal matter for exudativorous callitrichids (de Vries et al. 2024a), 32.5% exudates and 25% insects for *Nycticebus*, and 21% exudates and 65% insects for *Perodicticus* (Boyer 2008). Of the primate exudate feeders included in this study, *Phaner* most closely resembles *Petaurus* in proportions of exudate versus insect, with 65% exudates and only 17.5% insects in its diet (Hladik et al. 1980; Boyer 2008). Nonetheless, the molars of *Phaner* (ariaDNE = 0.075; RFI = 0.45) are much less blunt and low-crowned than those of *Petaurus* (ariaDNE = 0.051; RFI = 0.35). We also note that the m1 of *Petaurus australis* and other *Petaurus* species has a specialised trigonid dominated by the protoconid (Archer 1984a: figs. 202-204), which forms a shearing complex with the P3 of the upper dentition (Moore and Sanson 1995: fig. 3). This tooth could play a functional role in mastication fulfilled by more distal molars in exudativorous primates but would not have been captured in our m2 and m3 analyses.

The other petaurid included in our analysis, *Dactylopsila trivirgata,* is also characterised by extremely blunt and low-crowned molars that set it apart from other marsupial insectivores. In fact, Spradley (2017) referred to this species as an “evolutionary outlier” and excluded it from some analyses. Spradley’s (2017) results (based on a PCA using OPCR, DNE, RFI, and the natural log of m2 area) show *Dactylopsila* specimens plotting outside the range of all other marsupial insectivores, and within the range of marsupial frugivores. Although, these results may have been influenced by Spradley’s (2017) decision to classify *Petaurus australis* and *P. breviceps* as frugivores, whereas we classified *P. australis* as an insectivore (see justification for this in Methods: Dietary classification).

In contrast to *Petaurus* species (which are primarily exudate-feeders; see above), *Dactylopsila* appears to have a highly insectivorous diet, without significant intake of exudates. *Dactylopsila trivirgata* generally targets wood-boring larvae and extracts them with an elongated digit, similar to the aye-aye *Daubentonia* (Cartmill 1974; Rawlins and Handasyde 2002; Spradley 2017; St Clair et al. 2018), although it also feeds on social insects (Smith 1982). Whereas *Daubentonia* supplements its insectivorous diet with harder foods (notably *Canarium* seeds) and is classified as a hard-object feeder in this study (see Methods: Dietary justification for a discussion), *Dactylopsila trivirgata* is classified here as an insectivore based on its high intake of invertebrate prey (Rawlins and Handasyde 2002). It is possible that the molar morphology of *Dactylopsila trivirgata* has been retained from a more exudativorous ancestor (St Clair et al. 2018). The m1 of *D. trivirgata* resembles that of *Petaurus* species in having a prominent protoconid that forms a shearing complex with the P3 (Archer 1984a fig. 208). However, the P3 *D. trivirgata* is more bladelike than that of *Petaurus* species and is oriented obliquely, whilst an enlarged parastyle on M1 (absent in *Petaurus*) may also form part of the shearing complex with the m1 protoconid (Ride 1956: fig. 3; Archer 1984a: fig. 208; Beck et al. 2022: character 123); this increased emphasis on shearing relative to the condition in *Petaurus* may be an adaptation to a diet focused on soft-bodied invertebrates (see comments by Beck 2009: p7-8). Thus, like *Petaurus* (see above), the dietary signal of insectivory in *D. trivirgata* may be reflected to a greater extent in the teeth anterior to m2 and m3; this could be tested by future dental topographic analyses that consider these teeth (Pineda-Munoz et al. 2017; de Vries et al. 2024b). On the other hand, Moore and Sanson (1995) found that the dentition of *Petaurus breviceps* was not as efficient at breaking down insect larvae as that of the dasyurid *Dasyuroides byrnei*, which, unlike *Petaurus*, has a relatively generalised tribosphenic dentition. We also note that Smith (1982) reported that arthropod remains in the gut of *Dactylopsila trivirgata* were “remarkably intact”, implying little oral processing. Thus, the dentitions of *Petaurus* and *Dactylopsila* may indeed be less effective for masticating invertebrates than the tribosphenic dentitions of insectivorous didelphids and dasyurids. This highlights one of the fundamental difficulties in studying the relationship between dental shape and diet in mammals, namely that molar shape does not need to be optimal for a particular diet to be consumed. As stated by Muizon and Argot (2003) “Therefore, one should keep in mind that all deductions made from functional anatomy analysis only represent a hypothesis on what an animal could have done, not what it actually did” (Muizon and Argot 2003: pp44–45).

The remaining marsupial insectivores form two distinct clusters. One cluster comprises didelphids (order Didelphimorphia) and dasyurids (order Dasyuromorphia), both of which are characterised by a comparatively unspecialised tribosphenic dentition (Archer 1976; Beck et al. 2022) with medium curvature (ariaDNE = 0.072-0.08) and relief (RFI = 0.44-0.50). The other cluster comprises caenolestids (order Paucituberculata), the m2 and m3 of which are also recognisably tribosphenic but which show some distinctive features not seen in didelphids or dasyurids, including a posterolingually-deflected paracristid and a long, labially-deflected entocristid (Abello et al. 2021; Beck et al. 2022). These additional features contribute to a higher curvature (ariaDNE = 0.088-0.096) and relief (RFI = 0.57-0.68) that is outside the range of insectivorous didelphids and dasyurids. There is still relatively limited information on diet and feeding for most didelphids, dasyurids, and caenolestids (Flannery 1995; Gardner 2007; Voss and Jansa 2021; Dickman and Calver 2022; Lessa et al. 2022; Baker and Gynther 2023), so it is unclear whether there are any obvious differences in masticatory efficiency between these clades. However, since didelphids, dasyurids and caenolestids all exhibit relatively generalised, premolariform lower premolars (Beck et al. 2022), it seems unlikely that there are major differences in the way they use their premolars are used to process food items.

In summary, some evidence suggests that petaurid molars may be less efficient at breaking down invertebrate prey items than those of other insectivorous marsupials, hinting at a potential constraint on form in this clade (developmental, functional, and/or phylogenetic). To support this further, there is a need to apply dental topographic methods to m1 as well to assess whether functions of more distal molars in other marsupials have shifted to this locus in petaurids (Pineda-Munoz et al. 2017; de Vries et al. 2024b). There is as yet no evidence for variation in dental efficiency between the molars of didelphids, dasyurids and caenolestids, and the fact that caenolestids cluster separately from didelphids and dasyurids presumably reflects the long independent evolutionary history of Paucituberculata (although we note that the dentally more similar didelphids and dasyurids also diverged from each other early in the the history of Marsupialia, Beck 2022). Future analyses should attempt to explicitly take into account and/or model the impact of phylogeny, as Fulwood et al. (Fulwood et al. 2021b) attempted for strepsirrhine primates, as well as other tooth loci (de Vries et al. 2024b).

#### Folivores

Like insectivorous marsupials, folivorous marsupials exhibit more varied molar morphology than primates do (at least as measured by the DTMs used here) and exhibit greater morphospace occupation. Among the folivorous marsupials we analysed, several distinct molar types are represented: strongly selenodont in the koala (*Phascolarctos*) and ringtails (pseudocheirids), semi-lophodont (= “incipiently lophoid” or “bunolophodont”) in phalangerids and the potoroid *Aepyprmnus*, and strongly lophodont in macropodids (Beck et al. 2022). By contrast, none of the folivorous primates in our sample have strongly selenodont or strongly lophodont molars, although we note that our sample lacks cercopithecoids, which are (bi)lophodont (Delson 1975; Rasmussen et al. 2019).

Some of our marsupial folivores are mixed feeders that incorporate grasses in their diet (the potoroid *Aepyprymnus* and most macropodids; Arman and Prideaux 2015), and these species are characterised by more curved (higher ariaDNE) and higher-crowned molars (higher RFI) than those of marsupial folivores that do not feed on grasses; by contrast, none of the primate folivores included here are known to eat grasses (only a single living primate, the cercopithecoid Theropithecus gelada, habitually feeds on grasses, Fashing et al. 2014). This may explain why marsupial folivores occupy a greater region of morphospace. However, perhaps surprisingly, the non-grass-eating folivorous marsupials and primates do not overlap in morphospace. Instead, the non-grass-eating marsupial folivores (most phalangerids, the koala, and *Petropseudes dahli*, or the ‘non-macropodoids’ group, see above) have comparatively low-crowned and blunt teeth and overlap with primate frugivores and hard-object feeders, whereas the mixed-feeding marsupial folivores overlap or plot more closely to primate folivores. The pattern of lower sharpness and relief for softer diets like leaves and fruits versus higher sharpness and relief for tougher and more fibrous diets that include grasses seen in marsupial folivores is congruent with predictions based on dental function (Bunn et al. 2011).

The differences in molar topography between marsupial and primate folivores with similar diets (i.e., those that do not eat grasses) may be another example of clade-specific offsets in dental topography. Winchester et al. (2014) found that platyrrhines, on average, showed higher relief (RFI) and complexity (OPCR) than “prosimians” with the same diets, but they did not find the same offset for curvature (DNE) or shearing ratio and shearing quotient (Winchester et al. 2014). Winchester et al. (2014) also found a general trend of platyrrhine primates occupying narrower distributions of DTM values than “prosimian” primates, who occupied the extremes of the topographic ranges. We observe a similar pattern when comparing marsupial and primate insectivore and folivores: marsupial insectivores and folivores plot at the extremes of the topographic ranges, with primate insectivores and folivores in the middle of the morphospaces occupied by the marsupials of the same diet. We agree with the suggestion of Winchester et al. (2014) that the explanation of this phenomenon is most likely complex and suspect that it is the result of a combination of various factors such as differences in evolutionary rates and constraints within our subsamples, as well as differences in the ranges of ecological variation and competition. However, as noted before, our sample of primate folivores is somewhat limited (we lack any representatives of Cercopithecoidea or Hominoidea), and the larger morphospace occupied by marsupial insectivores is largely driven by the distinctive morphology of *Petaurus* (which is classified as an insectivore here, even though it is primarily exudativorous; see ‘Dietary classifications’ above) and *Dactylopsila.* Nevertheless, for the insectivore and folivore groups, our sample supports greater molar topographic diversity among marsupials than primates.

#### TriTaHI metric

We propose a simple 2D “trigonid-talonid height index” (TriTaHI) metric here, which reflects the observation that the height difference between the trigonid and talonid is typically greater in insectivores and faunivores than in folivores and frugivores, in both primates and marsupials (Zimicz 2012, 2014; St Clair and Boyer 2016; Goin et al. 2020). For our marsupial sample, we found that TriTaHI is indeed significantly greater in faunivores than in folivores, and this metric has a high dietary predictive power even in isolation. Furthermore, combining TriTaHI with 3D-DTMs helps better separate these marsupial faunivores and folivores in PCA plots and improves classification accuracy.

Marsupials and primates show quite different patterns of TriTaHI, with primate TriTaHI being considerably lower, thus having trigonids and talonids more similar in heights, than marsupials in most dietary categories (see Results). This suggests fundamental differences in tooth shape and is reflected in the marked decrease in classification accuracy of this metric when it is used with the combined marsupial and primate sample. Indeed, it has been remarked by several authors that crown primates and their stem relatives (“plesiadapiforms”) are typically characterized by lower TriTaHI values than those of more generalized tribosphenic mammals (e.g., Szalay 1968; Rose 2006). However, we found this general pattern to be reversed in the folivore category, with folivorous primates exhibiting a greater difference in trigonid and talonid height compared to folivorous marsupials. We also note that primate hard-object feeders, which on average tend to have the lowest sharpness and relief of the different primate diets, show higher TriTaHI compared to primate frugivores, which could functionally be related to processing harder foods. Despite clade-specific differences in this metric, TriTaHI may still prove useful in discriminating between dietary groups when comparing more closely related taxa and/or when combined with methods that incorporate phylogenetic information, such as the Bayesian multilevel approach of Fulwood et al. (2021b), and we encourage other researchers to investigate its performance when applied to other clades.

#### QDA prediction accuracies

In general, we found that the accuracy of dietary prediction using QDA was greater for our primate-only analyses than for our marsupial-only analyses, implying that the aspects of molar shape captured by the specific DTMs used here do not correlate as strongly with diet in marsupials as in primates. For example, ariaDNE is 67% accurate for primate species means and only 41% for marsupial species means (Table 5). By contrast, RFI and lnOA perform equally well for the marsupial-only and primate-only samples, although the marsupial-only sample includes one fewer dietary category (four diets) compared to the primate-only sample (five diets), and thus has a higher chance at success by chance alone (25% versus 20%). For marsupials, we also cannot rule out the possibility that ariaDNE values are affected by differences in scanning and material modalities and pick up on noise introduced by variable scanning resolutions and modalities for our marsupial sample (in contrast, materials and scanning modalities were consistent for the primate sample). However, there is no clear offset in ariaDNE values between original specimens and casts, unlike what we observed for values of OPCR, ariaDNE CV, and, to a lesser extent, DNE and convex DNE (Online Resource 2: Fig. S5). Regardless of whether marsupial molar morphology shows an overall weaker correlation with diet, or whether less accurate dietary classification found here is an artefact of the specific DTMs used, the fact that marsupial dietary groups show greater overlap in dental topographic values than do primates means that it is overall more difficult to accurately predict marsupial diet.

### Applications and limitations of our study

Based on the results of the combined marsupial and primate QDAs, which show a moderately accurate dietary prediction when combining the DTMs ariaDNE, RFI, and lnOA (65% accurate leave-one-out approach and six dietary categories, average of 66% with cross-validation approach with four or three dietary categories), we conclude that our combined comparative sample should allow the diet of fossil marsupials and their close fossil relatives (Metatheria) and primates and their close fossil relatives (“plesiadapiforms”) to be predicted with reasonable confidence. There are no specialist hard-object feeders among living marsupials, but some fossil metatherians have been suggested to have included hard-shelled seeds and/or nuts as a major component of their diet (Pascual et al. 1994; Goin et al. 2020; Crichton et al. 2022). Likewise, no living “prosimians” or platyrrhines regularly consume vertebrate prey (some catarrhines such as the cercopithecoid *Papio* and the hominids *Pan* and *Homo* do eat vertebrates, but it is typically not the predominant diet, and we have not included catarrhines in our primate sample). To our knowledge, no fossil primate or primate relative has been suggested to have been a specialised faunivore, but there seems to be no a priori reason why primates and their relatives could not have evolved to fill this dietary niche. Dental topography in combination with discriminant analysis (e.g., QDA) is a powerful tool for predicting diet based on tooth morphology, but taxa cannot be assigned to a particular diet if this diet is not present among the discriminant training sample. Given that our combined marsupial and primate comparative sample still results in relatively high dietary prediction accuracy, we conclude that it is suitable for predicting the diets of extinct metatherians, primates and fossil relatives, and potentially other more distantly related mammaliaform clades, including diets not represented among living marsupials (e.g., hard-object feeding) or primates (e.g., faunivory). The dataset and methods presented here should therefore be valuable tools for future studies that attempt to infer the palaeoecology of fossil mammals.

## CONCLUSIONS

DTMs capture marsupial diets moderately well, with ariaDNE, RFI, the 2D metric TriTaHI, and lnOA of marsupial m2s and m3s combined resulting in 69.2% dietary predictive accuracy. The DTMs used in this study are less successful at predicting marsupial diets than they are for primate diets, with on average > 10% higher classification accuracy for primates than for marsupials for a given selection of DTMs used. The 2D metric TriTaHI, however, has strong dietary predictive power for marsupials when used in isolation, whereas its dietary predictive power is 10% lower for primates, and it loses most of its predictive power when marsupial and primate samples are combined. Compared to primates, marsupial folivores and insectivores exhibit a considerably larger range in ariaDNE and RFI values, overlapping with other dietary categories and increasing the error of the discriminant function analyses. Using a cross-validation approach on the marsupial and primate samples, only three DTMs were needed to reach the highest classification accuracy: ariaDNE, RFI and lnOA. With these DTMs, a moderately accurate dietary prediction of 65% (when using the leave-one-out approach and six dietary categories) or an average of 66% (when using the cross-validation approach with four or three dietary categories) is reached. We conclude that our combined comparative sample allows diets of extinct marsupials and close relatives (non-marsupial metatherians) and extinct primates and close fossil relatives (“plesiadapiforms”) to be predicted with reasonable confidence.

## DATA AVAILABILITY STATEMENT

The datasets generated during and/or analysed during the current study are included in this published article and its supplementary files. Processed digital surface meshes of the marsupial and “prosimian” teeth analysed in this article are available on the Morphosource database as MorphoSource project ID: 000759532, and those of the platyrrhine sample as part of MorphoSource project ID: 000471738.

## ACKNOWLEDGEMENTS

We thank the following researchers for generously sharing their data on MorphoSource that were used in this study: R. Benson, D. Fusco, A. Harrington, J. Ivory-Church, T. Macrini, J. Maisano, E. Mein, J. Ritsevski, J. Rodgers, T. Rowe, M. Silcox, B. Sulbaran, V. Weisbecker, J. van Zoelen. Their funding sources are listed in the Online Resource 2: Table S8. We thank E.L. Fulwood for code for calculating ariaDNE CV, and we thank R. Engelman for helpful discussion on marsupial dental topography and processing pipelines for dental topographic analysis.

## CONTRIBUTIONS

D.V., R.M.D.B., D.A.C. conceived the study. Data processing was done by A.G., D.V., R.D.M.B., with assistance from G.B., N.B., A.C., E.G., E.S. Analyses were performed by D.V. with assistance from A.G. The first draft of the manuscript was written by D.V. and R.M.D.B. and all authors commented on later versions of the manuscript. All authors read and approved the final manuscript.

## ETHICS DECLARATION

D.A.C. is the Editor-in-Chief of the Journal of Mammalian Evolution and R.M.D.B. is an Associate Editor for the Journal of Mammalian Evolution, but neither were involved in the evaluation of this manuscript. The rest of the authors declare no competing interests.

