## supplementary information for "Dental topography and diet in marsupials and comparisons with primates"

**Journal of Mammalian Evolution: Dental topography and diet in marsupials and comparisons with primates**

**Online Resource 2**

**Content** **page**

Table S1: Paired t-test/Wilcoxon signed-rank test *p*-values of m2 vs. m3 marsupial data 1
Full results of paired t-test (*t*-values, df, *p*-values) of m2-only vs. m3-only marsupial data 2
ANOVA & Tukey HSD or Kurskal-Wallis & Dunn test results of marsupial data 3
m2-only 3
m3-only 5
total marsupial sample (m2 & m3) 7
Fig. S1. PCA plot of marsupial sample for ariaDNE, RFI, lnOA, TriTaHI, and OPCR 12
Fig. S2. PCA plots for marsupial m2-only sample and m3-only sample 13
Full results of Welch t-test/Wilcoxon rank-sum test of marsupial versus primate specimen data 15
Fig. S3. Boxplots of marsupial and primate primate data with individual data points 16
Fig. S4. Marsupial insectivore molar shape variation 17
Table S2. Leave-one-out QDAs results based on <u>specimen values</u> vs. [*species averages*] 18
QDA results: combined sample classification accuracy per dietary category 19
Table S3. Exact same marsupial sample: m2-only versus m3-only versus m2&m3 comparison 21
Table S4. QDA results (marsupial = training set, primates = test set) of <u>specimen values</u> vs. [*species averages*] 22
Table S5. QDA category specific results (marsupials m2 = training set, primates = test set) 23
Table S6. QDA category specific results (marsupials m3 = training set, primates = test set) 23
Table S7. QDA results (primates = training set, marsupials = test set) of <u>specimen values</u> vs. [*species averages*] 24
Fig. S5. Sensitivity of curvature metrics to material. 26
Fig. S6. 2D-visualisation of QDA for <u>marsupial</u> (m2&m3, m2-only, m3-only) 27
Fig. S7. 2D-visualisation of QDA for <u>primate</u> 28
Fig. S8. 2D-visualisation of QDA for <u>marsupial (m2 & m3) and primate</u> 29
Table S8. MorphoSource data information and funding sources 30
Fig. S9. PCA plot of total sample (Fig. 7) with primate data labeled 33
Justification of diets 34

**Table S1.** The *p* values of paired t-test or, if one or both groups had non-normally distributed data, <u>Wilcoxon signed-rank test (underlined)</u> of m2-only vs. m3-only. Om = omnivore; Fr-In = frugivore-insectivore; In = insectivore; Fau = faunivore; and Fo = folivore. Significant ($p < 0.05$) results are **bolded**. Frugivores were excluded due to small sample size.

| Metric | Om | Fr-In | In | Fau | Fo |
|---|---|---|---|---|---|
| ariaDNE | 0.871 | 0.158 | 0.196 | 0.096 | **0.014** |
| RFI | 0.495 | 0.938 | 0.847 | 0.146 | 0.255 |
| ln(OA) | 0.890 | 0.560 | **0.041** | **0.007** | 0.424 |
| Trigonid-Talonid RI | 0.263 | <u>0.125</u> | <u>0.148</u> | 0.189 | **0.017** |
| OPCR | 0.994 | 0.551 | 0.359 | 0.647 | 0.781 |
| ariaDNE CV | 0.531 | 0.577 | <u>0.365</u> | <u>1.0</u> | <u>0.119</u> |

Full results of paired t-test (*t*-values, df, and *p*-values) or, if one or both groups had non-normally distributed data, <u>Wilcoxon signed-rank test</u> (*V* values and *p*-values, underlined) of m2-only vs. m3-only marsupial datasets. Significant *p*-values (<0.05) are bolded.

**ariaDNE**
Folivory: t = -2.8282, df = 13, p-value = **0.01424**
Insectivory: t = -1.3854, df = 10, p-value = 0.1961
Frugivory-Insectivory: t = 1.8721, df = 3, p-value = 0.1579
Faunivory: t = -2.164, df = 4, p-value = 0.09645
Omnivory: t = -0.20535, df = 1, p-value = 0.8711
**RFI**
Folivory: t = -1.1915, df = 13, p-value = 0.2547
Insectivory: t = -0.19788, df = 10, p-value = 0.8471
Frugivory-Insectivory: t = -0.083983, df = 3, p-value = 0.9384
Faunivory: t = -1.803, df = 4, p-value = 0.1457
Omnivory: t = 1.0153, df = 1, p-value = 0.4952
**lnOA**
Folivory: t = -0.82613, df = 13, p-value = 0.4236
Insectivory: t = 2.3499, df = 10, p-value = **0.04065**
Frugivory-Insectivory: t = 0.65281, df = 3, p-value = 0.5604
Faunivory: t = -5.1398, df = 4, p-value = **0.006792**
Omnivory: t = 0.17385, df = 1, p-value = 0.8904
**TriTaHI**
Folivory: t = -2.7194, df = 13, p-value = **0.01753**
Insectivory: <u>V = 16, p-value = 0.1475</u>
Frugivory-Insectivory: <u>V = 0, p-value = 0.125</u>
Faunivory: t = -1.5806, df = 4, p-value = 0.1891
Omnivory: t = -2.2848, df = 1, p-value = 0.2626
**OPCR**
Folivory: t = -0.28446, df = 13, p-value = 0.7805
Insectivory: t = -0.96208, df = 10, p-value = 0.3587
Frugivory-Insectivory: t = 0.67002, df = 3, p-value = 0.5508
Faunivory: t = -0.49395, df = 4, p-value = 0.6472
Omnivory: t = 0.0091463, df = 1, p-value = 0.9942
**ariaDNE CV**
<u>Folivory: V = 78, p-value = 0.1189</u>
<u>Insectivory: V = 44, p-value = 0.3652</u>
Frugivory-Insectivory: t = 0.62363, df = 3, p-value = 0.5771
<u>Faunivory: V = 8, p-value = 1</u>
Omnivory: t = -0.90833, df = 1, p-value = 0.5306

ANOVA & Tukey HSD or Kurskal-Wallis & Dunn test results of marsupial data (m2-only, m3-only, m2&m3 combined).
As mentioned in the main text, the frugivore category was dismissed from these results as the sample size was only 2 (one marsupial m2, one marsupial m3).

### ***Marsupial m2 only dataset***

**ariaDNE (ANOVA & Tukey HSD)**

ANOVA:

| | Df | Sum Sq | Mean Sq | F value | Pr(>F) |
|---|---|---|---|---|---|
| diet | 5 | 0.001426 | 0.0002852 | 2.713 | **0.0357** * |
| Residuals | 35 | 0.003679 | 0.0001051 | | |

Shapiro-Wilk normality test:
data: ariadne_aov$residuals
W = 0.97714, p-value = 0.569

Tukey HSD:

| | diff | lwr | upr | p adj |
|---|---|---|---|---|
| Folivory-Faunivory | -0.0074503701 | -0.023166406 | 0.008265666 | 0.7099012 |
| Frugivory-Faunivory | -0.0273579082 | -0.061198017 | 0.006482201 | 0.1718424 |
| Frugivory-Insectivory-Faunivory | 0.0053114614 | -0.014226135 | 0.024849057 | 0.9619973 |
| Insectivory-Faunivory | -0.0003943466 | -0.017056064 | 0.016267371 | 0.9999997 |
| Omnivory-Faunivory | 0.0009984318 | -0.024847378 | 0.026844242 | 0.9999967 |
| Frugivory-Folivory | -0.0199075381 | -0.051694786 | 0.011879709 | 0.4265629 |
| Frugivory-Insectivory-Folivory | 0.0127618315 | -0.002954204 | 0.028477867 | 0.1683197 |
| Insectivory-Folivory | 0.0070560236 | -0.004897594 | 0.019009641 | 0.4918889 |
| Omnivory-Folivory | 0.0084488019 | -0.014644098 | 0.031541702 | 0.8770473 |
| Frugivory-Insectivory-Frugivory | 0.0326693696 | -0.001170739 | 0.066509479 | 0.0635851 |
| Insectivory-Frugivory | 0.0269635616 | -0.005301716 | 0.059228840 | 0.1464246 |
| Omnivory-Frugivory | 0.0283563400 | -0.009478052 | 0.066190732 | 0.2381463 |
| Insectivory-Frugivory-Insectivory | -0.0057058080 | -0.022367526 | 0.010955910 | 0.9039144 |
| Omnivory-Frugivory-Insectivory | -0.0043130296 | -0.030158840 | 0.021532781 | 0.9957393 |
| Omnivory-Insectivory | 0.0013927784 | -0.022353823 | 0.025139380 | 0.9999738 |

**OPCR (ANOVA)**

ANOVA:

| | Df | Sum Sq | Mean Sq | F value | Pr(>F) |
|---|---|---|---|---|---|
| diet | 5 | 1965 | 393.1 | 0.771 | 0.577 |
| Residuals | 35 | 17844 | 509.8 | | |

Shapiro-Wilk normality test
data: opcr_aov$residuals
W = 0.95058, p-value = 0.07334

**RFI (ANOVA)**

ANOVA:

| | Df | Sum Sq | Mean Sq | F value | Pr(>F) |
|---|---|---|---|---|---|
| diet | 5 | 0.07337 | 0.014674 | 1.99 | 0.104 |

| | | | |
|---|---|---|---|
| Residuals | 35 | 0.25808 | 0.007374 |

Shapiro-Wilk normality test:
data: rfi_aov$residuals
W = 0.97955, p-value = 0.6583

**ariaDNE CV (ANOVA)**
ANOVA**:**

| | Df | Sum Sq | Mean Sq | F value | Pr(>F) |
|---|---|---|---|---|---|
| diet | 5 | 0.0298 | 0.005954 | 0.601 | 0.7 |
| Residuals | 35 | 0.3469 | 0.009911 | | |

Shapiro-Wilk normality test:
data: ariadneCV_aov$residuals
W = 0.954, p-value = 0.09677

**lnOA (ANOVA & Tukey HSD)**
ANOVA:

| | Df | Sum Sq | Mean Sq | F value | Pr(>F) |
|---|---|---|---|---|---|
| diet | 5 | 25.93 | 5.187 | 14.65 | **9.29e-08** *** |
| Residuals | 35 | 12.39 | 0.354 | | |

Shapiro-Wilk normality test:
data: lnOA_aov$residuals
W = 0.95485, p-value = 0.1037

Tukey HSD:

| | diff | lwr | upr | p adj |
|---|---|---|---|---|
| Folivory-Faunivory | 0.63879984 | -0.2734506 | 1.551050266 | 0.3057310 |
| Frugivory-Faunivory | -0.25858769 | -2.2228651 | 1.705689701 | 0.9986188 |
| Frugivory-Insectivory-Faunivory | -1.14364528 | -2.2777214 | -0.009569192 | **0.0471049** |
| Insectivory-Faunivory | -1.12041676 | -2.0875601 | -0.153273427 | **0.0153714** |
| Omnivory-Faunivory | -0.20698404 | -1.7072257 | 1.293257601 | 0.9982705 |
| Frugivory-Folivory | -0.89738754 | -2.7425049 | 0.947729792 | 0.6875989 |
| Frugivory-Insectivory-Folivory | -1.78244512 | -2.6946955 | -0.870194695 | **0.0000153** |
| Insectivory-Folivory | -1.75921660 | -2.4530743 | -1.065358909 | **0.0000001** |
| Omnivory-Folivory | -0.84578388 | -2.1862305 | 0.494662779 | 0.4182982 |
| Frugivory-Insectivory-Frugivory | -0.88505758 | -2.8493350 | 1.079219814 | 0.7510983 |
| Insectivory-Frugivory | -0.86182907 | -2.7346941 | 1.011035946 | 0.7345244 |
| Omnivory-Frugivory | 0.05160365 | -2.1445252 | 2.247732545 | 0.9999997 |
| Insectivory-Frugivory-Insectivory | 0.02322852 | -0.9439148 | 0.990371849 | 0.9999997 |
| Omnivory-Frugivory-Insectivory | 0.93666123 | -0.5635804 | 2.436902876 | 0.4299856 |
| Omnivory-Insectivory | 0.91343272 | -0.4649586 | 2.291824045 | 0.3644611 |

**TriTaHI (ANOVA & Tukey HSD)**
ANOVA:

| | Df | Sum Sq | Mean Sq | F value | Pr(>F) |
|---|---|---|---|---|---|
| diet | 5 | 0.4683 | 0.09366 | 7.142 | **0.000107 ***** |
| Residuals | 35 | 0.4590 | 0.01311 | | |

Shapiro-Wilk normality test:
data: TriTaHI_aov$residuals
W = 0.95338, p-value = 0.09203

| | diff | lwr | upr | p adj |
|---|---|---|---|---|
| Folivory-Faunivory | -0.28966193 | -0.46521157 | -0.11411229 | **0.0002370** |
| Frugivory-Faunivory | -0.20726521 | -0.58526250 | 0.17073208 | 0.5710047 |
| Frugivory-Insectivory-Faunivory | -0.07236906 | -0.29060590 | 0.14586778 | 0.9150657 |
| Insectivory-Faunivory | -0.11620798 | -0.30232098 | 0.06990502 | 0.4298890 |
| Omnivory-Faunivory | -0.17285874 | -0.46155893 | 0.11584146 | 0.4762449 |
| Frugivory-Folivory | 0.08239672 | -0.27266991 | 0.43746334 | 0.9808085 |
| Frugivory-Insectivory-Folivory | 0.21729287 | 0.04174323 | 0.39284251 | **0.0081833** |
| Insectivory-Folivory | 0.17345395 | 0.03993089 | 0.30697701 | **0.0049549** |
| Omnivory-Folivory | 0.11680319 | -0.14114673 | 0.37475312 | 0.7472655 |
| Frugivory-Insectivory-Frugivory | 0.13489615 | -0.24310114 | 0.51289344 | 0.8878121 |
| Insectivory-Frugivory | 0.09105723 | -0.26934904 | 0.45146351 | 0.9721899 |
| Omnivory-Frugivory | 0.03440647 | -0.38820734 | 0.45702029 | 0.9998672 |
| Insectivory-Frugivory-Insectivory | -0.04383892 | -0.22995192 | 0.14227408 | 0.9795044 |
| Omnivory-Frugivory-Insectivory | -0.10048968 | -0.38918987 | 0.18821052 | 0.8978041 |
| Omnivory-Insectivory | -0.05665076 | -0.32190259 | 0.20860108 | 0.9867433 |

### ***Marsupial m3 only dataset***

**ariaDNE (ANOVA)**

ANOVA:

| | Df | Sum Sq | Mean Sq | F value | Pr(>F) |
|---|---|---|---|---|---|
| diet | 5 | 0.001258 | 0.0002516 | 2.132 | 0.0853 |
| Residuals | 34 | 0.004012 | 0.0001180 | | |

Shapiro-Wilk normality test
data: ariadne_aov$residuals
W = 0.97215, p-value = 0.4198

**OPCR (Kruskal-Wallis)**

ANOVA:

| | Df | Sum Sq | Mean Sq | F value | Pr(>F) |
|---|---|---|---|---|---|
| diet | 5 | 3668 | 733.6 | 1.532 | 0.206 |
| Residuals | 34 | 16285 | 479.0 | | |

Shapiro-Wilk normality test
data: opcr_aov$residuals
W = 0.92327, p-value = **0.009772**

Kruskal-Wallis rank sum test
data: OPCR by diet
Kruskal-Wallis chi-squared = 7.4396, df = 5, p-value = 0.1899

**RFI (ANOVA)**

ANOVA:

| | Df | Sum Sq | Mean Sq | F value | Pr(>F) |
|---|---|---|---|---|---|
| diet | 5 | 0.08672 | 0.017345 | 2.051 | 0.0961 |
| Residuals | 34 | 0.28753 | 0.008457 | | |

Shapiro-Wilk normality test
data:  rfi_aov$residuals
W = 0.98288, p-value = 0.7943

**ariaDNE CV (Kruskal-Wallis)**
ANOVA:

| | Df | Sum Sq | Mean Sq | F value | Pr(>F) |
|---|---|---|---|---|---|
| diet | 5 | 0.1225 | 0.024491 | 2.572 | **0.0446 *** |
| Residuals | 34 | 0.3237 | 0.009522 | | |

Shapiro-Wilk normality test:
data:  ariadneCV_aov$residuals
W = 0.93972, p-value = **0.03384**

Kruskal-Wallis rank sum test:
data:  ariaDNE_CV by diet
Kruskal-Wallis chi-squared = 7.6309, df = 5, p-value = 0.1778

**lnOA (ANOVA & TukeyHSD)**
ANOVA:

| | Df | Sum Sq | Mean Sq | F value | Pr(>F) |
|---|---|---|---|---|---|
| diet | 5 | 26.89 | 5.377 | 15.49 | **6.06e-08 ***** |
| Residuals | 34 | 11.80 | 0.347 | | |

Shapiro-Wilk normality test:
data:  lnOA_aov$residuals
W = 0.95573, p-value = 0.1195

Tukey HSD:

| | diff | lwr | upr | p adj |
|---|---|---|---|---|
| Folivory-Faunivory | 0.5218933 | -0.3371654 | 1.3809519 | 0.4587785 |
| Frugivory-Faunivory | -0.5169776 | -2.4378912 | 1.4039359 | 0.9632658 |
| Frugivory-Insectivory-Faunivory | -1.4069308 | -2.5548962 | -0.2589654 | **0.0091083** |
| Insectivory-Faunivory | -1.3153835 | -2.2179660 | -0.4128010 | **0.0013241** |
| Omnivory-Faunivory | -0.3211818 | -1.5787149 | 0.9363513 | 0.9706033 |
| Frugivory-Folivory | -1.0388709 | -2.8756156 | 0.7978738 | 0.5366929 |
| Frugivory-Insectivory-Folivory | -1.9288240 | -2.9295971 | -0.9280510 | **0.0000208** |
| Insectivory-Folivory | -1.8372767 | -2.5432352 | -1.1313183 | **0.0000001** |
| Omnivory-Folivory | -0.8430751 | -1.9678469 | 0.2816967 | 0.2372469 |
| Frugivory-Insectivory-Frugivory | -0.8899531 | -2.8782876 | 1.0983813 | 0.7548982 |
| Insectivory-Frugivory | -0.7984058 | -2.6559054 | 1.0590937 | 0.7843843 |
| Omnivory-Frugivory | 0.1957958 | -1.8577472 | 2.2493388 | 0.9997084 |
| Insectivory-Frugivory-Insectivory | 0.0915473 | -0.9468265 | 1.1299211 | 0.9998014 |
| Omnivory-Frugivory-Insectivory | 1.0857490 | -0.2725421 | 2.4440400 | 0.1804676 |
| Omnivory-Insectivory | 0.9942017 | -0.1641528 | 2.1525561 | 0.1273301 |

**TriTaHI (Kruskal-Wallis & Dunn test)**

ANOVA:

| | Df | Sum Sq | Mean Sq | F value | Pr(>F) |
|---|---|---|---|---|---|
| diet | 5 | 0.5551 | 0.11101 | 8.49 | **2.7e-05** *** |
| Residuals | 34 | 0.4446 | 0.01308 | | |

Shapiro-Wilk normality test
data:  TriTaHI_aov$residuals
W = 0.94459, p-value = **0.04948**

Kruskal-Wallis rank sum test:
data:  TriTaHI by diet
Kruskal-Wallis chi-squared = 18.783, df = 5, p-value = **0.002109**

Comparison of x by group (Bonferroni)

| Col Mean-<br>Row Mean | Faunivor | Folivory | Frugivor | Fru-Ins | Insectiv |
|---|---|---|---|---|---|
| Folivory | 3.736480<br>**0.0014*** | | | | |
| Frugivor | 0.884339<br>1.0000 | -0.822714<br>1.0000 | | | |
| Fru-Ins | 0.353380<br>1.0000 | -2.802021<br>**0.0381** | -0.650328<br>1.0000 | | |
| Insectiv | 1.590963<br>0.8371 | -2.512725<br>0.0899 | -0.141460<br>1.0000 | 0.992231<br>1.0000 | |
| Omnivory | 0.987935<br>1.0000 | -1.749239<br>0.6019 | -0.222239<br>1.0000 | 0.615989<br>1.0000 | -0.167146<br>1.0000 |

***Total marsupial dataset (m2 & m3)***

**ariaDNE (ANOVA and Tukey HSD)**

ANOVA:

| | Df | Sum Sq | Mean Sq | F value | Pr(>F) |
|---|---|---|---|---|---|
| diet | 5 | 0.002656 | 0.0005311 | 5.098 | **0.000446** *** |
| Residuals | 75 | 0.007814 | 0.0001042 | | |

Shapiro-Wilk normality test:
data: ariadne_aov$residuals
W = 0.97986, p-value = 0.2321

Tukey HSD:

| | diff | lwr | upr | p adj |
|---|---|---|---|---|
| Folivory-Faunivory | -0.008823381 | -0.0192570993 | 0.001610338 | 0.1453642 |
| Frugivory-Faunivory | -0.029362962 | -0.0523105239 | -0.006415400 | **0.0046177** |
| Frugivory-Insectivory-Faunivory | 0.002196431 | -0.0112211312 | 0.015613993 | 0.9967657 |
| Insectivory-Faunivory | -0.002333852 | -0.0133575038 | 0.008689801 | 0.9892921 |
| Omnivory-Faunivory | -0.000196325 | -0.0162973997 | 0.015904750 | 1.0000000 |
| Frugivory-Folivory | -0.020539581 | -0.0422979175 | 0.001218756 | 0.0753665 |
| Frugivory-Insectivory-Folivory | 0.011019812 | -0.0002436502 | 0.022283274 | 0.0587595 |
| Insectivory-Folivory | 0.006489529 | -0.0017782099 | 0.014757268 | 0.2088132 |
| Omnivory-Folivory | 0.008627056 | -0.0057284034 | 0.022982515 | 0.4989767 |
| Frugivory-Insectivory-Frugivory | 0.031559393 | 0.0082228614 | 0.054895924 | **0.0022973** |
| Insectivory-Frugivory | 0.027029110 | 0.0049818058 | 0.049076415 | **0.0075784** |
| Omnivory-Frugivory | 0.029166637 | 0.0041905015 | 0.054142772 | **0.0127252** |
| Insectivory-Frugivory-Insectivory | -0.004530283 | -0.0163423107 | 0.007281745 | 0.8708792 |
| Omnivory-Frugivory-Insectivory | -0.002392756 | -0.0190435129 | 0.014258001 | 0.9982615 |
| Omnivory-Insectivory | 0.002137527 | -0.0126522550 | 0.016927308 | 0.9982132 |

**OPCR (Kruskal-Wallis)**

ANOVA:

| | Df | Sum Sq | Mean Sq | F value | Pr(>F) |
|---|---|---|---|---|---|
| diet | 5 | 5357 | 1071.4 | 2.336 | **0.0501** |
| Residuals | 75 | 34406 | 458.7 | | |

Shapiro-Wilk normality test:
data: opcr_aov$residuals
W = 0.95264, p-value = **0.004566**

Kruskal-Wallis rank sum test:
data: OPCR by diet
Kruskal-Wallis chi-squared = 8.9497, df = 5, p-value = 0.1111

**RFI (ANOVA & Tukey HSD)**

ANOVA:

| | Df | Sum Sq | Mean Sq | F value | Pr(>F) |
|---|---|---|---|---|---|
| diet | 5 | 0.1600 | 0.0320 | 4.386 | **0.00148** ** |
| Residuals | 75 | 0.5471 | 0.0073 | | |

Shapiro-Wilk normality test:
data: rfi_aov$residuals
W = 0.98783, p-value = 0.6449

Tukey HSD:

| | diff | lwr | upr | p adj |
|---|---|---|---|---|
| Folivory-Faunivory | -0.10877431 | -0.196082801 | -0.02146582 | **0.0063175** |
| Frugivory-Faunivory | -0.13554736 | -0.327570650 | 0.05647593 | 0.3169977 |
| Frugivory-Insectivory-Faunivory | -0.05029902 | -0.162576064 | 0.06197803 | 0.7785158 |
| Insectivory-Faunivory | -0.09609924 | -0.188344241 | -0.00385423 | **0.0362358** |
| Omnivory-Faunivory | 0.01373361 | -0.120998847 | 0.14846606 | 0.9996714 |
| Frugivory-Folivory | -0.02677305 | -0.208844997 | 0.15529890 | 0.9980591 |
| Frugivory-Insectivory-Folivory | 0.05847529 | -0.035776420 | 0.15272701 | 0.4628608 |
| Insectivory-Folivory | 0.01267508 | -0.056508677 | 0.08185883 | 0.9945080 |
| Omnivory-Folivory | 0.12250792 | 0.002382631 | 0.24263321 | **0.0429838** |
| Frugivory-Insectivory-Frugivory | 0.08524834 | -0.110029815 | 0.28052650 | 0.7966348 |
| Insectivory-Frugivory | 0.03944812 | -0.145041887 | 0.22393813 | 0.9887973 |
| Omnivory-Frugivory | 0.14928097 | -0.059717256 | 0.35827919 | 0.3041847 |
| Insectivory-Frugivory-Insectivory | -0.04580022 | -0.144642285 | 0.05304185 | 0.7531623 |
| Omnivory-Frugivory-Insectivory | 0.06403263 | -0.075299522 | 0.20336477 | 0.7595274 |
| Omnivory-Insectivory | 0.10983284 | -0.013926819 | 0.23359250 | 0.1110764 |

**ariaDNE CV (Kruskal-Wallis)**

ANOVA:

| | Df | Sum Sq | Mean Sq | F value | Pr(>F) |
|---|---|---|---|---|---|
| diet | 5 | 0.1344 | 0.026871 | 2.927 | 0.0181 * |
| Residuals | 75 | 0.6886 | 0.009181 | | |

Shapiro-Wilk normality test
data: dne_aov$residuals
W = 0.96201, p-value = **0.01685**

Kruskal-Wallis rank sum test
data: ariaDNE_CV by diet
Kruskal-Wallis chi-squared = 9.9473, df = 5, p-value = 0.07674

**lnOA (Kruskal-Wallis & Dunn test)**

ANOVA:

| | Df | Sum Sq | Mean Sq | F value | Pr(>F) |
|---|---|---|---|---|---|
| diet | 5 | 52.72 | 10.544 | 32.49 | **<2e-16 ***** |
| Residuals | 75 | 24.34 | 0.325 | | |

Shapiro-Wilk normality test:
data: lnOA_aov$residuals
W = 0.95862, p-value = **0.01041**

Kruskal-Wallis rank sum test:
data: lnOA by diet
Kruskal-Wallis chi-squared = 55.914, df = 5, p-value = **8.465e-11**

Dunn test:

Comparison of x by group (Bonferroni)

| Col Mean- Row Mean | Faunivor | Folivory | Frugivor | Frugi-Ins | Insectiv |
|---|---|---|---|---|---|
| Folivory | -1.578544 | | | | |
| | 0.8583 | | | | |
| Frugivor | 0.683639 | 1.477959 | | | |
| | 1.0000 | 1.0000 | | | |
| Frugivor-Ins | 2.965993 | 4.995487 | 1.033081 | | |
| | **0.0226*** | **0.0000*** | 1.0000 | | |
| Insectiv | 3.521148 | 6.686954 | 1.049019 | -0.083007 | |
| | **0.0032*** | **0.0000*** | 1.0000 | 1.0000 | |
| Omnivory | 0.501496 | 1.709782 | -0.304819 | -1.905124 | -2.078548 |
| | 1.0000 | 0.6548 | 1.0000 | 0.4257 | 0.2824 |

**TriTaHI (Kruskal-Wallis & Dunn test)**

ANOVA:

| | Df | Sum Sq | Mean Sq | F value | Pr(>F) |
|---|---|---|---|---|---|
| diet | 5 | 1.0269 | 0.20538 | 16.42 | **6.45e-11 *** ** |
| Residuals | 75 | 0.9381 | 0.01251 | | |

Shapiro-Wilk normality test:

data: TriTaHI_aov$residuals

W = 0.95913, p-value = **0.01118**

Kruskal-Wallis rank sum test:

data: TriTaHI by diet

Kruskal-Wallis chi-squared = 37.256, df = 5, p-value = **5.323e-07**

Dunn test:

Comparison of x by group (Bonferroni)

| Col Mean- Row Mean | Faunivor | Folivory | Frugivor | Fru-Ins | Insectiv |
|---|---|---|---|---|---|
| Folivory | 5.190643 | | | | |
| | **0.0000*** | | | | |
| Frugivor | 1.032999 | -1.399596 | | | |
| | 1.0000 | 1.0000 | | | |
| Fru-Ins | 0.910335 | -3.723831 | -0.492375 | | |
| | 1.0000 | **0.0015*** | 1.0000 | | |

| | | | | | |
|---|---|---|---|---|---|
| Insectiv | 2.040486 | -3.829837 | -0.054936 | 0.870226 | |
| | 0.3098 | **0.0010*** | 1.0000 | 1.0000 | |
| Omnivory | 1.243711 | -2.377675 | -0.147329 | 0.469084 | -0.166907 |
| | 1.0000 | 0.1307 | 1.0000 | 1.0000 | 1.0000 |

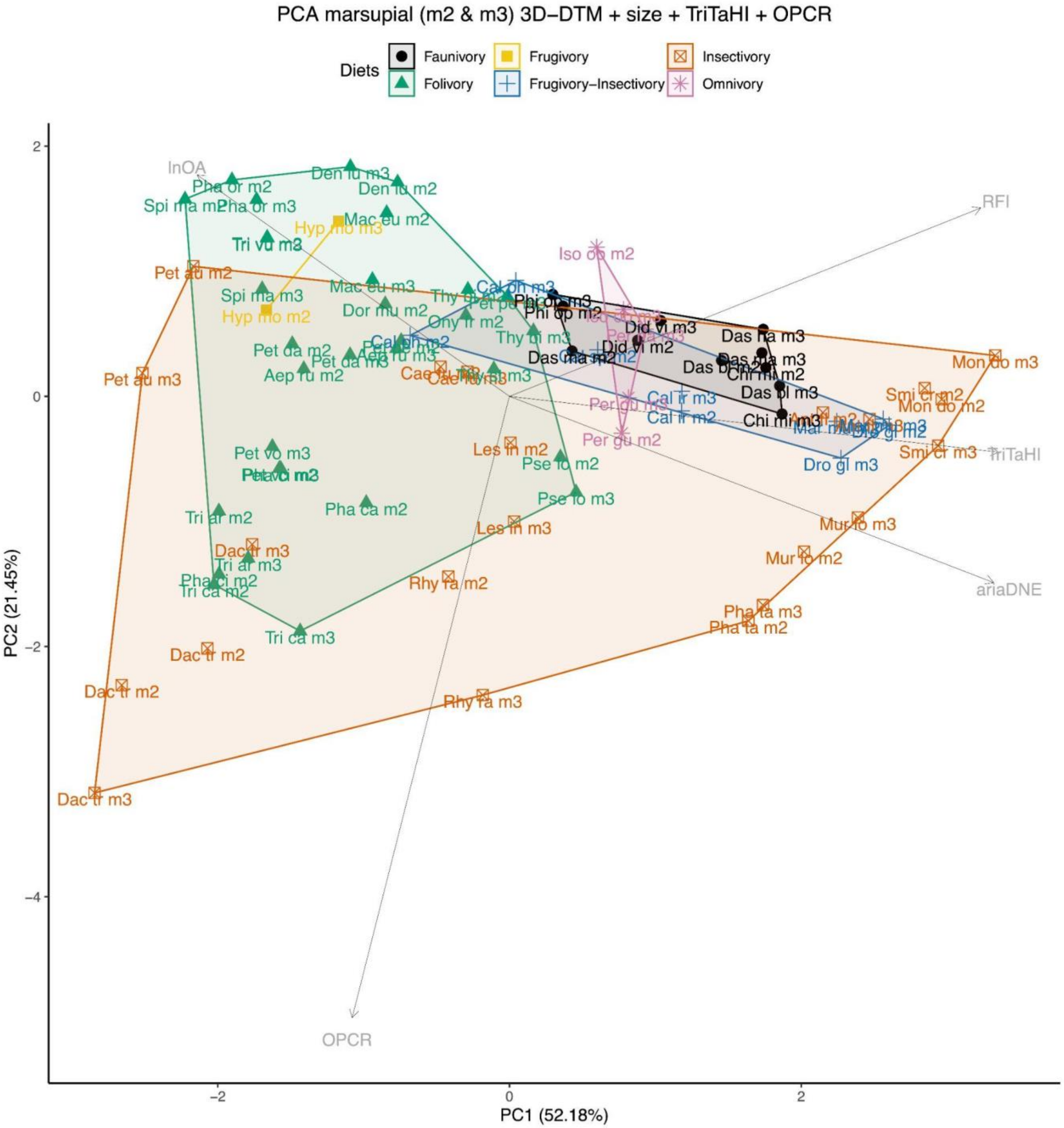

**Fig. S1** Principal Component Analysis of 3D-DTM (= ariaDNE and RFI), lnOA, TriTaHI, and OPCR plot showing PC1 and PC2 scores for total marsupial sample, capturing 73.63% of the variation. Note that the PC loading arrows are not to scale.

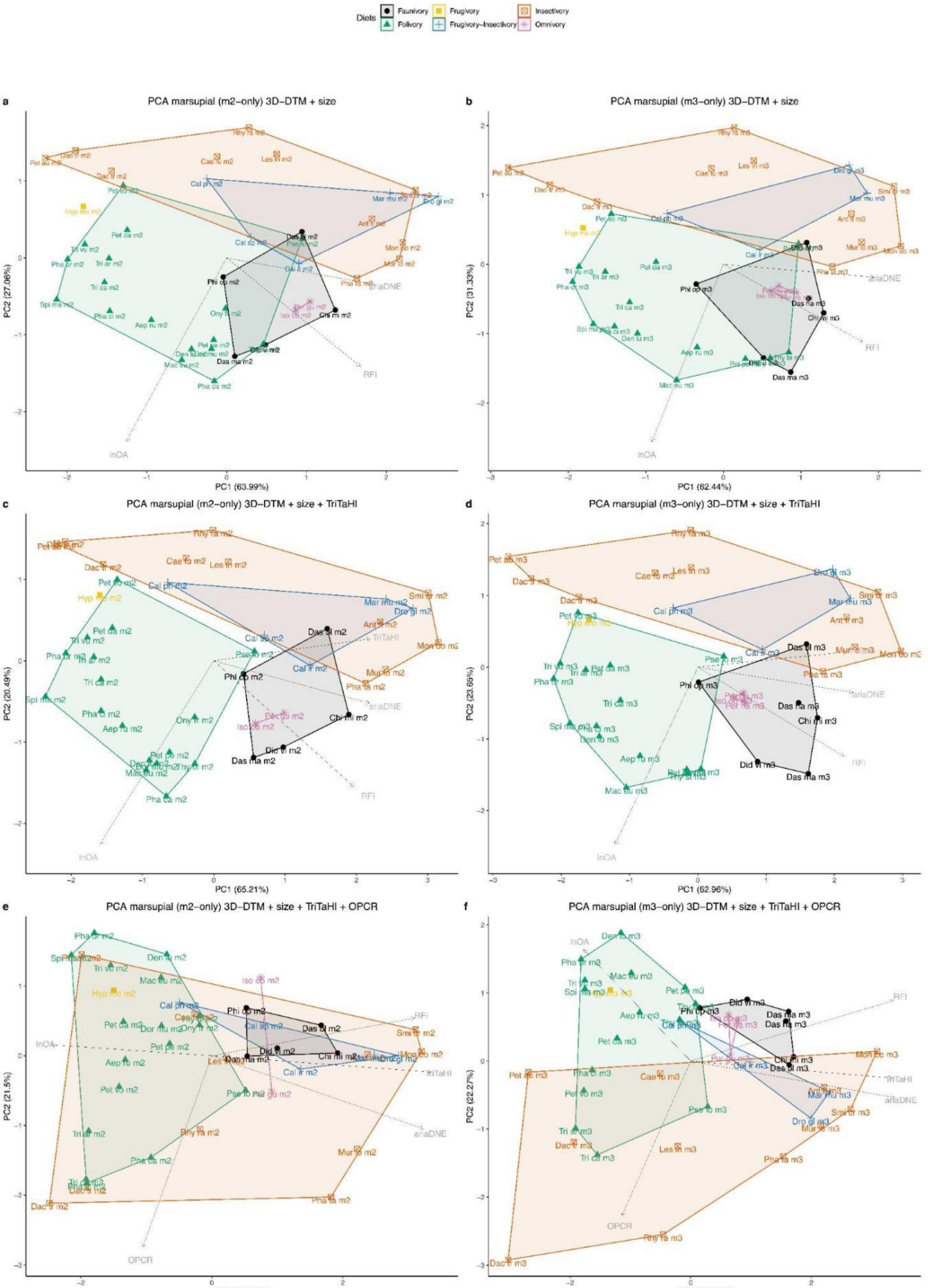
Diets
Faunivory
Folivory
Frugivory
Frugivory–Insectivory
Insectivory
Omnivory
a
PCA marsupial (m2–only) 3D–DTM + size
PC1 (63.99%)
PC2 (27.06%)
b
PCA marsupial (m3–only) 3D–DTM + size
PC1 (62.44%)
PC2 (31.33%)
c
PCA marsupial (m2–only) 3D–DTM + size + TriTaHI
PC1 (65.21%)
PC2 (20.49%)
d
PCA marsupial (m3–only) 3D–DTM + size + TriTaHI
PC1 (62.96%)
PC2 (23.65%)
e
PCA marsupial (m2–only) 3D–DTM + size + TriTaHI + OPCR
PC1 (53.13%)
PC2 (21.5%)
f
PCA marsupial (m3–only) 3D–DTM + size + TriTaHI + OPCR
PC1 (51.44%)
PC2 (22.27%)
lnOA
RFI
ariaDNE
TriTaHI
OPCR

**Fig. S2** Principal Component Analysis plots of marsupial m2-only (a, c, and e) and m3-only samples (b, d, and f) using 3D-DTM (= ariaDNE and RFI), lnOA (a and b), those metrics as well as TriTaHI (c and d), and finally all those metrics plus OPCR (e and f).

Full results of Welch t-test (*t*-values, df, and *p*-values) or, if one or both groups had non-normally distributed data, Wilcoxon signed-rank test (*W*-values and *p*-values, underlined) of marsupial (m2 & m3) vs. primate (platyrrhine & strepsirrhine) specimen-based datasets. Significant *p*-values (<0.05) are bolded. Note that results are listed for the frugivory category, but these were not included in the main manuscript as the marsupial frugivore sample was only *n* = 2.

**ariaDNE**
Folivory: W = 250, p-value = **0.0003203**
Insectivory: W = 52, p-value = **0.003131**
Frugivory-Insectivory: t = 1.9366, df = 8.7774, p-value = 0.08558
Frugivory: t = -22.401, df = 4.5413, p-value = **7.987e-06**
**RFI**
Folivory: W = 419, p-value = 0.2158
Insectivory: t = -1.354, df = 28.498, p-value = 0.1864
Frugivory-Insectivory: t = 2.7299, df = 9.549, p-value = **0.02205**
Frugivory: t = -2.6624, df = 1.392, p-value = 0.1684
**lnOA**
Folivory: W = 313, p-value = **0.007084**
Insectivory: W = 93, p-value = 0.1677
Frugivory-Insectivory: W = 183, p-value = 0.2717
Frugivory: t = -6.4616, df = 4.6025, p-value = **0.001796**
**TriTaHI**
Folivory: t = -8.292, df = 61.996, p-value = **1.235e-11**
Insectivory: W = 122, p-value = 0.7358
Frugivory-Insectivory: W = 426, p-value = **0.0001863**
Frugivory: t = 4.1925, df = 1.0991, p-value = 0.1317
**OPCR**
Folivory: t = -8.9947, df = 60.607, p-value = **9.233e-13**
Insectivory: W = 20, p-value = **9.461e-06**
Frugivory-Insectivory: t = -22.276, df = 51.328, p-value **< 2.2e-16**
Frugivory: t = -10.788, df = 1.5434, p-value = **0.01945**
**ariaDNE CV**
Folivory: W = 2, p-value **< 2.2e-16**
Insectivory: t = -6.6215, df = 24.971, p-value = **6.184e-07**
Frugivory-Insectivory: t = -5.9879, df = 8.59, p-value = **0.0002476**
Frugivory: t = -34.37, df = 3.964, p-value = **4.672e-06**

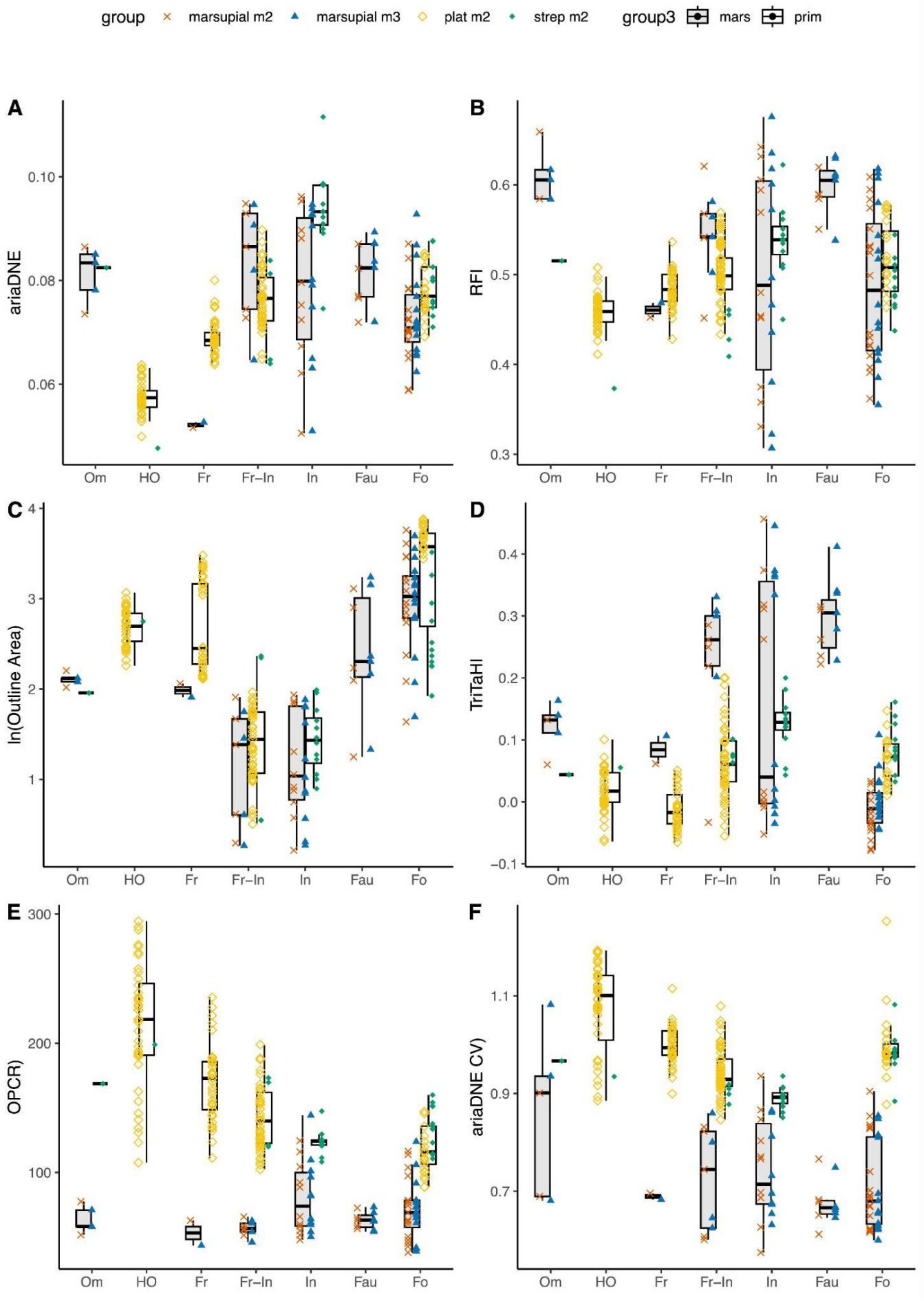

group
marsupial m2
marsupial m3
plat m2
strep m2
group3
mars
prim
A
ariaDNE
B
RFI
C
ln(Outline Area)
D
TriTaHI
E
OPCR)
F
ariaDNE CV)
Om
HO
Fr
Fr−In
In
Fau
Fo

**Fig. S3** Boxplots of marsupial (m2 [orange cross] & m3 [blue triangle]) data in grey and primate (platyrrhine [yellow open diamonds] & strepsirrhine [green filled diamonds]) in white, both displaying specimen data distribution for different diets. All significantly different ($p < 0.05$) pairs are marked by brackets and asterisks. Om = omnivore; HO = hard-object feeder; Fr = frugivore; Fr-In = frugivore-insectivore; In = insectivore; Fau = faunivore; and Fo = folivore. * = $p < 0.05$; ** = $p < 0.01$; and *** = $p < 0.001$. Note that omnivores, hard-object feeders, frugivores, and faunivores were excluded from the t-tests due to small sample sizes.

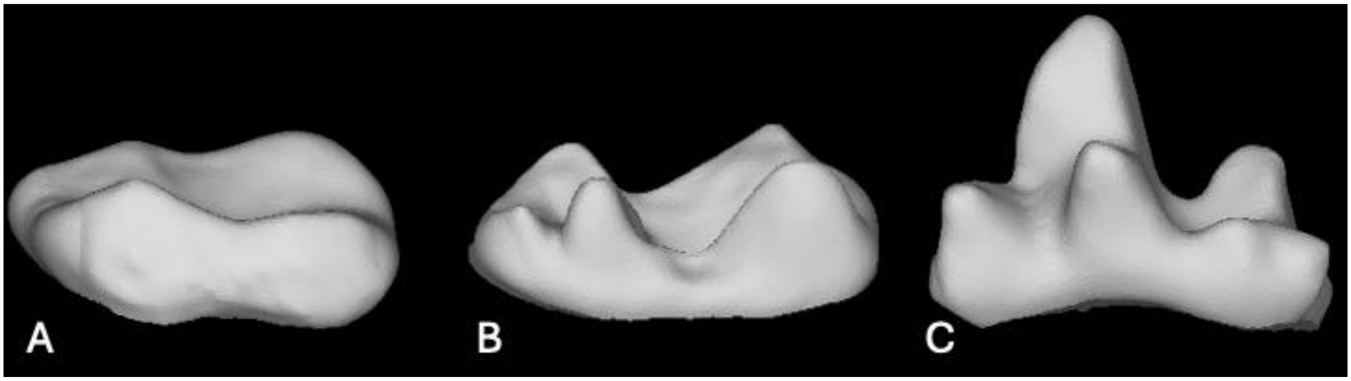

**Fig. S4** Marsupial insectivore molar shape variation. A. *Petaurus australis* QM J8598 m2, member of 'group 1'. B. *Lestoros inca* MVZ116049 m2, member of 'group 2'. C. *Smithopsis crassicaudata* SAMA M12809 m2, member of 'group 3'. See main text for information on grouping. Not to scale.

Table S2. Leave-one-out QDAs accuracy based on specimen values or [*species averages]*.

| Metric | Marsupial m2s | Marsupial m3s | Marsupial all | Platyrrhines [*species averages only 3 diets, hence NA*] | Strepsirrhines [*species averages only 2 diets, hence NA*] | Primates [5] | Marsupial m2s + Primates [6] | Marsupial m3s + Primates [6] | Marsupial all + Primates [6] |
|---|---|---|---|---|---|---|---|---|---|
| RFI+ariaDNE | 34.21% [*40.5%*] | 55.56% [*57.1%*] (59.38%*) [*61.3%**] | 51.35% [*46.2%*] | 70.34% [*NA*] | 71.43% (83.33%*) [*NA*] | 62.64% [*62.7%*] | 55.87% [*47.2%*] | 61.14% [*52.9%*] | 56.4% [*48.4%*] |
| RFI+ariaDNE+ size | 55.26% [*51.4%*] | 69.44% [*68.6%*] (78.13%*) [*77.4%**] | 77.03% [*64.1%*] | 97.24% [*NA*] | 82.14% (95.83%*) [*NA*] | 91.95% [*80.4%*] | 79.34% [*60.7%*] | 81.99% [*63.2%*] | 79.6% [*64.8%*] |
| RFI+ariaDNE+ size+TrigTaHI | 57.89% [*56.8%*] | 87.5%* [*80.6%**] | 79.73% [*69.2%*] | 97.24% [*NA*] | 91.67%* [*NA*] | 91.38% [*74.5%*] | 83.1% [*67.4%*] | 82.46% [*70.1%*] | 81.2% [*69.2%*] |
| RFI+ariaDNE+ size+ariaDNE CV | 52.63% [*51.4%*] | 81.25%* [*80.6%**] | 78.39% [*66.7%*] | 94.48% [*NA*] | 91.67%* [*NA*] | 88.51% [*76.5%*] | 84.04% [*71.9%*] | 86.73% [*80.5%*] | 83.6% [*75.8%*] |
| RFI+ariaDNE+ size+OPCR | 63.16% [*62.2%*] | 78.13%* [*77.4%**] | 79.73% [*61.5%*] | 99.31% [*NA*] | 91.67%* [*NA*] | 93.10% [*74.5%*] | 86.85% [*73.0%*] | 88.15% [*78.2%*] | 87.6% [*79.1%*] |

**Marsupial samples (m2 only, m3 only, and all):** 4 different dietary categories (Folivory, Insectivory, Frugivory-Insectivory, Faunivory), unless marked with an * = Folivory, Insectivory, Faunivory)
**Platyrrhine sample:** 4 dietary categories (Folivory, Frugivory, Hard-Object feeder, Frugivory-Insectivory)
**Strepsirrhine sample:** 3 dietary categories (Folivory, Frugivory-Insectivory, Insectivory), unless makes with a * = Folivory, Insectivory
**Primates sample**: 5 dietary categories (Folivory, Frugivory, Hard-object feeder, Frugivory-Insectivory, Insectivory)
**Mars m2 and/or m3 + primates sample**: 6 dietary categories (Faunivory, Folivory, Frugivory, Frugivory-Insectivory, Hard-object feeder, Insectivory)

**QDA combined sample classification accuracy per dietary category**

**QDA primates + mars m2**

ariaDNE + RFI (6 diets, overall: 47.2%)

| Faunivory | Folivory | Frugivory | Frugivory-Insectivory |
|---|---|---|---|
| 0.8000000 | 0.2758621 | 0.7000000 | 0.3684211 |

| Hard-object feeder | Insectivory |
|---|---|
| 0.8888889 | 0.4705882 |

ariaDNE + RFI + lnOA (6 diets, overall: 60.7%)

| Faunivory | Folivory | Frugivory | Frugivory-Insectivory |
|---|---|---|---|
| 0.2000000 | 0.5862069 | 0.5000000 | 0.7368421 |

| Hard-object feeder | Insectivory |
|---|---|
| 0.8888889 | 0.5294118 |

ariaDNE + RFI + lnOA+ TriTaHI (6 diets, overall: 67.4%)

| Faunivory | Folivory | Frugivory | Frugivory-Insectivory |
|---|---|---|---|
| 0.0000000 | 0.8275862 | 0.7000000 | 0.6315789 |

| Hard-object feeder | Insectivory |
|---|---|
| 0.7777778 | 0.5882353 |

**QDA primates + mars m3**

ariaDNE + RFI (6 diets, overall: 52.9%)

| Faunivory | Folivory | Frugivory | Frugivory-Insectivory |
|---|---|---|---|
| 0.8333333 | 0.3333333 | 0.7000000 | 0.4444444 |

| Hard-object feeder | Insectivory |
|---|---|
| 0.8888889 | 0.5294118 |

ariaDNE + RFI + lnOA (6 diets, overall: 63.2%)

| Faunivory | Folivory | Frugivory | Frugivory-Insectivory |
|---|---|---|---|
| 0.5000000 | 0.5555556 | 0.5000000 | 0.6666667 |

| Hard-object feeder | Insectivory |
|---|---|
| 0.8888889 | 0.7058824 |

ariaDNE + RFI + lnOA+ TriTaHI (6 diets, overall: 70.1%)

| Faunivory | Folivory | Frugivory | Frugivory-Insectivory |
|---|---|---|---|
| 0.3333333 | 0.7777778 | 0.6000000 | 0.6666667 |

| Hard-object feeder | Insectivory |
|---|---|
| 0.8888889 | 0.7058824 |

**QDA primates + mars m2 & m3**

<u>ariaDNE + RFI (*7 diets*, overall: 42.1%)</u>

| Faunivory | Folivory | Frugivory | Frugivory-Insectivory |
|---|---|---|---|
| 0.6666667 | 0.2333333 | 0.7000000 | 0.2631579 |
| Hard-object feeder | Insectivory | Omnivory | |
| 0.8888889 | 0.4705882 | 0.2500000 | |

<u>ariaDNE + RFI (6 diets, overall: 48.4%)</u>

| Faunivory | Folivory | Frugivory | Frugivory-Insectivory |
|---|---|---|---|
| 0.8333333 | 0.2666667 | 0.7000000 | 0.3684211 |
| Hard-object feeder | Insectivory | | |
| 0.8888889 | 0.5294118 | | |

<u>ariaDNE + RFI + lnOA (*7 diets*, overall: 60%)</u>

| Faunivory | Folivory | Frugivory | Frugivory-Insectivory |
|---|---|---|---|
| 0.1666667 | 0.6000000 | 0.5000000 | 0.7368421 |
| Hard-object feeder | Insectivory | Omnivory | |
| 0.8888889 | 0.6470588 | 0.0000000 | |

<u>ariaDNE + RFI + lnOA (6 diets, overall: 64.8%)</u>

| Faunivory | Folivory | Frugivory | Frugivory-Insectivory |
|---|---|---|---|
| 0.5000000 | 0.6000000 | 0.5000000 | 0.7368421 |
| Hard-object feeder | Insectivory | | |
| 0.8888889 | 0.6470588 | | |

<u>ariaDNE + RFI + lnOA+ TriTaHI (6 diets, overall: 69.2%)</u>

| Faunivory | Folivory | Frugivory | Frugivory-Insectivory |
|---|---|---|---|
| 0.3333333 | 0.8333333 | 0.5000000 | 0.6315789 |
| Hard-object feeder | Insectivory | | |
| 0.8888889 | 0.6470588 | | |

For the QDAs including all specimens (marsupial all + primates) with only ariaDNE + RFI, or ariaDNE + RFI + size as the metrics, it sample sizes were large enough to include the Omnivory category as well, resulting in slightly lower accuracies than when this category was excluded (i.e., 42.1% versus 48.4% for ariaDNE and RFI, and 60.0% versus 64.8% for ariaDNE, RFI, and size).

**Table S3.** Exact same sample (36 taxa) m2-only versus m3-only versus m2&m3 comparison (species averages)

| Metric | Marsupial m2-only | Marsupial m3-only | Marsupial m2 & m3 | Primate |
|---|---|---|---|---|
| RFI+ariaDNE | 48.5% | 57.6% | 57.6% | 62.7% |
| RFI+ariaDNE+size | 45.5% | 66.7% | 63.6% | 80.4% |
| RFI+ariaDNE+size+TrigTaHI | 72.4%* | 72.4%* | 72.4%* | 74.5% |

**Marsupial samples (m2 only, m3 only, and all):** 4 different dietary categories (folivory, insectivory, frugivory-insectivory, faunivory), unless marked with a * = folivory, insectivory, faunivory)
Notes: m2 performs better in insectivore and frugivore-insectivore categories for ariaDNE and RFI than the m3 does, and better for insectivore and folivore for ariaDNE, RFI, size.

**Table S4.** QDA results (marsupials = training set, primates = test set) of specimen values vs. [*species averages*]. Training accuracy is calculated using the entire training sample (not leave-one-out). Training set included Faunivory, Folivory, Frugivory-Insectivory, and Insectivory categories. Accuracy of the QDA test was calculated using shared categories only, i.e., Folivory, Insectivory, and, in some cases, Frugivory-Insectivory. Results marked with an * = Frugivore-Insectivory category was excluded as the sample for this category was too small.

| Train sample (train accuracy) | Train accuracy | Variables | Test sample | Test accuracy | Folivory accuracy | Frugivory-Insectivory accuracy | Insectivory accuracy |
|---|---|---|---|---|---|---|---|
| Marsupial m2 | 52.6%<br>[*51.4%*] | ariaDNE + RFI | Primates | 32.0%<br>[*27.3%*] | 5/32<br>[*3/12*] | 25/53<br>[*5/14*] | 1/12<br>[*1/7*] |
| | 73.7%<br>[*73.0%*] | ariaDNE + RFI + TriTaHI | | 40.2%<br>[*36.4%*] | 5/32<br>[*0/12*] | 30/53<br>[*8/14*] | 4/12<br>[*4/7*] |
| | **84.2%**<br>**[*89.2%*]** | **ariaDNE + RFI + size** | | **64.9%**<br>[*69.7%*] | 25/32<br>[*10/12*] | 30/53<br>[*7/14*] | 8/12<br>[*6/7*] |
| | 94.7%<br>[*94.7*] | ariaDNE + RFI + size + TriTaHI | | 63.9%<br>[*63.6%*] | 25/32<br>[*7/12*] | 28/53<br>[*8/14*] | 9/12<br>[*6/7*] |
| Marsupial m3 | 69.4%<br>[*71.4%*] | ariaDNE + RFI | Primates | 33.0%<br>[*33.3%*] | 13/32<br>[*4/12*] | 10/53<br>[*1/14*] | 9/12<br>[*6/7*] |
| | 80.6%<br>[*80.0%*] | ariaDNE + RFI + TriTaHI | | 25.8%<br>[*33.3%*] | 15/32<br>[*5/12*] | 0/53<br>[*0/14*] | 10/12<br>[*6/7*] |
| | 88.9%<br>[*91.4%*] | **ariaDNE + RFI + size** | | **46.4%**<br>[*45.5%*] | 28/32<br>[*10/12*] | 10/53<br>[*2/24*] | 7/12<br>[*3/7*] |
| | 100%*<br>[*100%**] | ariaDNE + RFI + size + TriTaHI | | 82.2%*<br>[*73.7%**] | 27/32<br>[*8/12*] | NA | 10/12<br>[*6/7*] |
| Marsupial m2 & m3 | 64.9%<br>[*56.4%*] | ariaDNE + RFI | Primates | 33.0%<br>[*36.4%*] | 6/32<br>[*4/12*] | 24/53<br>[*4/14*] | 2/12<br>[*4/7*] |
| | 70.3%<br>[*79.5%*] | ariaDNE + RFI + TriTaHI | | 36.1%<br>[*36.4%*] | 15/32<br>[*4/12*] | 19/53<br>[*2/14*] | 1/12<br>[*6/7*] |
| | 90.5% | **ariaDNE + RFI + size** | | **69.1%** | 27/32 | 33/53 | 7/12 |

| | | | | | | | |
|---|---|---|---|---|---|---|---|
| | [*89.7%*] | | | [*69.7%*] | [*10/12*] | [*7/14*] | [*6/7*] |
| | 96.0% [*100%*] | ariaDNE + RFI + size + TriTaHI | | 63.9% [*48.5%*] | 25/32 [*9/12*] | 31/53 [*1/14*] | 6/12 [*6/7*] |

**Table S5.** QDA category-specific results with ariaDNE, RFI, size as metrics. Train sample = marsupial m2 sample (vertical categories), test sample = primate sample (horizontal categories). Bolded numbers show the results of **overlapping diets**.

| | **Folivory** | Frugivory | **Frugivory-Insectivory** | Hard-object feeding | **Insectivory** | Omnivory |
|---|---|---|---|---|---|---|
| Faunivory | 1 | 0 | 0 | 0 | 0 | 0 |
| **Folivory** | **10** | 7 | 1 | 9 | 0 | 0 |
| **Frugivory-Insectivory** | 1 | 2 | **7** | 0 | 1 | 1 |
| **Insectivory** | 0 | 0 | 6 | 0 | **6** | 0 |

**Table S6.** QDA category-specific results with ariaDNE, RFI, size as metrics. Train sample = marsupial m3 sample (vertical categories), test sample = primate sample (horizontal categories). Bolded numbers show the results of **overlapping diets**.

| | **Folivory** | Frugivory | **Frugivory-Insectivory** | Hard-object feeding | **Insectivory** | Omnivory |
|---|---|---|---|---|---|---|
| Faunivory | 1 | 0 | 1 | 0 | 0 | 0 |
| **Folivory** | **10** | 7 | 1 | 7 | 4 | 0 |
| **Frugivory-Insectivory** | 0 | 1 | **2** | 2 | 0 | 0 |
| **Insectivory** | 1 | 1 | 10 | 0 | **3** | 1 |

**Table S7.** QDA results (primates = training set, marsupials = test set) of specimen values vs. [*species averages*]. Training accuracy is calculated using the entire training sample (not leave-one-out). Training set of platyrrhine sample includes Folivory, Frugivory, Frugivory-Insectivory, Hard-object feeder, and the training set using the total primate sample included the same categories as well as the Insectivory category. Accuracy of the QDA test was calculated using shared categories only, i.e., Folivory, Frugivory, and Frugivory-Insectivory.

| Train sample (train accuracy) | Train accuracy | Variables | Test sample | Test accuracy | Folivory accuracy | Frugivory accuracy | Frugivory-Insectivory accuracy | Insectivory accuracy |
|---|---|---|---|---|---|---|---|---|
| Platyrrhine m2 | 73.8% | ariaDNE + RFI | Marsupials (m3 & m3) | 44.2% | 14/32 | 0/2 | 5/9 | NA |
| | 80% | **ariaDNE + RFI + TriTaHI** | | **48.8%** | 13/32 | 0/2 | 8/9 | NA |
| | 97.9% | ariaDNE + RFI + size | | 32.56% | 5/32 | 0/2 | 9/9 | NA |
| | 97.2% | ariaDNE + RFI + size + TriTaHI | | 32.56% | 5/32 | 0/2 | 9/9 | NA |
| Total primate sample | 66.1% [*72.5%*] | ariaDNE + RFI | Marsupials (m3 & m3) | 33.9% [*14.7%*] | 16/32 [*2/18*] | 0/2 [*0/1*] | 1/9 [*1/5*] | 5/22 [*2/10*] |
| | 71.8% [*70.6%*] | ariaDNE + RFI + TriTaHI | | 30.8% [*23.5%*] | 11/32 [*0/18*] | 0/2 [*0/1*] | 3/9 [*2/5*] | 6/22 [*6/10*] |
| | 93.7% [*92.2%*] | **ariaDNE + RFI + size** | | 50.8% **[*61.8%*]** | 21/32 [*14/18*] | 0/2 [*0/1*] | 5/9 [*5/5*] | 7/22 [*2/10*] |
| | 94.8% [*94.1%*] | ariaDNE + RFI + size + TriTaHI | | 32.3% [*29.4%*] | 14/32 [*7/18*] | 0/2 [*0/1*] | 5/9 [*2/5*] | 2/22 [*1/10*] |

Breakdown of QDA that includes **all primates** and **marsuspial m2 & m3s** (last column of table 7) per dietary category:

**RFI + ariaDNE (overall: 48.8%)**

| Faunivory | Folivory | Frugivory | Frugivory-Insectivory | Hard-object feeder | Insectivory | Omnivory |
|---|---|---|---|---|---|---|
| 0.7272727 | 0.1875000 | 0.8205128 | 0.3709677 | 0.8750000 | 0.6176471 | 0.5000000 |

**RFI + ariaDNE + size (overall: 64.8%)**

| Faunivory | Folivory | Frugivory | Frugivory-Insectivory | Hard-object feeder | Insectivory | Omnivory |
|---|---|---|---|---|---|---|
| 0.7272727 | 0.7187500 | 0.8717949 | 0.8064516 | 0.8750000 | 0.7352941 | 0.6666667 |

**RFI + ariaDNE + size + TriTaHI (overall: 69.2%)**

| Faunivory | Folivory | Frugivory | Frugivory-Insectivory | Hard-object feeder | Insectivory | Omnivory |
|---|---|---|---|---|---|---|
| 0.9090909 | 0.7812500 | 0.8717949 | 0.7419355 | 0.8750000 | 0.7647059 | 0.3333333 |

**RFI + ariaDNE + size + ariaDNE CV (overall: 75.8%)**

| Faunivory | Folivory | Frugivory | Frugivory-Insectivory | Hard-object feeder | Insectivory | Omnivory |
|---|---|---|---|---|---|---|
| 0.7272727 | 0.7500000 | 0.8974359 | 0.8225806 | 0.9500000 | 0.8529412 | 0.5000000 |

**RFI + ariaDNE + size + OPCR (overall: 79.1%)**

| Faunivory | Folivory | Frugivory | Frugivory-Insectivory | Hard-object feeder | Insectivory | Omnivory |
|---|---|---|---|---|---|---|
| 0.8181818 | 0.8593750 | 0.8974359 | 0.8387097 | 0.9250000 | 0.8823529 | 0.3333333 |

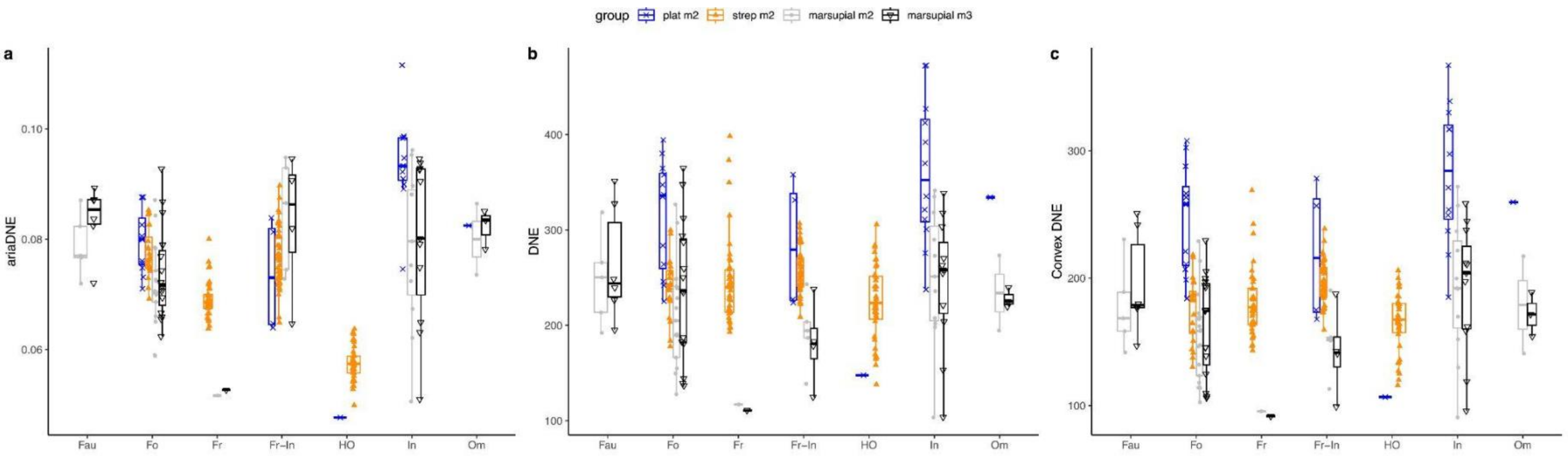

**Fig. S5** Sensitivity of curvature metrics to material. Plat m2 (blue) & strep m2 (orange) are µCT scans of plastic casts. Marsupial m2 (light grey) and marsupial m3 (black) are a mix of surface and µCT scans of biological specimens.

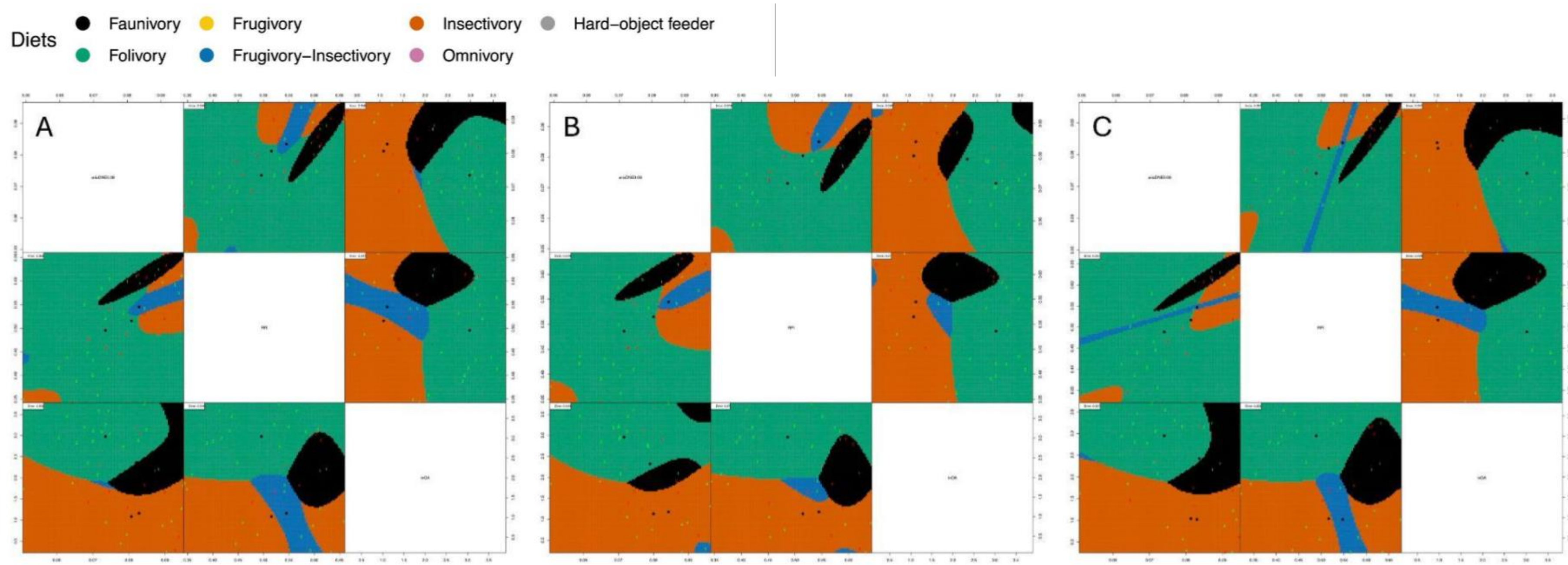

**Fig. S6** A. 2D-visualisation of QDA for marsupial (m2 & m3) species averages. B. marsupial m2-only. C. marsupial m3-only. 1 = faunivory, 2 = folivory, 3 = frugivory-insectivory, 4 = insectivory. Green number = correct, red number = incorrect in the training sample.

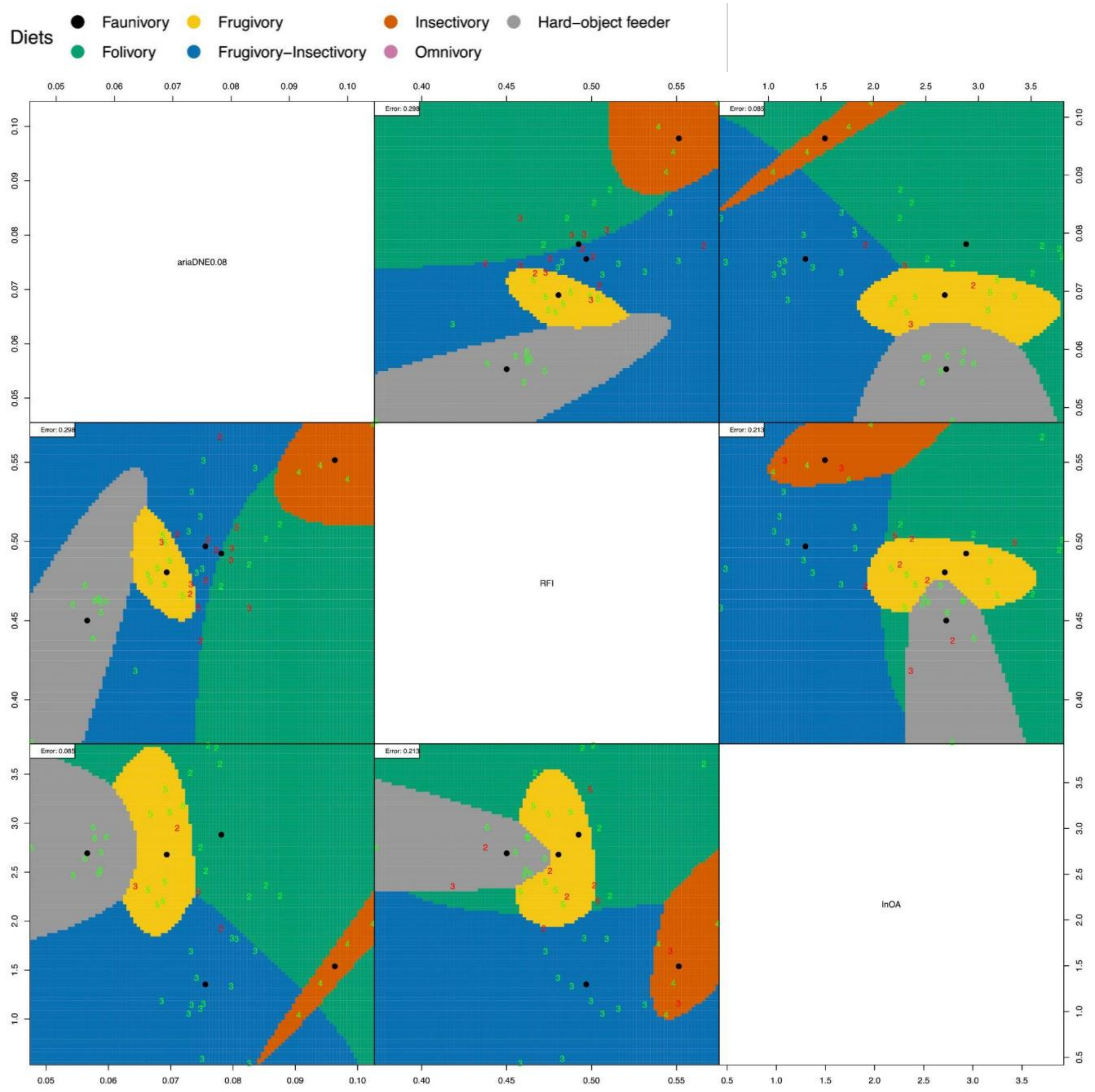

**Fig. S7** 2D-visualisation of QDA for primate-only species averages. 2 = folivory, 3 = frugivory-insectivory, 4 = insectivory, 5 = frugivory. Green number = correct, red number = incorrect in the training sample.

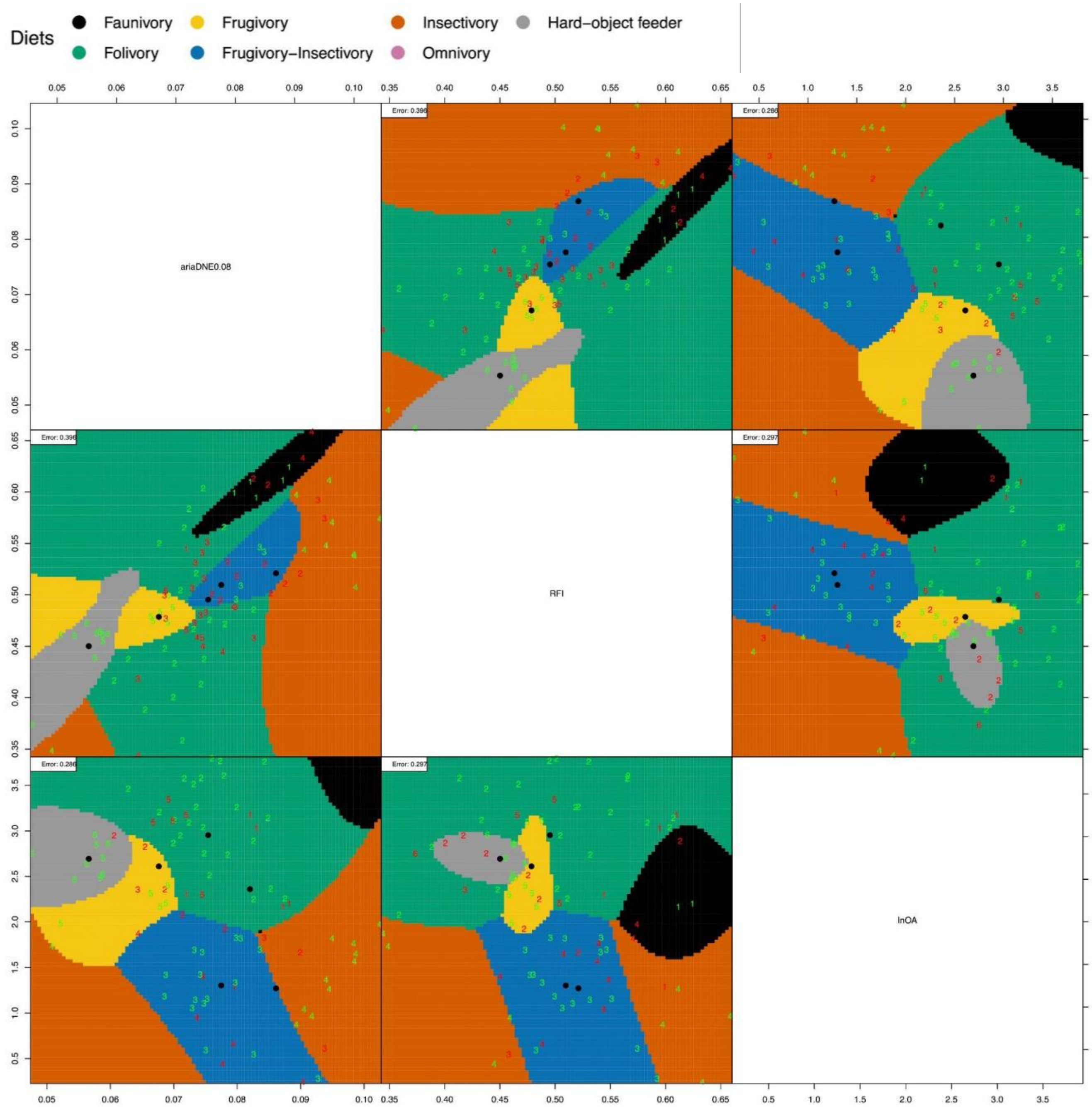

**Fig. S8** 2D-visualisation of QDA for total sample of marsupial (m2 & m3) and primate species averages. 1 = faunivory, 2 = folivory, 3 = frugivory-insectivory, 4 = insectivory, 5 = frugivory, 6 = hard-object feeder. Green number = correct, red number = incorrect in training sample.

Table S8. MorphoSource files information and funding acknowledgements.

| File | resolution | data uploaded by | Funding acknowledgement listed on MorphoSource | doi and ARK |
|---|---|---|---|---|
| Aeprymnus_rufescens_AMNH_22788 | 0.057922 | Arianna Harrington | NSF BCS 1825129 (to D.M. Boyer and A.R. Harrington) | doi:10.17602/M2/M76018 |
| Antechinus_flavipes_SAMA_M27013 | 0.026 | Jacob van Zoelen | CABAH | ark:/87602/m4/508065 |
| Caenolestes_fuliginosus_KU124015 | 0.014,0.014, 0.03 | Jessie Maisano | Ted Macrini provided access to these data , with data collection funded by NSF DEB-0309369 and data upload to MorphoSource funded by DBI-1902242. The files were downloaded from www.MorphoSource.org, Duke University. | ark:/87602/m4/M116130 |
| Caluromys_philander_AMNH95526 | 0.03711,0.037110, 0.079950 | Jessie Maisano | Ted Macrini provided access to these data , with data collection funded by NSF DEB-0309369 and data upload to MorphoSource funded by DBI-1902242. The files were downloaded from www.MorphoSource.org, Duke University. | ark:/87602/m4/M116113 |
| Caluromys_sp_DU_EA_162_m2 | 0.042338 | Arianna Harrington | NSF BCS 1552848 (to D M Boyer), NSF DBI 1458192 (to G F Gunnell) | doi:10.17602/M2/M58092 |
| Caluromysiops_irrupta_FMNH60698 | 0.089676 | Roger Benson | Roger Benson provided access to these data originally appearing in Marin-Serra A, Benson RBJ. 2020. Developmental constraints do not influence long-term phenotypic evolution of marsupial forelimbs as revealed by interspecific disparity and integration patterns. American Naturalist 195(3) , the collection of which was funded by the European Research Council (ERC) starting grant TEMPO (ERC-2015-STG-677774) to Roger Benson. The files were downloaded from www.MorphoSource.org, Duke University. | ark:/87602/m4/M82280 |
| Chironectes_minimus_AMNH129701 | 0.03223,0.032230, 0.035120 | Jessie Maisano | Jeri Rodgers provided access to these data , with data upload to MorphoSource funded by DBI-1902242. The files were downloaded from www.MorphoSource.org, Duke University. | ark:/87602/m4/M114558 |
| Dactylopsila_trivirgata_AMNH101984 | 0.017284 | Mary Silcox | Mary T. Silcox provided access to these data NSERC Discovery Grant to Mary T. Silcox. The files were downloaded from www.MorphoSource.org, Duke University. | ark:/87602/m4/M75882 |
| Dactylopsila_trivirgata_AMNH101985 | 0.016863 | Mary Silcox | Mary T. Silcox provided access to these data NSERC Discovery Grant to Mary T. Silcox. The files were downloaded from www.MorphoSource.org, Duke University. | ark:/87602/m4/M75886 |
| Dasycercus_blythi_SAMA_M3635 | 0.036 | Jacob van Zoelen | CABAH | ark:/87602/m4/508066 |
| Dasyurus_hallucatus_SAMA_M15386 | 0.054 | Jacob van Zoelen | CABAH | ark:/87602/m4/508067 |
| Dasyurus_maculatus_SAMA_M27408 | mandible = 4867150 polygons | Erin Mein | ARC Centre of Excellence for Australian Biodiversity and Heritage | ark:/87602/m4/537548 |
| Dendrolagus_lumholtzi_AMNH65254 | 0.068800,0.068800,0.160200 | Jessie Maisano | Ted Macrini provided access to these data , with data collection funded by NSF DEB-0309369 and data upload to MorphoSource funded by DBI-1902242. The files were downloaded from www.MorphoSource.org, Duke University. | ark:/87602/m4/M116135 |
| Didelphis_virginiana_TMM_M_2517 | 0.059600,0.059600,0.132000 | Jessie Maisano | Ted Macrini provided access to these data , with data collection funded by NSF DEB-0309369 and data upload to MorphoSource funded by DBI-1902242. The files were downloaded from www.MorphoSource.org, Duke University. | ark:/87602/m4/M116136 |
| Dorcopsis_muelleri_SAM_M13754 | mandible = 3337923 | Jorgo Ritsevski | CABAH | 10.17602/M2/M592786 |
| Dromiciops_gliroides_FMNH_127463 | 0.016100,0.016100,0.039500 | Jessie Maisano | Ted Macrini provided access to these data , with data collection funded by NSF DEB-0309369 and data upload to MorphoSource funded by DBI-1902242. The files were downloaded from www.MorphoSource.org, Duke University. | ark:/87602/m4/M116137 |
| Hypsiprymnodon_moschatus_AMNH184580 | 0.037210,0.037210,0.079920 | Barbara Sulbaran | Ted Macrini provided access to these data with data collection funded by NSF DEB-0309369 and data upload to MorphoSource funded by DBI-1902242. The files were downloaded from www.MorphoSource.org, Duke University. | ark:/87602/m4/M117097 |

| File | resolution | data uploaded by | Funding acknowledgement listed on MorphoSource | doi and ARK |
|---|---|---|---|---|
| Isoodon_obesulus_SAMA_M25975 | 0.056 | Diana Fusco | CABAH | ark:/87602/m4/536302 |
| Lestoros_inca_MVZ116049 | 0.017957 | Arianna Harrington | NSF BCS 1552848 (to D M Boyer), NSF DBI 1458192 (to G F Gunnell) | doi:10.17602/M2/M57778 |
| Macropus_eugenii_TMM_M1047 | 0.0623 | Barbara Sulbaran | Ted Macrini provided access to these data , with data collection funded by NSF DEB-0309369 and data upload to MorphoSource funded by DBI-1902242. The files were downloaded from www.MorphoSource.org, Duke University. | 10.17602/M2/M167398 |
| Marmosa_murina_NHMUK_1881 | 0.041269 | Roger Benson | Roger Benson provided access to these data originally appearing in Marin-Serra A, Benson RBJ. 2020. Developmental constraints do not influence long-term phenotypic evolution of marsupial forelimbs as revealed by interspecific disparity and integration patterns. American Naturalist 195(3) , the collection of which was funded by the European Research Council (ERC) starting grant TEMPO (ERC-2015-STG-677774) to Roger Benson. The files were downloaded from www.MorphoSource.org, Duke University. | ark:/87602/m4/M68271 |
| Monodelphis_domestica_TMMM_7599 | 0.045000,0.045000,0.090000 | Jessie Maisano | Tim Rowe and Ted Macrini provided access to these data , with data upload to MorphoSource funded by DBI-1902242. The files were downloaded from www.MorphoSource.org, Duke University. | 10.17602/M2/M168948 |
| Murexia_longicaudata_SAMA_M2816 | 0.35 | Diana Fusco | CABAH | ark:/87602/m4/557696 |
| Onychogale_frenta_UMZCa12_59_3 | 0.098246,0.098246,0.196491 | Roger Benson | Roger Benson provided access to these data originally appearing in Marin-Serra A, Benson RBJ. 2020. Developmental constraints do not influence long-term phenotypic evolution of marsupial forelimbs as revealed by interspecific disparity and integration patterns. American Naturalist 195(3) , the collection of which was funded by the European Research Council (ERC) starting grant TEMPO (ERC-2015-STG-677774) to Roger Benson. The files were downloaded from www.MorphoSource.org, Duke University. | ark:/87602/m4/M82448 |
| Perameles_gunnii_NTMU7600 | mandible = 645312 polygons | Diana Fusco | CABAH | ark:/87602/m4/538502 |
| Perameles_nasuta_MAGNT_U7608 | mandible = 69827 polygons | Diana Fusco | CABAH | ark:/87602/m4/546239 |
| Petauroides_volans_QM_J4643 | 0.036 | Jessica Ivory-Church | Australian Research Council Discovery Early Career Award DE120102034 | private file of Vera Weisbecker |
| Petaurs_australis_QM_J8598 | 0.036 | Jessica Ivory-Church | Australian Research Council Discovery Early Career Award DE120102035 | private file of Vera Weisbecker |
| Petrogale_penicillata_FMNH64435 | 0.068177,0.068177,0.136354 | Roger Benson | Roger Benson provided access to these data originally appearing in Marin-Serra A, Benson RBJ. 2020. Developmental constraints do not influence long-term phenotypic evolution of marsupial forelimbs as revealed by interspecific disparity and integration patterns. American Naturalist 195(3) , the collection of which was funded by the European Research Council (ERC) starting grant TEMPO (ERC-2015-STG-677774) to Roger Benson. The files were downloaded from www.MorphoSource.org, Duke University. | ark:/87602/m4/M82308 |
| Petropseudes_dahli_AMNH183391 | 0.044920,0.044920,0.048660 | Jessie Maisano | eri Rodgers provided access to these data , with data upload to MorphoSource funded by DBI-1902242. The files were downloaded from www.MorphoSource.org, Duke University. | ark:/87602/m4/M114566 |
| Phalanger_carmelitae_SAMA_M2901 | 0.043 | Diana Fusco | CABAH | ark:/87602/m4/535483 |
| Phalanger_orientalis_AMNH157211 | 0.054000,0.054000,0.121 | Barbara Sulbaran | Ted Macrini provided access to these data , with data collection funded by NSF DEB-0309369 and data upload to MorphoSource funded by DBI-1902242. The files were downloaded from www.MorphoSource.org, Duke University. | ark:/87602/m4/M167380 |
| Phascogale_tapoatafa_SAMA_M3824 | 1072904 | Diana Fusco | CABAH | ark:/87602/m4/M167380 |

| File | resolution | data uploaded by | Funding acknowledgement listed on MorphoSource | doi and ARK |
|---|---|---|---|---|
| Phascolarctos_cinereus_AMNH65608 | 0.08338 | Arianna Harrington | NSF BCS 1825129 (to D.M. Boyer and A.R. Harrington) | doi:10.17602/M2/M99914 |
| Philander_opossum_QM_J3461 | 0.053 | Vera Weisbecker | Australian Research Council DE120102034; DP170103227 | private file of Vera Weisbecker |
| Pseudochirulus_forbesi_AMNH104136 | 0.03223,0.032230,0.035470 | Jessie Maisano | Jeri Rodgers provided access to these data , with data upload to MorphoSource funded by DBI-1902242. The files were downloaded from www.MorphoSource.org, Duke University. | ark:/87602/m4/M114568 |
| Rhyncholestes_raphanurus_MVZ163773 | 0.018392 | Arianna Harrington | NSF BCS 1552848 (to D M Boyer), NSF DBI 1458192 (to G F Gunnell) | doi:10.17602/M2/M57244 |
| Smithopsis_crassicaudata_SAMA_M12809 | 0.023 | Jacob van Zoelen | CABAH | ark:/87602/m4/508327 |
| Spilocuscus_maculatus_FMNH31752 | 0.075973,0.075973,0.151946 | Roger Benson | Roger Benson provided access to these data, the collection of which was funded by the European Research Council (ERC) starting grant TEMPO (ERC-2015-STG-677774) to Roger Benson. The files were downloaded from www.MorphoSource.org, Duke University. | ark:/87602/m4/M164405 |
| Thylogale_billardieri_SAMAM2868 | mandible = 1215244 polygons | Jacob van Zoelen | CABAH | ark:/87602/m4/568545 |
| Thylogale_stigmatica_MAGNT_U8203 | mandible = 2039374 | Diana Fusco | CABAH | ark:/87602/m4/538617 |
| Trichosurus_arnhemensis_MAGNT_U5993 | mandible = 746401 polygons | Diana Fusco | CABAH | ark:/87602/m4/544616 |
| Trichosurus_caninus_MAGNT_U7946 | mandible = 1809823 polygons | Diana Fusco | CABAH | 10.17602/M2/M546257 |
| Trichosurus_vulpecula_TMM849 | 0.053000,0.053000,0.116000 | Barbara Sulbaran | Ted Macrini provided access to these data , with data collection funded by NSF DEB-0309369 and data upload to MorphoSource funded by DBI-1902242. The files were downloaded from www.MorphoSource.org, Duke University. | ark:/87602/m4/M167392 |

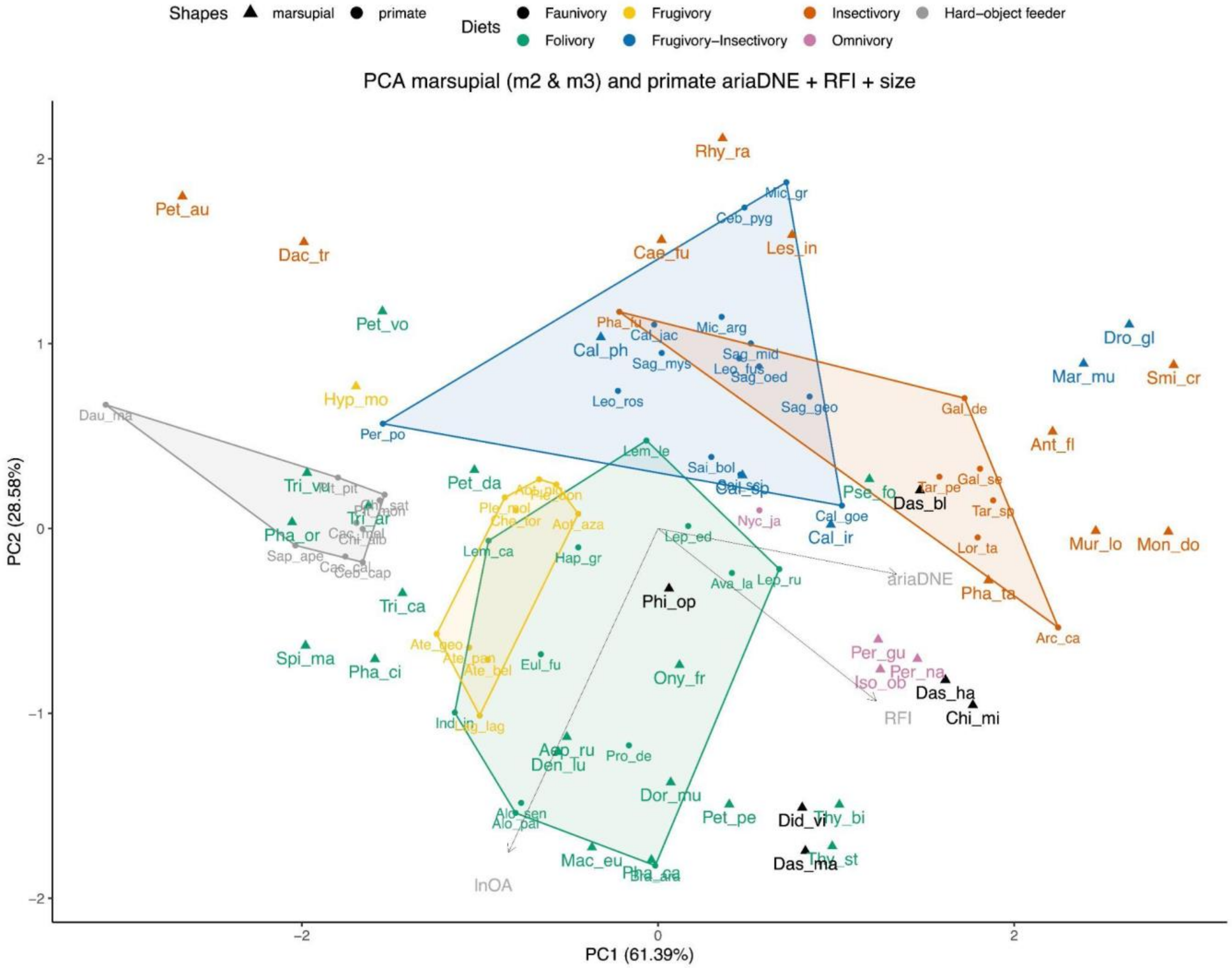

**Fig. S9** Principal Component Analysis of ariaDNE, RFI, and lnOA plot showing PC1 and PC2 scores for total marsupial (m2 & m3) and primate species averaged data, capturing 89.97% of the variation. Note that the PC loading arrows are not to scale. Convex hulls are primate-only. Marsupial taxon abbreviations can be found in Table 3.

**Dietary category justifications**

From de Vries et al. (2024 table 3):

Exudate feeders: Fruit = 0.28, Animal matter = 0.21, Exudates = 0.52

Hard-object feeders ('seed eaters' in original source): Seeds >0.32, Fruit = 0.19-0.47, Leaves <0.1, Animal matter < 0.1

Folivore: Leaves >0.5, Fruit 0.4-0.46

Frugivore: Fruit > 0.58, Leaves 0.12-0.28, Animal matter <0.13

Frugivore-insectivore: Fruit >0.56, Leaves <0.06, Animal matter >0.13

Additional categories:

Insectivore: >50% insects, less fruit consumption than in the frugivore-insectivore category

Faunivore: regular consumption of vertebrate prey

Omnivore: animals, fruit, and non-reproductive parts of plants (e.g. leaves, roots) all form major parts of the diet

### Primates

*Avahi laniger*: folivore (Harcourt 1991; Faulkner and Lehman 2006).

*Galagoides demidoff*: insectivore - diet is primarily insects: 70% insects, 19% fruit, 10% gums, and a very small amount of leaves and buds (Charles-Dominique 1977; Hladik 1979).

*Galago senegalensis*: insectivore - primary diet is insects and acacia gum, so if exudates are excluded, then, it's insectivore (Harcourt 1986).

*Lemur catta*: folivore - diet can be >50% leaves at certain times of the year (Simmen et al. 2003; Gould et al. 2011:2, 3).

*Indri indri*: folivore (Britt et al. 2002; Powzyk and Mowry 2003, 2006; Randrianarison et al. 2022).

*Lepilemur edwardsi*: folivore (Thalmann 2001).

*Lepilemur leucopus*: folivore (Dröscher and Kappeler 2014).

*Lepilemur ruficaudatus*: folivore (Ganzhorn et al. 2004).

*Microcebus griseorufus*: frugivore-insectivore (Crowley et al. 2014:2) - eats considerable amounts of exudates, but if excluded then diet is predominately fruit and insects.

*Phaner furcifer*: insectivore - diet consists nearly entirely of tree exudates and insect secretions (Petter et al. 1971) and its exudativorous diet is complemented with insects (Charles-Dominique and Petter 1980; only about 10%, Hladik et al. 1980).

*Propithecus deckenii*: folivore (55%, Fabrice et al. 2024).

*Tarsius pelengensis*: insectivore (Syahrullah et al. 2023).

*Hapalemur griseus*: folivore, >80% bamboo or other leafy materials (Grassi 2006).

*Arctocebus calabarensis*: insectivore (Charles-Dominique 1977).

*Daubentonia madagascariensis*: hard-object feeder based on >25% of extremely hard shelled *Canarium* seeds in their diet (Sterling 1994; Randimbiharinirina et al. 2018), see main text for full justification

*Eulemur fulvus*: folivore (Sussman 1977; Ganzhorn 1986; Overdorff 1993).

*Loris tardigradus*: insectivore (Nekaris and Jayewardene 2003).

*Nycticebus javanicus*: exudativorous (gums and nectar) according to Cabana et al. (2017, figure 1), but when exudates are excluded, its diet consists mostly of insects and leaves, with a lower contribution of flowers and fruits (based on % time feeding), and we therefore group it as an omnivore.

*Perodicticus potto*: frugivorous with a considerable contribution of gums and insects to its diet (Nekaris and Bearder 2007). As we exclude the exudativorous component, we classify it as a frugivore-insectivore.

### Marsupials

*Caenolestes fuliginosus*: highly insectivorous based on gut contents, with lepidopteran larvae,

centipedes, unidentified arachnids making up 75% of gut contents of eight specimens, both in volume and in frequency of being present in each specimen (Barkley and Whitaker 1984).
*Lestoros inca*: insectivore very little published on its behaviour and diet. We classified this as an insectivore following (Siciliano Martina 2013), based on general Canaestolidae behaviour.
*Rhyncholestes raphanurus*: insectivore as it mainly consumed invertebrates such as arthropods and annelids (54.7%), with its secondary dietary component being made of by plant material and fungi (39%, Meserve et al. 1988).
*Caluromys philander*: frugivore-insectivore. Classified as frugivore by Julien-Laferrière (1999), with the majority of its diet being fruits. But an estimate of 25% of insects in its diet is reported (Atramentowicz 1982; Julien-Laferrière 1999), therefore being an Frugivore-insectivore in our dietary scheme.
*Caluromys* sp: frugivore-insectivore, following our classification of *Caluromys philander.*
*Caluromysiops irrupta*: frugivore-insectivore. Frugivore-omnivore according to Robinson and Redford (1986), which is defined as >50% fruits, remainder mostly invertebrates and vertebrates.
*Chironectes minimus*: faunivore, its diet consists mainly of crustaceans (Medellín 1991) and fishes (Mondolfi and Padilla 1958; Galliez et al. 2009).
*Didelphis virginiana*: faunivore. Its diet is described as being highly opportunistic, with the bulk being made up of animal foods, mostly vertebrate prey and insects (McManus 1974).
*Marmosa murina*: frugivore-insectivore. A reported 80% of studied gut contents had arthropods present, compared to 100% of gut specimens containing fruit (Parreira Claro and Hannibal 2022).
*Monodelphis domestica*: insectivore. Wild populations focus on predating on invertebrates, whereas captive members also accept vertebrates into their diet in the laboratory (Streilein 1982).
*Philander opossum*: faunivore. They feed on small mammals, birds, eggs, carrion, and fruit (Hunsaker 2012).
*Dactylopsila trivirgata*: insectivore, mostly feeding on invertebrates (Rawlins and Handasyde 2002).
*Petaurus australis*: insectivore. The bulk of its diet is exudates, but this is supplemented by a variety of arthropods (Smith and Russell 1982; Harcourt 1986; Schülke 2003 fig. 2).
*Aepyprymnus rufescens*: grasses, herbaceous plants, roots, and fungi (Baker and Gynther 2023).
*Dendrolagus lumholtzi*: folivore as its diet is mainly rainforest foliage (Martin 2005).
*Macropus eugenii*: folivore as its diet mainly consists of grasses and foliage (Williamson 1986).
*Onychogalea frenata*: folivore, as its diet consists mainly of herbaceous plants (Dawson et al. 1992); 40-50% graze and the rest browse (Tierney 1985).
*Petrogale penicillata*: folivore, as its diet consisted of 10–40% grass, 30–50% browse, 12–45% forbs and minor quantities of orchid/lilies and sedges (Tuft et al. 2011).
*Dromiciops gliroides*: insectivore-frugivore
*Phascolarctos cinereus*: specialised folivore of *Eucalyptus* (Marsh et al. 2021; Eisenhofer et al. 2023)
*Petauroides volans*: folivore, with its diet nearly exclusively consisting of young leaves and flower buds of a few *Eucalyptus* species (Harris and Maloney 2010).
*Petropseudes dahli*: folivore as the primary component of its diet is leaves, although it occasionally consumes flowers and fruit as well (Runcie 2002).
*Pseudochirulus forbesi*: folivore (Hume et al. 1993; Stephens et al. 2006)
*Phalanger orientalis*: folivore, with their diet consisting of leaves, fruit, and bark (Farida 2022).
*Spilocuscus maculatus*: folivore with foliage, fruits, and shoots being the most common parts of its diet. It is described as being principally folivorous and partially frugivorous by Heinsohn (2002), or as feeding primarily on fruits and leaves (Latinis 1996). However, according to Saragih et al. (2010): "Approximately 64.4% of cuscus diets is the combination between pulp of fruit and epidermis of fruit, while 21.1% is shoot". As it is inconclusive whether the largest component of the diet of *S. maculatus* is fruits or leaves, we assign this taxon to being a folivore, in which the proportions of leaves to fruits are closest to 50:50 (Leaves >0.5, Fruit 0.4-0.46, see above), in contrast to our definition of frugivores for which fruits are at least double as abundant in the diet as leaves are (Fruit > 0.58, Leaves < 0.28).

*Trichosurus vulpecula*: folivore. Highly variable diet, but leaves comprising the majority of its diet (Wilson and Mittermeier 2015).
